# Rapid global phaseout of animal agriculture has the potential to stabilize greenhouse gas levels for 30 years and offset 68 percent of CO_2_ emissions this century

**DOI:** 10.1101/2021.04.15.440019

**Authors:** Michael B. Eisen, Patrick O. Brown

## Abstract

Animal agriculture contributes significantly to global warming through ongoing emissions of the potent greenhouse gases methane and nitrous oxide, and displacement of biomass carbon on the land used to support livestock. However, because estimates of the magnitude of the effect of ending animal agriculture often focus on only one factor, the full potential benefit of a more radical change remains underappreciated. Here we quantify the full “climate opportunity cost” of current global livestock production, by modeling the combined, long-term effects of emission reductions and biomass recovery that would be unlocked by a phaseout of animal agriculture. We show that, even in the absence of any other emission reductions, persistent drops in atmospheric methane and nitrous oxide levels, and slower carbon dioxide accumulation, following a phaseout of livestock production would, through the end of the century, have the same cumulative effect on the warming potential of the atmosphere as a 25 gigaton per year reduction in anthropogenic CO_2_ emissions, providing half of the net emission reductions necessary to limit warming to 2°C. The magnitude and rapidity of these potential effects should place the reduction or elimination of animal agriculture at the forefront of strategies for averting disastrous climate change.

**Significance Statement:** The use of animals to produce food has a negative impact on the climate, but the benefits of a global switch to a plant based diet are underappreciated. We show that the global warming impact, through the rest of this century, of eliminating greenhouse gas emissions from livestock and allowing native ecosystems to regrow on the land currently used to house and feed livestock, would be equivalent to a 68% reduction in carbon dioxide emissions. We hope putting clearer numbers on the “climate opportunity cost” of our continued use of animals as food technology will help policymakers and the public properly prioritize dietary change as a climate defense strategy.

**Declaration of Conflict of Interest:** Patrick Brown is the founder and CEO of Impossible Foods, a company developing alternatives to animals in food-production. Michael Eisen is an advisor to Impossible Foods. Both are shareholders in the company and thus stand to benefit financially from reduction of animal agriculture.

## Introduction

The use of animals as a food-production technology has well-recognized negative impacts on our climate. The historical reduction in terrestrial biomass as native ecosystems were transformed to support grazing livestock and the cultivation of feed and forage crops accounts for as much as a third of all anthropogenic CO_2_ emissions to date (Friedlingstein et al., 2020; Hayek et al., 2021). Livestock, especially large ruminants, and their supply chains, also contribute significantly to anthropogenic emissions of the potent greenhouse gases (GHGs) methane and nitrous oxide (Gerber et al., 2013; MacLeod et al., 2018; Steinfeld et al., 2006).

Solving the climate crisis requires massive cuts to GHG emissions from transportation and energy production. But even in the context of large-scale reduction in emissions from other sources, major cuts in food-linked emissions are likely necessary by 2075 to limit global warming to 1.5°C (Clark et al., 2020). While a reduction of food-linked emissions can likely be achieved by increasing agricultural efficiency, reducing food waste, limiting excess consumption, increasing yields, and reducing the emission intensity of livestock production (Cusack et al., 2021; Hristov et al., 2013a, 2013b; Montes et al., 2013; Poore and Nemecek, 2018; Springmann et al., 2018a), they are not anticipated to have the same impact as a global transition to a plant-rich diet (Clark et al., 2020; Gerber et al., 2013).

Nutritionally balanced plant-dominated diets are common, healthy and diverse (Agnoli et al., 2017; American Dietetic Association and Dietitians of Canada, 2003; Craig et al., 2009; Tilman and Clark, 2014; Willett et al., 2019), but are rarely considered in comprehensive strategies to mitigate climate change (IPCC, 2018), and there is controversy about their viability and the magnitude of their climate benefit (Liu et al., 2021). One source of this discordance is that widely cited estimates of livestock contributions to global warming (Gerber et al., 2013; Steinfeld et al., 2006; Twine, 2021) account only for ongoing emissions, and not for the substantial and reversible warming impact of historical land use change (Hayek et al., 2021; Strassburg et al., 2020).

The Food and Agriculture Organization (FAO) of the United Nations estimates that emissions from animal agriculture represent around 7.1 Gt CO_2_eq per year (Gerber et al., 2013), 14.5% of annual anthropogenic greenhouse gas emissions, although this is based on outdated data and likely now represents and underestimate (Twine, 2021), and recent estimates (Hayek et al., 2021) suggest that on the order of 800 Gt CO_2_ equivalent carbon could be fixed via photosynthesis if native biomass were allowed to recover on the 30% of Earth’s land surface current devoted to livestock production. Thus, crudely, eliminating animal agriculture has the potential to reduce net emissions by the equivalent of around 1,350 Gt CO_2_ this century. To put this number in perspective, total anthropogenic CO_2_ emissions since industrialization are estimated to be around 1,650 Gt (Friedlingstein et al., 2020).

However, a substantial fraction of the emissions impact of animal agriculture comes from methane (CH_4_) and nitrous oxide (N_2_O), which decay far more rapidly than CO_2_ (the half-lives of CH_4_ and N_2_O are around 9 and 115 years, respectively), and recent studies have highlighted the need to consider these atmospheric dynamics when assessing their impact (Allen et al., 2018, 2016; Cain et al., 2019). Of critical importance, many of the beneficial effects on greenhouse gas levels of eliminating livestock would accrue rapidly, via biomass recovery and decay of short-lived atmospheric CH_4_, and their cooling influence would be felt for an extended period of time.

Our goal here was to accurately quantify the full impact of current animal agriculture on the climate, taking into account the currently unrealized opportunities for emission reduction and biomass recovery together, and explicitly considering the impact of their kinetics on warming. Our approach differs from other recent studies (Springmann et al., 2018b; Xu et al., 2021) in that we did not attempt to predict how global food production and consumption might change with growing populations, economic development, advances in agriculture, climate change and other socioeconomic factors. Nor do we tackle the social, economic, nutrition and agricultural challenges inherent to such a large change in global production.

We used publicly available, systematic data on livestock production in 2019 (FAO, 2021), livestock-linked emissions (FAO, 2021; MacLeod et al., 2018), and biomass recovery potential on land currently used to support livestock (Hayek et al., 2021) to predict how the phaseout of all or parts of global animal agriculture production would alter net anthropogenic emissions. We then used a simple climate model to project how these changes would impact the evolution of atmospheric GHG levels and warming for the rest of the century.

We calculated the combined impact of reduced emissions and biomass recovery by comparing the cumulative reduction, relative to current emission levels, of the global warming potential of GHGs in the atmosphere for the remainder of the 21st century under different livestock replacement scenarios to those that would be achieved by constant annual reductions in CO_2_ emissions.

## Results

### Modeling the effect of eliminating animal agriculture on GHG levels

We implemented a simple climate model that projects atmospheric GHG levels from 2020 to 2100 based on a time series of annual emissions of CO_2_, CH_4_ and N_2_O and a limited set of parameters. We then compared various hypothetical dietary perturbations to a “business as usual” (BAU) reference in which emissions remain fixed at 2019 levels, based on global emissions data from FAOSTAT (FAO, 2021).

The dietary scenarios include the immediate replacement of all animal agriculture with a plant-only diet (IMM-POD), a more gradual transition, over a period of 15 years, to a plant-only diet (PHASE-POD), and versions of each where only specific animal products were replaced.

We updated estimates of global emissions from animal agriculture using country-, species- and product-specific emission intensities from the Global Livestock Environmental Assessment Model (MacLeod et al., 2018), and country-specific data on primary production of livestock products from the Food and Agriculture Organization (FAO) database FAOSTAT (FAO, 2021).

Based on this analysis, in 2019 (the most recent year for which full data are available), global production of animal-derived foods led to direct emissions of 1.6 Gt CO_2_, due primarily to energy use (as our model assumes constant overall rates of consumption, we excluded emissions due to land clearing, which are associated with agricultural expansion), 120 Mt CH_4_ due primarily to enteric fermentation and manure management, and 7.0 Mt N_2_O due primarily to fertilization of feed crops and manure management (Figure 1 and Figure 1-S1).

**Figure 1.**
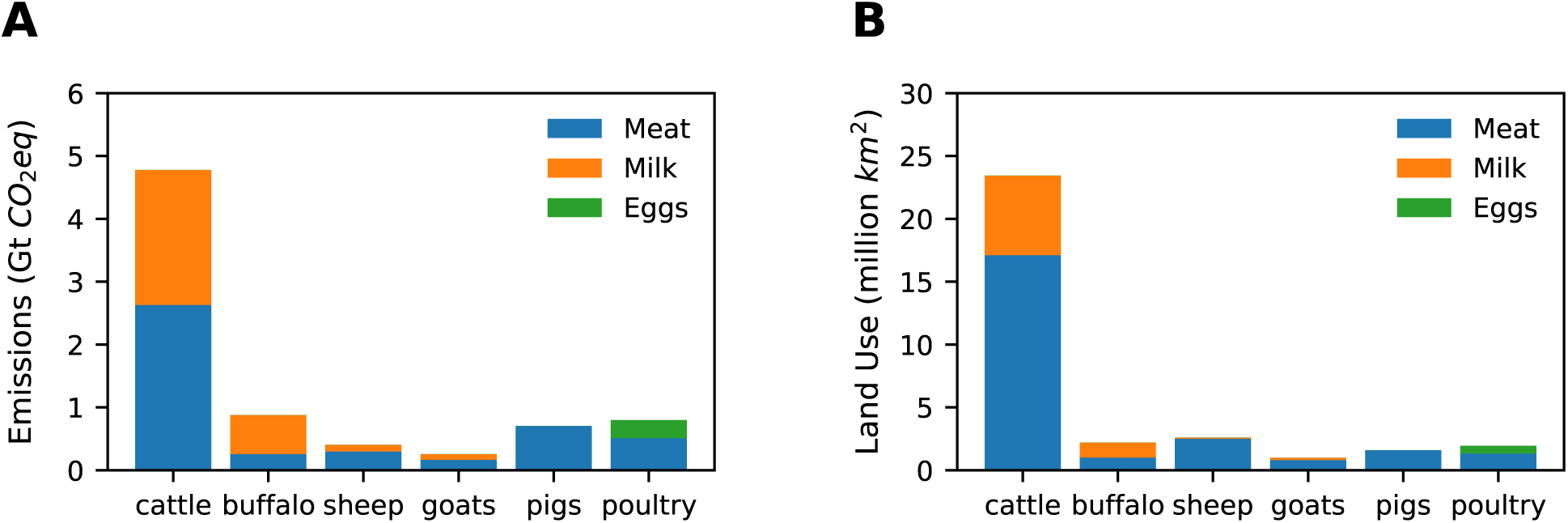
Global emissions and land use footprints of animal agriculture. Total *CO*_2_ equivalent emissions (A) assembled from species, product and country-specific production data from FAOSTAT for 2019 and species, product, region and greenhouse-gas specific emissions data from GLEAM (MacLeod et al., 2018), using *CO*_2_ equivalents of 34 for *CH*_4_ and 298 for *N*_2_*O*. Land use (B) assembled from species, product and country-specific production data from FAOSTAT for 2019 and species and product specific land use data from (Poore and Nemecek, 2018).

These numbers are broadly consistent with other recent estimates (Gerber et al., 2013; Steinfeld et al., 2006; Xu et al., 2021), and correspond, respectively, to 4% of CO_2_, 35% of CH_4_ and 66% of N_2_O emissions from all human activities, using total human emissions data from FAOSTAT (FAO, 2021). Combining the effects of the three gases, using global warming potentials from (Intergovernmental Panel on Climate Change, 2014), results in 6.3 Gt CO_2_eq, with the major difference from the 7.1 Gt CO_2_eq number cited above coming from our exclusion of ongoing land use change.

We modeled the recovery of biomass on land currently used in livestock production using data from (Hayek et al., 2021) who estimate that the return of land currently used in livestock production to its native state would sequester, over 30 years, 215.5 Gt of carbon (equivalent to 790 Gt of CO_2_) in plant and non-living biomass. A similar estimate was obtained by (Strassburg et al., 2020).

We assumed in all these hypothetical scenarios that non-agricultural emissions would remain constant; that food from livestock is replaced by a diverse plant based diet; and that, when land is removed from livestock production, the conversion of atmospheric CO_2_ into terrestrial biomass occurs linearly over the subsequent thirty years. (We consider alternative assumptions in the “Sensitivity Analysis” section below).

We emphasize that we are not predicting what will happen to global diets. Rather we are projecting simplified scenarios of dietary change forward through time to characterize and quantify the climate impact of current animal agriculture production. Our climate model is intentionally simple, considering only the partition of terrestrial emissions into the atmosphere, and the decay of methane and nitrous oxide, although it replicates the qualitative behavior of widely used MAGICC6 (Meinshausen et al., 2011).

Figure 2 shows annual emissions and projected atmospheric levels of CO_2_, CH_4_ and N_2_O under BAU and PHASE-POD through the end of the century (projections for IMM-POD and additional scenarios are shown in the supplemental versions of Figure 2).

**Figure 2.**
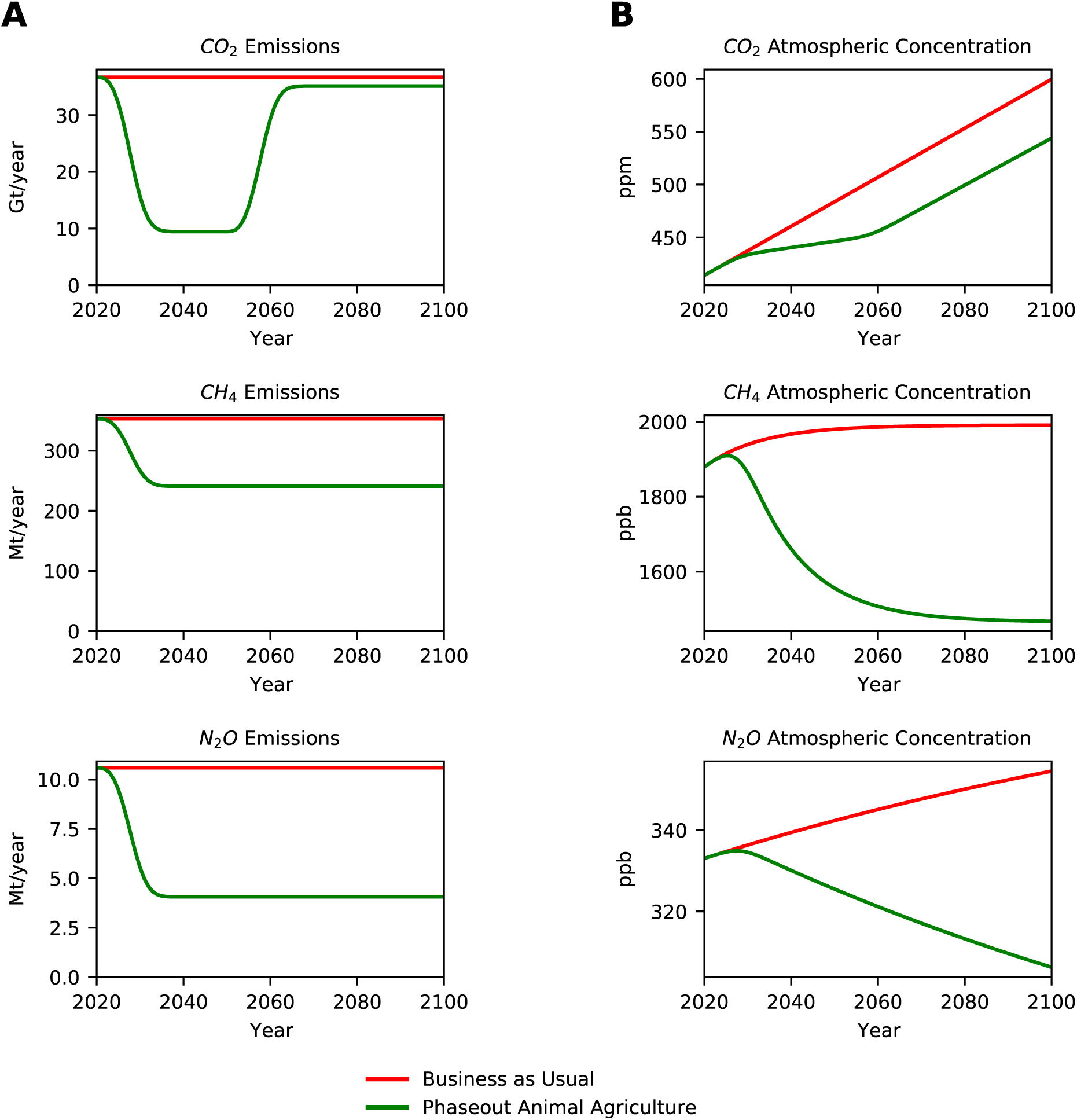
Impact of 15 year phaseout of animal agriculture on atmospheric greenhouse gas levels. (A) Projected annual emissions of *CO*_2_, *CH*_4_ and *N*_2_*O* for Business as Usual (red) and PHASE-POD (green) assuming a 15 year transition to new diet and 30 year carbon recovery. (B) Projected atmospheric concentrations of *CO*_2_, *CH*_4_ and *N*_2_*O* under each emission scenario.

### Rapid phaseout of animal agriculture would freeze increases in the warming potential of the atmosphere for 30 years

The impact of PHASE-POD on CO_2_ emissions would be greatest in the period between 2030 and 2060, when biomass recovery on land previously occupied by livestock or feed crops reaches its peak, slowing the rise of atmospheric CO_2_ levels during this interval.

Atmospheric CH_4_ and N_2_O levels continue to increase in both BAU and PHASE-POD during the transition period, but begin to drop in PHASE-POD as the abatement of animal agriculture-linked emissions accelerates. CH_4_, with a half-life in the atmosphere of around 9 years, approaches a new and lower steady-state level towards the end of the century, while N_2_O, with a half-life of around 115 years, does so over a longer time-scale.

To capture the combined global warming impact of the changing levels of these GHGs, we calculated radiative forcing (RF), the reduction in radiative cooling by GHG absorption of infrared radiation, using the formulae described in (Myhre et al., 1998; Shine, 2000) and used in MAGICC6 (Meinshausen et al., 2011).

Figure 3 shows that with PHASE-POD there would effectively be no net increase in RF between 2030 and 2060. And even after that 30-year pause in the previously monotonically increasing global warming potential of the atmosphere, the difference in RF between the POD and BAU scenarios would continue to increase, due to the absence of direct emissions from animal agriculture and the continuing decay of previously emitted CH_4_ and N_2_O towards lower steady-state values.

**Figure 3.**
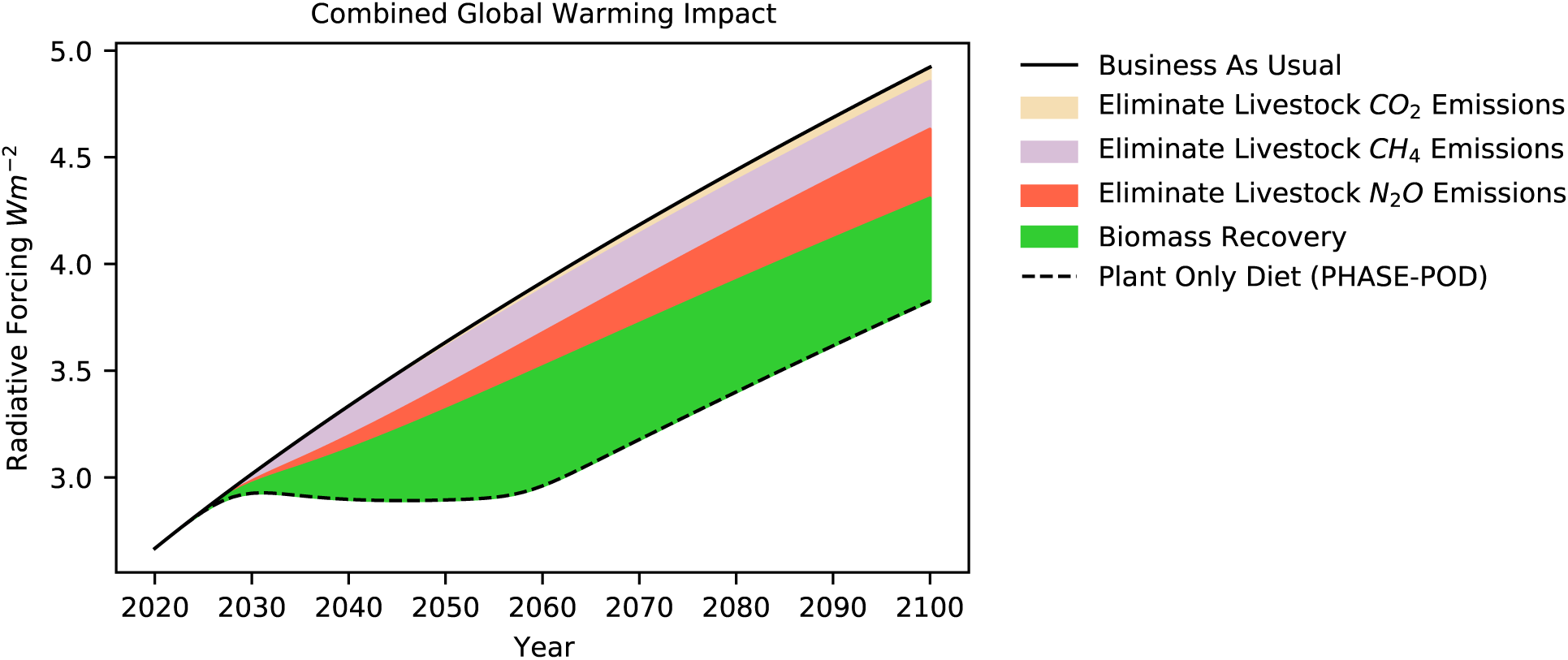
Phaseout of animal agriculture reduces global warming impact of atmosphere. Effect of eliminating emissions linked to animal agriculture and of biomass recovery on land currently used in animal agriculture on Radiative Forcing (RF), a measure of the instantaneous warming potential of the atmosphere. RF values computed from atmospheric concentrations in by formula of (Myhre et al., 1998; Ramaswamy et al., 2001) as modified in MAGICC6 (Meinshausen et al., 2011) with adjustment for gasses other than *CO*_2_, *CH*_4_ and *N*_2_*O* as described in text.

### Rapid phaseout of animal agriculture could achieve half of the emission reductions needed to meet Paris Agreement GHG targets

By the end of the century the RF under PHASE-POD would be 3.8 Wm^-2^ compared to 4.9 Wm^-2^ for BAU, a reduction in RF equivalent to what would be achieved by eliminating 1,680 Gt of CO_2_ emissions (Figure 4-S1), or 46 years of global anthropogenic CO_2_ emissions at the current rate of 36 Gt/year.

In 2010, the climate modeling community defined a series of four “Representative Concentration Pathways” that capture a wide range of future warming scenarios, leading to 2100 RF levels of 8.5, 6.0, 4.5 and 2.6 Wm^-2^ (which is approximately the RF of current atmospheric greenhouse gas levels), respectively (Moss et al., 2010; van Vuuren et al., 2011). These model pathways were extended after the Paris Agreement to include a target of 1.9 Wm^-2^. Although the exact relationship between RF and global warming is incompletely understood, 2100 RF values of 1.9 and 2.6 Wm^-2^ are generally used as targets for limiting warming in this century to 1.5°C and 2.0°C, respectively, over the baseline pre-industrial global average temperature (IPCC, 2018).

Reducing 2100 RF from 4.9 Wm^-2^ under BAU to 2.6 Wm^-2^ would require a reduction of atmospheric CO_2_ levels by 204 ppm, equivalent to 3,230 Gt of CO_2_ emissions (Figure 4 and Figure 4-S1), and an additional 47 ppm reduction, equivalent to 750 Gt of CO_2_ emissions, would be required to reach 1.9 Wm^-2^.

**Figure 4.**
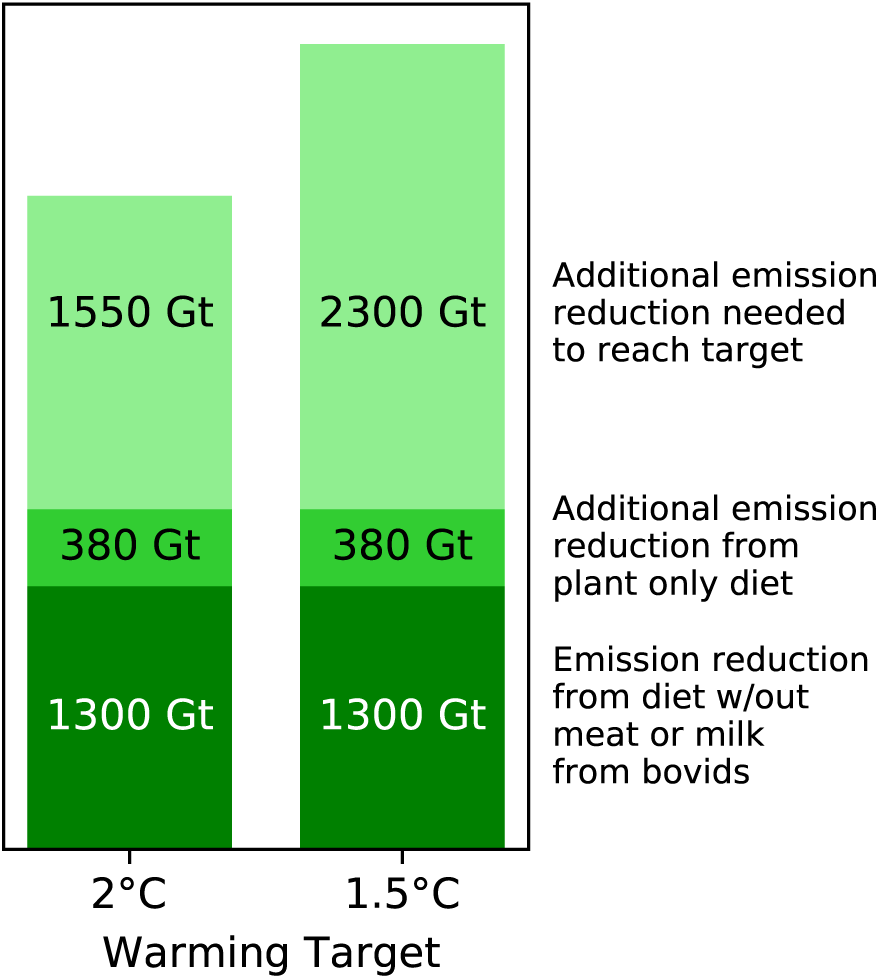
Impact of dietary transitions in curtailing global warming. Using projected *CH*_4_ and *N*_2_*O* levels in 2100 under business as usual diet as a baseline for RF calculation, we computed the *CO*_2_ reductions necessary to reduce RF from the business as usual diet level of RF=1.31 to the bovid-free diet level of RF=4.09 (1300 Gt *CO*_2_), the plant-only diet level of RF=3.83 (1680 Gt *CO*_2_), the 2.0 C global warming target of RF=2.6 (3230 Gt *CO*_2_) and the 1.5 C global warming target of RF=1.9 (3980 Gt *CO*_2_). For this analysis we used a corrected RF that accounts for the absence of other gases in our calculation by training a linear regression model on published MAGICC6 output to estimate from *CO*_2_, *CH*_4_ and *N*_2_*O* levels the residual RF impact of other gases.

Thus the 1,680 Gt of CO_2_ equivalent emissions reductions from the phased elimination of animal agriculture, would, without any other intervention to reduce GHG emissions, achieve 52% of the net GHG emissions reductions necessary to reach the 2100 RF target of 2.6 Wm^-2^ and 42% of the emissions reductions necessary to reach the 1.9 Wm^-2^ target (IPCC, 2018).

### Eliminating animal agriculture has the potential to offset 68 percent of current anthropogenic CO_2_ emissions

While widely used, such single point estimates of radiative forcing tell an incomplete story, as temperature change, and other climate impacts, depend cumulatively on the temporal trajectories of changing atmospheric greenhouse gas levels.

To capture such dynamic effects, we computed, for each dietary scenario, the integral with respect to time of the RF difference between the scenario and BAU, from 2021 (the start of the intervention in this model) to a given year “y”. We designate this cumulative RF difference for year *y,* CRFD^y^. We then determined, for each dietary scenario and year *y*, what level of reduction in annual CO_2_ emissions alone, relative to BAU, would yield the same CRFD^y^, and designate this annual CO*_2_* equivalent aCO_2_eq^y^ (see Figures 5-S1 to 5-S4 for details of these equivalences).

Critical features of aCO_2_eq are that it operates directly on RF inferred from combined trajectories of atmospheric levels of all GHGs, and thus can directly capture the effects of arbitrarily complex interventions, and that it equates the cumulative RF impact of an intervention over a specified time window to a single number: the sustained reductions in CO_2_ emissions that would have the same cumulative impact.

aCO_2_eq is closely related to, and motivated by similar goals as, CO_2_-forcing-equivalent (CO_2_-fe) emissions (Jenkins et al., 2018), which equates an arbitrary emission trajectory of all GHGs to a trajectory of CO_2_ emissions that would produce the same trajectory of RF, and GWP* (Allen et al., 2018, 2016; Cain et al., 2019), which uses various formulae to equate changes in GHG emissions to instantaneous CO_2_ pulses.

Figure 5 shows the aCO_2_eq for different scenarios for reference years 2050 (to capture short term impacts) and 2100 (Figure 5-S3 shows the full dependence of aCO_2_eq on the reference year). The aCO_2_eq^2100^ for PHASE-POD is -24.8 Gt/year. As global anthropogenic CO_2_ emissions are currently approximately 36 Gt/year, that PHASE-POD would have the same effect, through the end of the century, as a 68% reduction of CO_2_ emissions.

**Figure 5.**
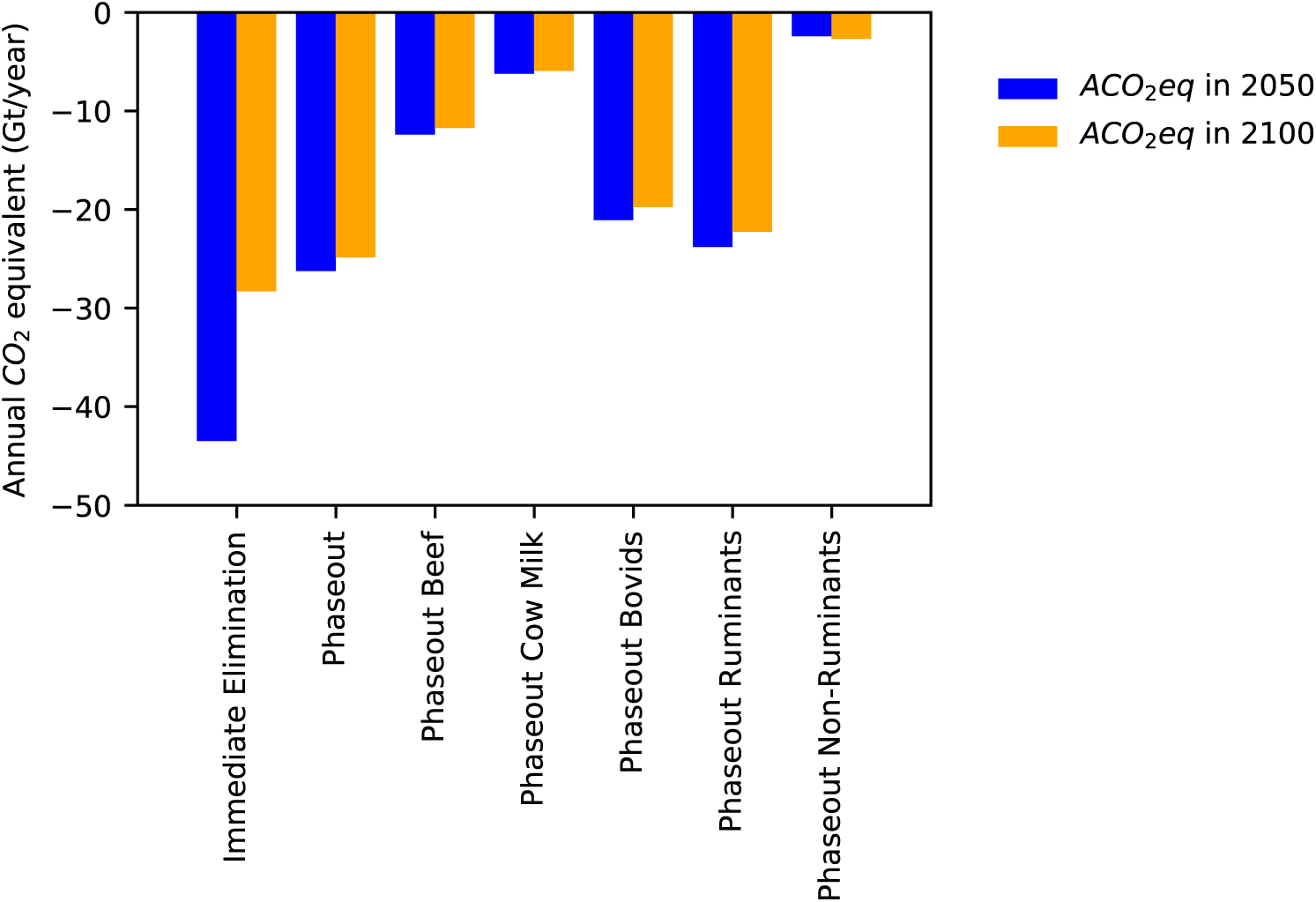
Annual *CO*_2_ equivalents (a*CO*_2_eq) of dietary scenarios. Bars show sustained reduction in annual *CO*_2_ emissions necessary to equal cumulative reduction in radiative forcing of given scenario in 2050 (blue) and 2100 (orange).

### Replacing ruminants achieves over 90 percent of climate benefit of eliminating animal agriculture

We next computed aCO_2_eq^2100^ for the 15 year phaseout of individual animal products and product categories (Figure 5 and 6A; Table 1), using the species- and product-specific emissions and land use values described above. Beef alone accounts for 47% of the benefits of phasing out all animal agriculture, and cow milk 24%. Meat and milk from bovids (cattle and buffalo) account for 79% of the climate opportunity. Although they provide less than 19% of the protein in the human diet (FAO, 2021), ruminants (cattle, buffalo, sheep and goats) collectively account for 90% of the aCO_2_eq^2100^ of all livestock.

**Figure 6.**
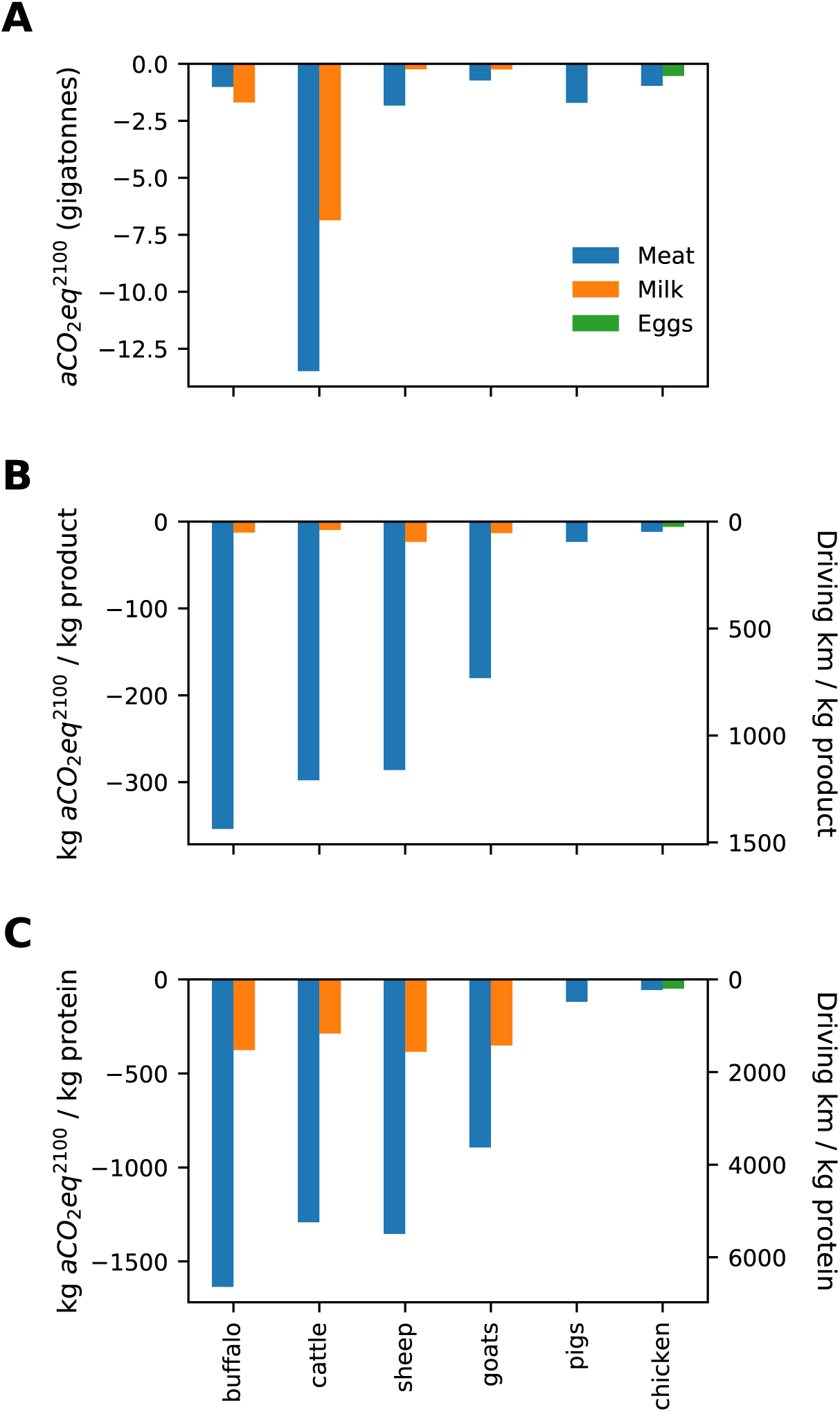
Emission equivalents and Emission Intensities of livestock products. (A) Total annualized CO_2_ equivalents through 2100, *aCO*_2_*eq*^2100^, for all tracked animal products, and Emission Intensities based on *aCO*_2_*eq*^2100^ on a per unit production (B) or per unit protein (C) basis. For (B) and (C) we also convert the values to driving equivalents using a value of 0.254 kg *CO*_2_*eq* per km driven of an internal combustion engine fueled sedan in the United States from life cycle analyses desribed in (Bieker, 2021).

**Table 1.** Product-specific emissions, land use and inferred impacts. Primary production data aggregated from FAOSTAT for 2019. Protein production data calculated from primary production data and protein conversion factors inferred from GLEAM. Emissions data based on protein production data and emission intensities from GLEAM. Land use data calculated from FAOSTAT protein production data and product-specific land use data from (Poore and Nemecek, 2018). Annualized CO_2_ equivalent emissions are for 2100 and calculated from atmospheric modeling results.

| Species | Commodity | Primary<br>Production<br>tonnes | Protein<br>Production<br>tonnes<br>protein | Emissions<br>CO<sub>2</sub><br>Mt | Emissions<br>CH<sub>4</sub><br>Mt | Emissions<br>N<sub>2</sub>O<br>Mt | Land<br>Use<br>Mkm<sup>2</sup> | aCO<sub>2</sub>eq<br>Gt/year | Emissions<br>Intensity<br>kg aCO<sub>2</sub> per<br>kg | Emissions<br>Intensity<br>kg aCO<sub>2</sub>eq per<br>kg protein | Driving<br>Equivalents<br>km driven per<br>kg |
| --- | --- | --- | --- | --- | --- | --- | --- | --- | --- | --- | --- |
| Buffalo | Meat | 4,290,212 | 619,200 | 29 | 5.00 | 0.20 | 1.0 | -1.0 | -354 | -1635 | 1394 |
| Cattle | Meat | 67,893,363 | 10,435,590 | 236 | 49.30 | 2.41 | 17.1 | -13.5 | -298 | -1292 | 1172 |
| Sheep | Meat | 9,648,245 | 1,354,398 | 32 | 5.02 | 0.33 | 2.5 | -1.8 | -286 | -1353 | 1126 |
| Goats | Meat | 6,128,372 | 821,383 | 21 | 3.34 | 0.11 | 0.8 | -0.7 | -180 | -893 | 709 |
| Pigs | Meat | 110,102,495 | 14,447,438 | 278 | 7.19 | 0.62 | 1.6 | -1.7 | -23 | -119 | 92 |
| Chickens | Meat | 123,898,557 | 17,393,440 | 306 | 0.29 | 0.52 | 1.3 | -1.0 | -12 | -56 | 47 |
| Ducks | Meat | 7,363,110 | 1,044,797 | 27 | 0.02 | 0.05 | 0.1 | -0.1 | -16 | -73 | 62 |
| Buffalo | Milk | 133,752,296 | 4,510,017 | 119 | 10.87 | 0.45 | 1.2 | -1.7 | -13 | -376 | 50 |
| Cattle | Milk | 712,883,270 | 23,889,273 | 338 | 37.63 | 1.78 | 6.3 | -6.9 | -10 | -287 | 38 |
| Sheep | Milk | 10,172,020 | 624,048 | 10 | 1.72 | 0.12 | 0.1 | -0.2 | -23 | -385 | 92 |
| Goats | Milk | 18,752,379 | 702,585 | 10 | 1.74 | 0.06 | 0.2 | -0.2 | -13 | -351 | 52 |
| Chickens | Eggs | 88,361,696 | 10,982,733 | 159 | 0.57 | 0.35 | 0.6 | -0.5 | -6 | -49 | 24 |

These product-specific aCO_2_eq’s can be interpreted on a per product unit (Figure 6B) or per protein unit (Figure 6C) as emissions intensities. Eliminating the consumption of a kilogram of beef, for example, is equivalent to an emissions reduction of 297 kg CO_2_. 38% (113 kg aCO_2_eq) comes from reduced emission, in line with the mean estimate of 99.5 kg CO_2_eq from a systematic meta analysis of GHG emissions from agricultural products (Poore and Nemecek, 2018), with the remaining 62% from biomass recovery.

As with the total numbers, ruminant meat has the largest emissions intensities, per unit (289 kg CO_2_eq per kg consumer product) and per protein (1,279 kg CO_2_eq per kg protein). The most efficient animal products on a per protein basis are chicken meat (56 kg CO_2_eq per kg protein) and eggs (49 kg CO_2_eq per kg protein), roughly 25 times lower than per protein emissions intensities for ruminant meat.

To connect these numbers to other sources of GHGs, we converted these emissions intensities to distances one would have to drive a typical 2021 model gas-fueled passenger car to produce the same emissions, based on a full life-cycle analysis of auto emissions (Bieker, 2021)(Figures 6B and C). One kg of beef, for example, has the same emissions impact as driving 1,172 km in a typical US car (or 339 miles per pound).

### Sensitivity to assumptions

Our default model assumes a gradual phaseout of animal agriculture over a period of 15 years, producing an aCO_2_eq^2100^ of -24.8 Gt/year . If we assume immediate elimination (Figure 2-S1), the aCO_2_eq^2100^ is -28.3 Gt/year (Figure 7A), a 14% increase in magnitude of the effect. If we assume a phaseout over 30 years (Figure 2-S2), the aCO_2_eq^2100^ is -21.3 Gt/year (Figure 7A), a 14% reduction.

**Figure 7.**
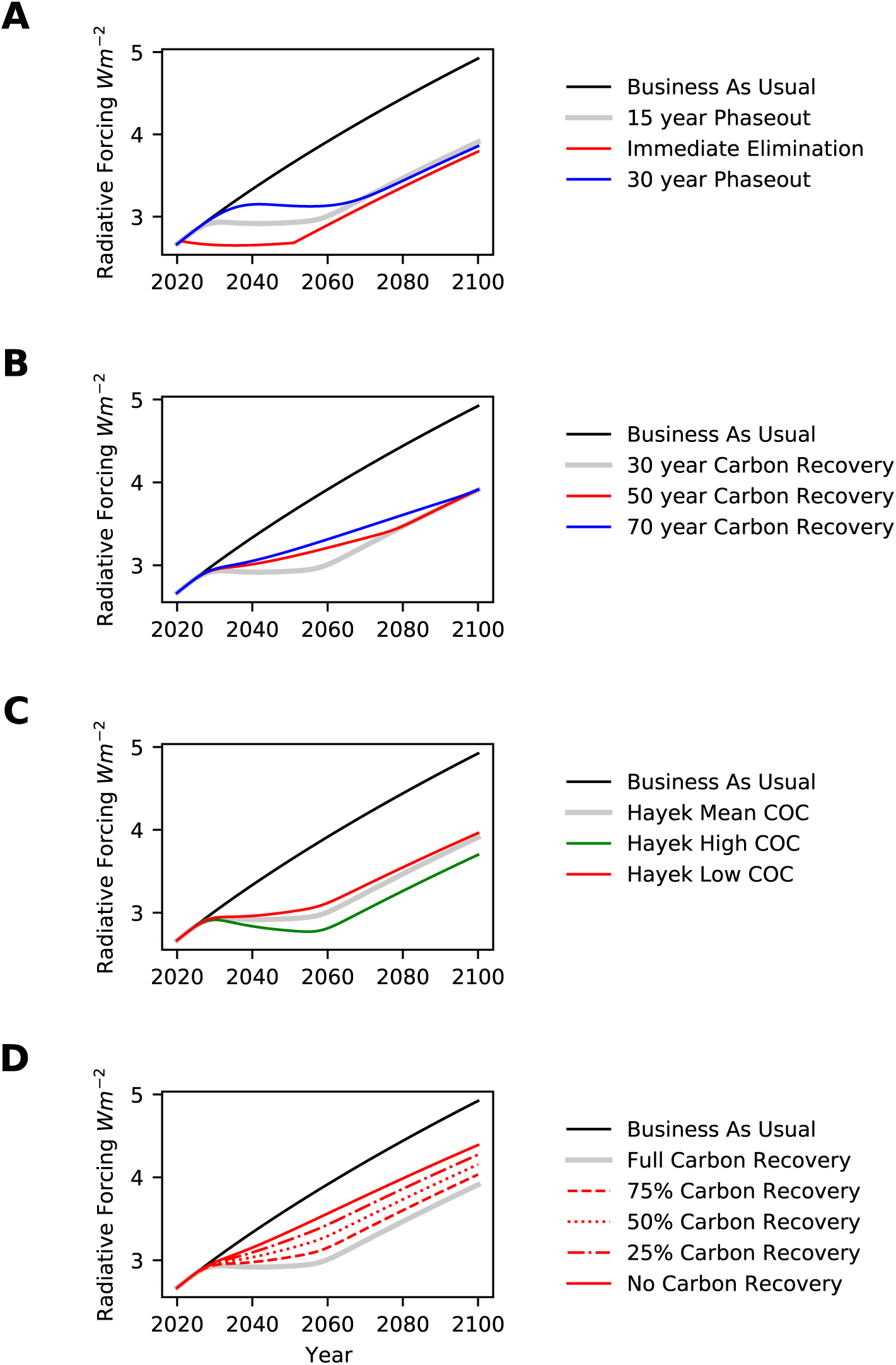
Sensitivity of impact of phaseout of animal agriculture to model assumptions. The grey line in each plot is PHASE-POD, the default scenario of 15 year phaseout, 30 year carbon recovery, livestock emissions from FAOSTAT, and a diverse plant replacement diet based on (Xu et al., 2021). (A) Effect of the immediate elimination of animal agriculture (red line) or a slower, 30 year, phaseout (blue line). (B) Effect of slower carbon recovery reaching completion after 50 years (red line) or 70 years (blue line). (C) Effect of using high (green line) or low (red line) estimates of above ground carbon recovery from (Hayek et al., 2020). (D) Effect of reducing either the efficiency or extent of carbon recovery.

Our default model also assumes that biomass will recover linearly over 30 years, following (Hayek et al., 2021), but there is considerable uncertainty in the literature, with estimates ranging from 25 to 70 years (Lennox et al., 2018; N’Guessan et al., 2019; Poorter et al., 2016). If we assume recovery takes 50 years (Figure 2-S3), the aCO_2_eq^2100^ is -22.4 Gt/year, and if it takes 70 years (Figure 2-S4), the aCO_2_eq^2100^ is -20.1 Gt/year, or reductions of 10% and 19% respectively (Figure 7B). We also note that passive recovery is not the only option. Further research is required to define optimal management practices for recovery of ecosystems currently impacted by animal agriculture and to estimate the rate and magnitude of their potential impact on climate. But there is evidence that deliberate, active management of ecosystem recovery to optimize for carbon sequestration could accelerate and increase the magnitude of carbon storage on land transitioning from intensive agricultural use (Griscom et al., 2017).

Estimates of the biomass recovery potential of land currently used for animal agriculture have a high degree of uncertainty. Using the low estimate (Figure 2-S5) of (Hayek et al., 2021), which addresses uncertainty in above-ground biomass yields an aCO_2_eq^2100^ of -21.2 Gt/year (Figure 7C), a 14% reduction in magnitude relative to the median value from (Hayek et al., 2021). Using the high estimate (Figure 2-S6) of (Hayek et al., 2021) yields an aCO_2_eq^2100^ of -28.1 Gt/year (Figure 7C), an increase in magnitude of 13% increase.

A major area of uncertainty not addressed by (Hayek et al., 2021) is the extent to which the carbon recovery potential of land that transitions away from use in animal agriculture would be realized in the face of other land use pressures. (Hayek et al., 2021) accounts for the land needed to replace animal derived foods in the global diet, but not for other potential large-scale non-food uses such as biofuel production. While it is beyond the scope of this work to model these uses explicitly, Figure 7D shows the expected RF trajectories if we assume reduced recovery fractions of 25% (Figure 2-S7), 50% (Figure 2-S8), 75% (Figure 2-S9) and 100% (Figure 2-S10), which yield aCO_2_eq^2100^ of -21.6, -18.3, -15.0, and -11.6 Gt/year respectively, highlighting the importance of carbon recovery in realizing the climate potential of ending animal agriculture. It is important to note that there is substantial variance in the biomass potential between regions and ecosystems, and recent modeling work by (Strassburg et al., 2020) indicates that half of the biomass recovery potential of land currently used for agriculture could be realized by restoration of 25% of the relevant land.

Our estimate of global emissions due to animal agriculture based on FAO data and analyses of 1.6 Gt CO_2_, 122 Mt CH_4_ and 7.0 Mt N_2_O differ in key ways from recent estimates of (Xu et al., 2021) of 3.2 Gt CO_2_, 102 Mt CH_4_ and 3.9 Mt N_2_O. Using these emissions estimates for livestock (Figure 2-S11) yields an aCO_2_eq^2100^ of PHASE-POD of -23.6 Gt/year (Figure 7-S1A), a 5% decrease in magnitude.

The models described above assume that the protein currently obtained from animal products would be replaced with a diverse plant based diet, scaled to replace animal products on a protein basis, and agriculture emissions data from FAOSTAT. We considered as an alternative emissions projected from a diverse plant based diet based on data from (Xu et al., 2021), scaled to replace animal products on a protein basis. This replacement diet (Figure 2-S12) yields an aCO_2_eq^2100^ for PHASE-POD of animal agriculture of -23.7 Gt/year (Figure 7-S1B), a 5% decrease in magnitude.

In some areas, the removal of land from use in animal agriculture may lead to an increase in wild ruminant population. Although this is difficult to model globally, this would offset some of the beneficial impacts of reductions in methane emissions from livestock (Kelliher and Clark, 2010).

This analysis only considered consumption of terrestrial animal products, neglecting emissions and land use (via feed production) associated with seafood capture and aquaculture. While the land and emissions impact of seafood consumption has received comparably little attention, several studies have pointed to at least 500 Mt of CO_2_ equivalent emissions per year from seafood (MacLeod et al., 2020; Parker et al., 2018; Poore and Nemecek, 2018). Recent work has also suggested that the disruption of carbon storage due to seafood harvesting via trawling repartitions from 0.58 up to 1.47 Gt CO_2_ equivalent carbon per year from sediment into the water column, with the potential to drive atmospheric increases of similar magnitude (Sala et al., 2021).

Widely used climate models consider temporal and spatial variation in emissions; feedback between a changing climate and anthropogenic and natural emissions, carbon sequestration, atmospheric chemistry and warming potential; the impact of climate on human social, political and economic behavior. Ours does not. We ran our model on emissions data from the pathways described in (Riahi et al., 2017) and compared our atmospheric level and RF outputs to theirs, and found them to be in broad qualitative agreement. Thus, while other models could provide more precise estimates, we do not believe they would alter our major conclusions.

## Discussion

Our analysis has provided a quantitative estimate of the potential climate impact of a hypothetical, radical global change in diet and agricultural systems. We have shown that the combined benefits of removing major global sources of CH_4_ and N_2_O, and allowing biomass to recover on the vast areas of land currently used to raise and feed livestock, would be equivalent to a sustained reduction of 25 Gt/year of CO_2_ emissions.

Crucially eliminating the use of animals as food technology would produce substantial negative emissions of all three major GHGs, a necessity, as even the complete replacement of fossil fuel combustion in energy production and transportation will no longer be enough to prevent warming of 1.5°C (Clark et al., 2020; IPCC, 2018).

The transition away from animal agriculture will face many obstacles and create many challenges. Meat, dairy and eggs are a major component of global human diets (FAO, 2021), and the raising of livestock is integral to rural economies worldwide, with more than a billion people making all or part of their living from animal agriculture.

Although animal products currently provide, according to the most recent data from FAOSTAT, 18% of the calories, 40% of the protein and 45% of the fat in the human food supply, they are not necessary to feed the global population. An estimated 375 million people already live on entirely plant based diets (Heinrich-Böll-Stiftung and the Earth Europe, 2014), and existing crops could replace the calories, protein and fat from animals with a vastly reduced land, water, GHG and biodiversity impact, requiring only minor adjustments to optimize nutrition (Springmann et al., 2018b).

The economic and social impacts of a global transition to a plant based diet would be acute in many regions and locales (Newton and Blaustein-Rejto, 2021), a major obstacle to their adoption. It is likely that substantial global investment will be required to ensure that the people who currently make a living from animal agriculture do not suffer when it is reduced or replaced. And, while it is expected that the phaseout of animal agriculture would lead to global increases in food availability as edible crops cease to be diverted for animal feed (Vågsholm et al., 2020), investment will also be required to prevent local food insecurity in regions where wide-scale access to a diverse and healthy plant-based diet is currently lacking and to ensure proper nutrition. But, in both cases, these investments must be compared to the economic and humanitarian disruptions of significant global warming (Howard and Sylvan, 2021; Stehfest et al., 2019).

Although, as discussed above, there are many uncertainties in our estimates, our assumption that “business as usual” means animal agriculture will continue at current levels was highly conservative, as rising incomes are driving ongoing growth in global animal product consumption (OECD-FAO Agricultural Outlook 2020-2029). It is estimated that global demand for animal based foods will increase by nearly 70 percent by 2050 (Searchinger et al., 2019). For example, using land use data from (Poore and Nemecek, 2018) and consumption data from FAOSTAT, extending the current diet of wealthy industrialized countries (OECD) to the current global population would require an additional 35 million km^2^ to support livestock production - an area roughly equal to the combined area of Africa and Australia.

While such an expansion may seem implausible, even partial destruction of Earth’s critical remaining native ecosystems would have catastrophic impacts not just on the climate, but on global biodiversity (IPBES, 2019; Newbold et al., 2015; World Wildlife Fund, 2020) and human health (Clark et al., 2019; Maron et al., 2018; Oliver et al., 2015; Satija et al., 2017; Springmann et al., 2016; Strassburg et al., 2020; Tilman and Clark, 2014).

Given these realities, even with the many challenges that upending a trillion dollar a year business and transforming the diets of seven billion people presents, it is surprising that changes in food production and consumption are not at the forefront of proposed strategies for fighting climate change. Although all of the strategies presented as part of the recent Intergovernmental Panel on Climate Change (IPCC) report on steps needed to keep global warming below 1.5°C (IPCC, 2018) acknowledge the need for significant negative emissions, none propose even a reduction in per capita livestock consumption below current levels (Figure 8).

**Figure 8.**
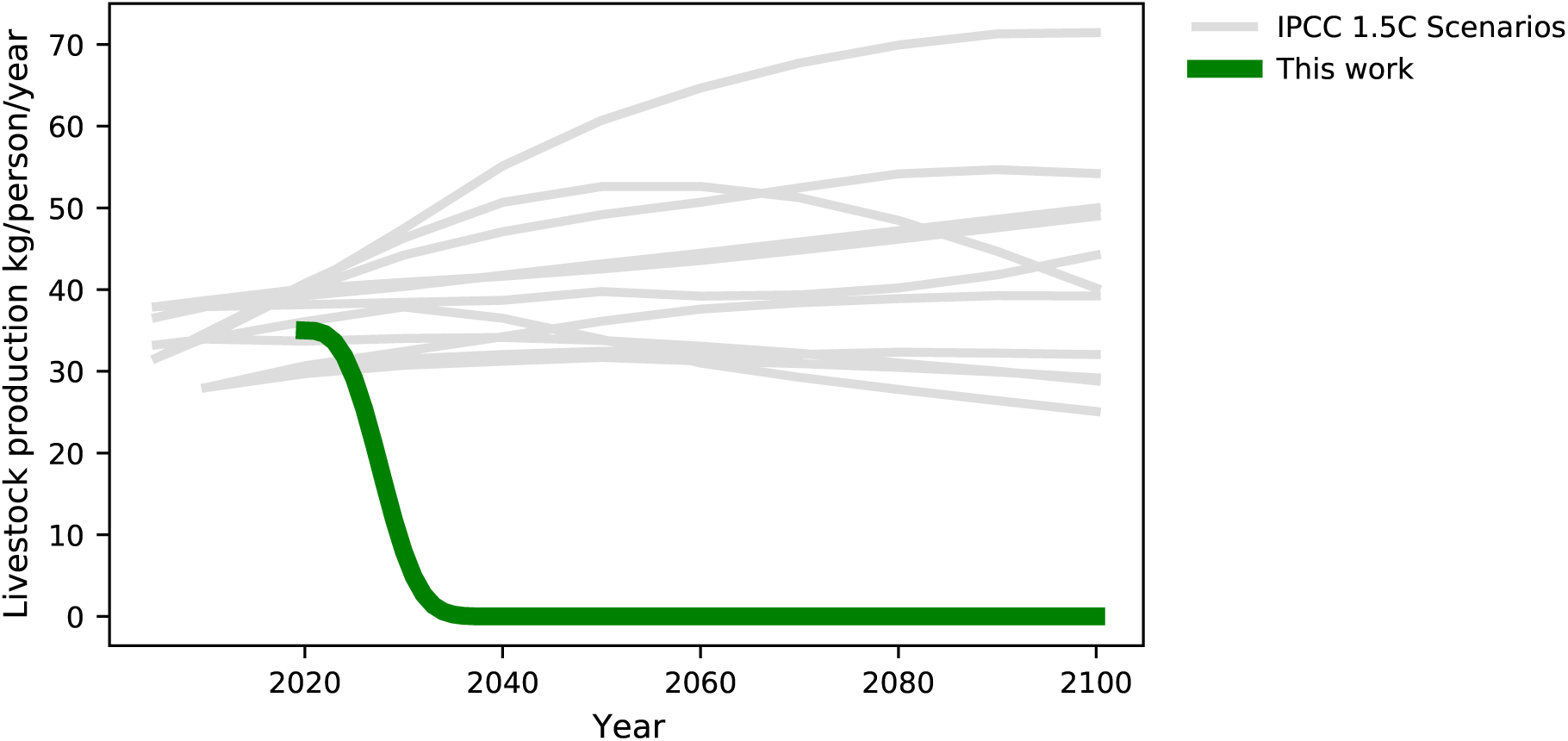
Projected per capita livestock production in SSP/IAM RF 1.9 scenarios. We downloaded data for the Shared Socioeconomic Pathways (SSPs) (Riahi et al., 2017) from the SSP database (Version 2.0; last updated December 2018), and plot here the inferred per capita livestock production for scenarios meant to reach an RF target of 1.9 in 2100. While there is widespread acknowledgement of the impact that ongoing animal agriculture has on the climate, it is notable that most of these scenarios, which represent the most aggressive proposed mitigation strategies in this modeling framework, anticipate an increase in per capita livestock consumption, and none anticipate any significant reduction below current levels, in contrast to the complete elimination we propose here.

Even if the negative emission technology the IPCC anticipates, BECCS (bio-energy combined with carbon capture and storage), proves to be viable at scale, it will require large amounts of land (Anderson and Peters, 2016), and the only way to get that land without massive collateral damage is by displacing animal agriculture, primarily land-intensive ruminants. Thus, all potential solutions to the climate crisis likely require some form of large scale dietary change.

It is important to emphasize that, as great as the potential climate impact of ending animal agriculture may be, even if it occurred, and even if all of the benefits we anticipate were realized, it would not be enough on its own to prevent catastrophic global warming. Rather we have shown that global dietary change provides a powerful complement to the indispensable transition from fossil fuels to renewable energy systems. The challenge we face is not choosing which to pursue, but rather in determining how best to overcome the many social, economic and political challenges incumbent in implementing both as rapidly as possible.

## Methods

### Data and Code Availability

Analyses were carried out in Python using Jupyter notebooks. All data, analyses and results presented here are available at github.com/mbeisen/LivestockClimateImpact.

### Updating Estimates of Emissions from Animal Agriculture

We obtained country, species, herd and product type specific CO_2_, CH_4_ and N_2_O emission data for terrestrial livestock from the public version of GLEAM 2.0 (MacLeod et al., 2018) downloaded from http://www.fao.org/gleam/results/en/. GLEAM contains data for cattle, buffalo, sheep, goats, pigs and chickens, and attributes emissions to meat, milk and eggs. Although GLEAM further breaks down emissions based on herd type and production system, we used aggregate data for all herds and production types in the country. We did not include CO_2_ emissions linked to land-use change, as this is associated with increases in livestock production which are explicitly not considered by our model.

We obtained livestock production data for 2019 (the most recent year available) from the “Production_LivestockPrimary” datafile in FAOSTAT (FAO, 2021). We extracted from Production_LivestockPrimary the amount (in tonnes), for all countries, of primary domestic production of meat from cattle, buffalo, sheep, goat, pig, chicken and duck, milk from cows, buffalo, sheep and goat, and eggs from poultry. We computed meat and protein yields from the carcass weight data reported by GLEAM.

We scaled the GLEAM emission data to current production data from FAOSTAT, using GLEAM data for entire herds based on carcass weight for meat, and production weight for milk and eggs. As GLEAM does not provide data for ducks, we used values for chicken. The scaling was done using country-specific livestock production data from FAOSTAT and regional data from GLEAM.

### Estimating species-specific land use

We combined livestock production data with average species and product-specific land use data from (Poore and Nemecek, 2018) to estimate species, product and country-specific land use data associated with animal agriculture. We use data for cattle meat for buffalo meat, and cow milk for milk from buffalo, goat and sheep. The data are reported in m*m*^2^ (*year*)(100*g protein*)^−1^ except for milk which is reported in *m*^2^ (*year*)(*liter*)^−1^ which we convert to *m*^2^ (*year*)(*kg primary production*) using conversion factors inferred from GLEAM, which reports both protein and primary production data.

The total land use for animal agriculture inferred from this analysis is 33.7 million km^2^, almost identical to the 33.2 million km^2^ estimated by (Hayek et al., 2021) from satellite imagery.

### Emissions from Agriculture

We used the Environment_Emissions_by_Sector_E_All_Data_(Normalized) data table from FAOSTAT, projecting from the most recent year of 2017 to 2019 by assuming that the average annual growth from 2000 to 2017 continued in 2018 and 2019.

### Replacement Diets

We modeled agricultural emissions under a business as usual (BAU) diet as remaining at 2019 levels. When modeling reductions in livestock consumption, we assumed protein from livestock products would be replaced with equivalent amount of protein from current food crops, and used per unit protein emission intensities computed from FAOSTAT to infer emissions from this replacement diet. As an alternative we used emission intensities from (Xu et al., 2021) as described in the Sensitivity section. For diets involving the removal of one or more specific animal products, we scaled these dietary replacement emissions by the fraction of animal protein obtained from that product, and scaled biomass recovery by the fraction of animal agriculture land attributed to that product.

### Replacement Scenarios

In all scenarios we assume annual non-agricultural emissions remain fixed at 2019 levels through 2100. For a BAU diet we added in total agricultural emissions from the FAOSTAT “Emissions Shares” data table, effectively fixing total emissions at 2019 levels. We assumed a 15 year phaseout of animal agriculture with an accelerated rate of conversion from BAU to PHASE-POD. The specific formula we use is 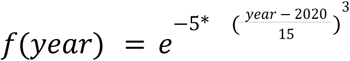 yielding the conversion dynamics shown below:

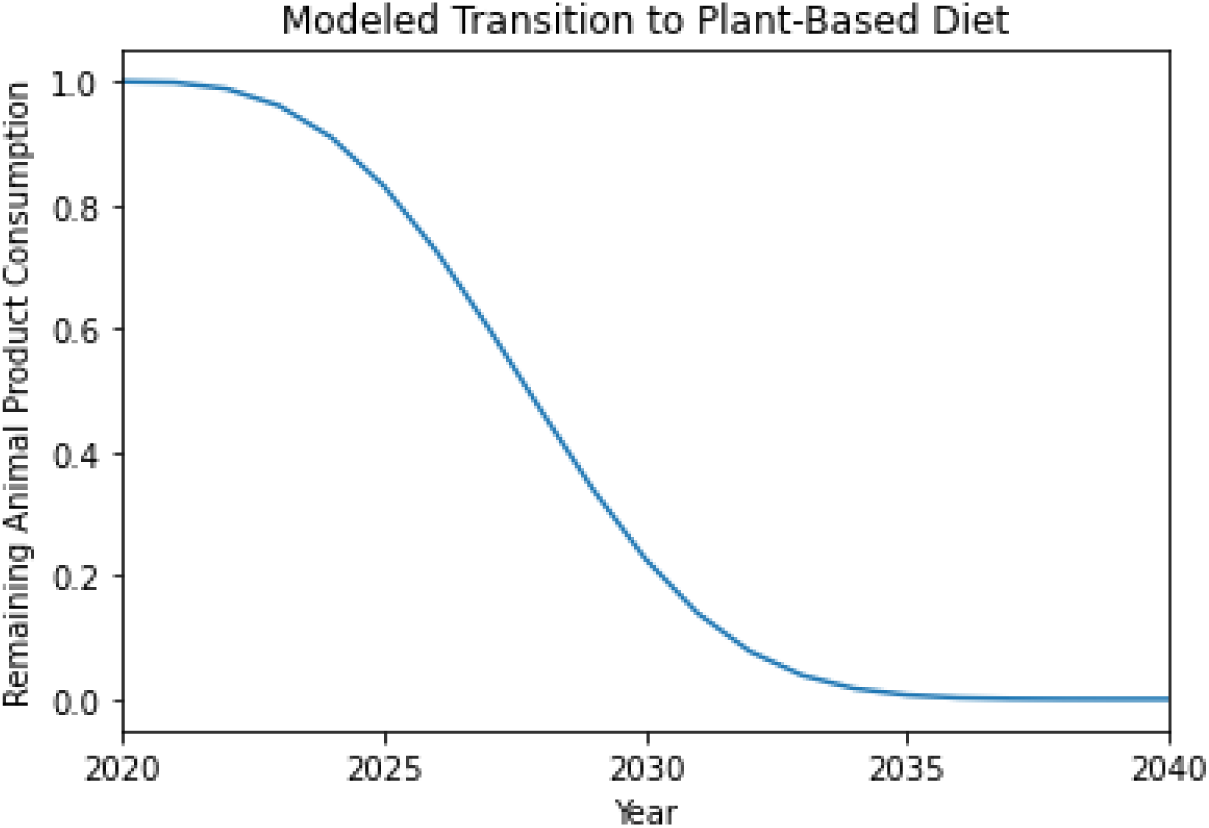

We also include in the supplemental data a version of the analysis in which the hypothetical transition is instantaneous (IMM-POD).

As the transition from BAU to PHASE-POD occurs, agriculture linked emissions are set to

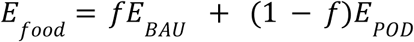

Where *f*is the fraction of the global diet that is still BAU.

We assume that, when animal-derived food consumption is reduced in a year by a fraction Δ*f*, that carbon recovery on a corresponding fraction of land begins immediately and continues at a constant rate until it reaches 100% after 30 years (Hayek et al., 2021) (see also Figure 7 for 50 and 70 year recovery timelines).

### Converting between emissions and atmospheric concentrations of GHGs

The total mass of gas in the atmosphere is 5.136 * 10^21^ g, at a mean molecular weight of 28.97 g/mole (Walker, 1977), or 1.77e+20 total moles of gas. Hence 1 ppb is 1.77*10^11^ moles and 1 ppm is 1.77 * 10^14^ moles.

We therefore use conversions from mass in Gt to ppb/ppm as follows:

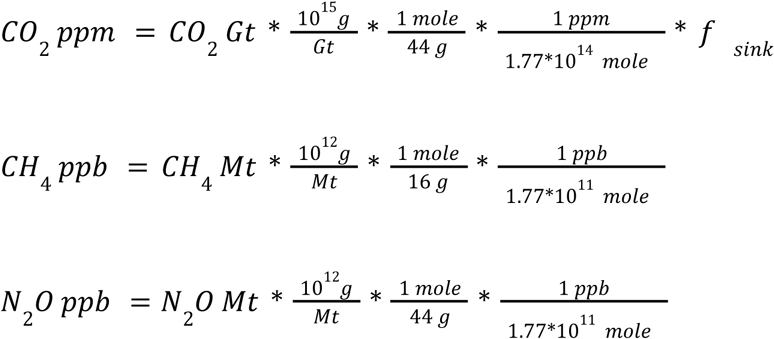

We use an *f_sink_* value of 0.50 reflecting the observation that approximately half of terrestrial CO_2_ emissions end up in land or ocean sinks rather than the atmosphere (Houghton, 2003).

### Estimating global non-anthropomorphic emissions

Both CH_4_ and N_2_O decay at appreciable rates, with half-lives of approximately 9 years for CH_4_ ^(^Morgenstern et al., 2017^)^ and 115 years for N_2_O (Prather et al., 2015), although these estimates are being continuously updated (Saunois et al., 2020). We balanced the corresponding decay equations against historical emissions and atmospheric levels, inferring unaccounted for and presumably non-anthropogenic sources leading to mole fraction equivalent increases of CH_4_ of 25 ppb/year and N_2_O of 1.0 ppb/year.

### Projections of Atmospheric Gas Levels

We ran projections on an annual basis starting in 2020 and continuing through 2100. For each gas:

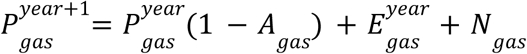

where:

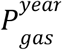 is the atmospheric concentration of *gas* in *year* in ppb for CH_4_ and N_2_O and ppm for CO_2_

*A_gas_* is the annual decay of *gas* and is equal to 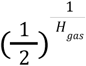 where *H_gas_* is the half-life of *gas* (we assume that CO_2_ does not decay)

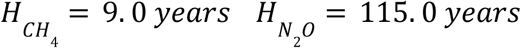

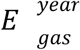 is the emissions of *gas* in *year* converted to atmospheric ppb for CO_2_ and N_2_O and ppm for CO_2_ as described above

*N_gas_* is the constant term to account for emissions not captured in *E*

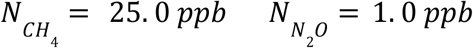

Starting conditions were obtained from the US National Ocean and Atmospheric Administration Global Monitorial Laboratory (“Carbon Cycle Greenhouse Gases,” n.d.):

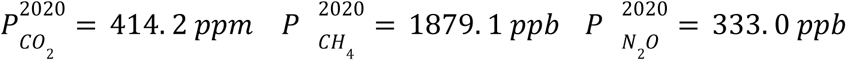

### Radiative Forcing

We adopt the commonly used formula for radiative forcing (RF) which derives from (Myhre et al., 1998; Ramaswamy et al., 2001) as modified in the climate modeling program MAGICC6 (Meinshausen et al., 2011).

Given atmospheric concentration of *C* ppm CO_2_, *M* ppb CH_4_ and *N* ppb N_2_O

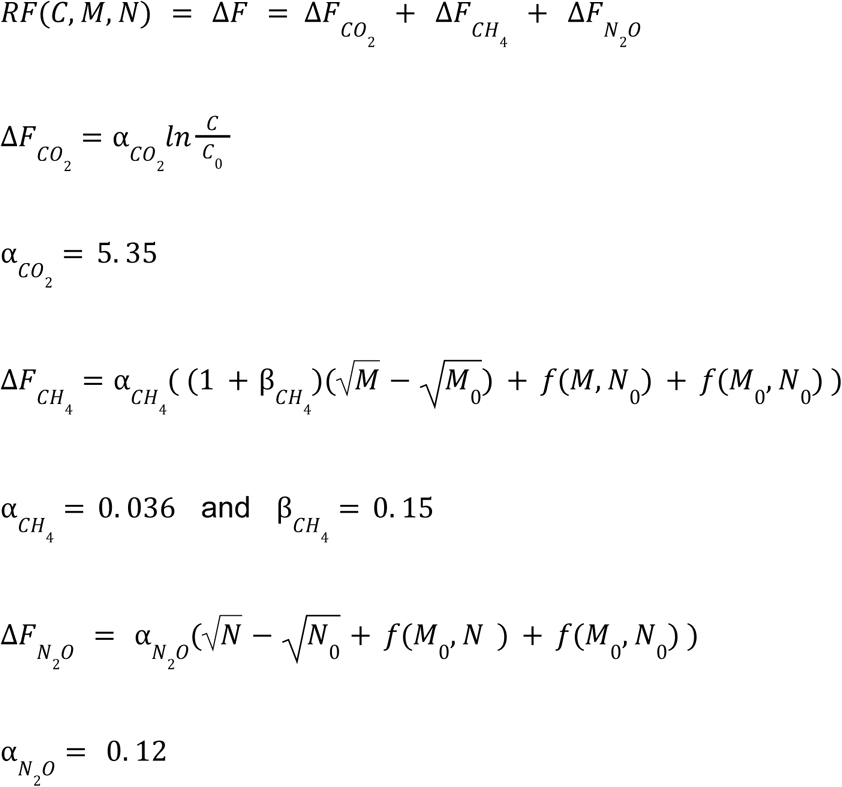

The function 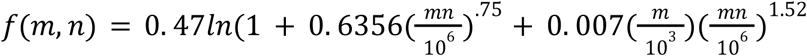 captures the overlap in spectra between CH_4_ and N_2_O.

*C_0_*, *M_0_* and *N_0_* are the preindustrial levels of the corresponding gasses.

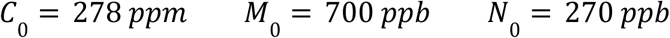

### Computing Emissions and Land Carbon Opportunity Cost

We define the combined emissions and land carbon opportunity cost (ELCOC) of animal agriculture as 2Δ*C* where

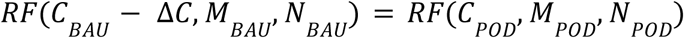

The factor of 2 accounts for the half of CO_2_ emissions that go to terrestrial sinks.

### Computing Carbon Emissions Budgets for RF 2.6 and 1.9

As the RF calculation used in MAGICC6 account for other gasses and effects beyond the three gasses used here, we used multivariate linear regression as implemented in the Python package scikit-learn to predict the complete RF output of MAGICC6 using data data downloaded from the Shared Socioeconomic Pathways (SSPs) (Riahi et al., 2017). The model was trained on atmospheric concentrations of CO_2_, CH_4_ and N_2_O to predict the difference between the MAGICC6 RF and the RF calculated using only CO_2_, CH_4_ and N_2_O. Then, for timepoints in our scenarios we computed RF as above from CO_2_, CH_4_ and N_2_O concentrations, and added to this the adjustment from the linear regression model. We use this RF in Figures 3 and 4.

In the SSP file:

C = Diagnostics|MAGICC6|Concentration|CO_2_
M = Diagnostics|MAGICC6|Concentration|CH_4_
N = Diagnostics|MAGICC6|Concentration|N_2_O

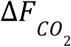 = Diagnostics|MAGICC6|Forcing|CO_2_

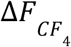 = Diagnostics|MAGICC6|Forcing|CH_4_

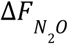 = Diagnostics|MAGICC6|Forcing|N_2_O

MAGICC6 RF = Diagnostics|MAGICC6|Forcing

### aCO_2_eq

To compute aCO_2_eq^y^, the annual CO_2_ equivalent emission change of each emissions scenario, we first ran scenarios in which annual CO_2_ emissions were reduced from 50 Gt/year to 1 Gt/year in increments of 1 Gt/year, then from 1 Gt/year to 10 Mt/year in increments of 10 Mt/year, and then from 1 Mt/year to 100 kT/year in increments of 100 kT/year. For each of these calibration scenarios, and for all years *y* from 2021 to 2100, we computed the total RF difference between the calibration scenario and BAU, from 2021 to *y*.

For each multi-gas emissions scenario, we similarly computed CRFD^y^, and determined what constant level of reduction in annual CO_2_ emissions alone by interpolation using the CRFD^y^ of the calibration scenarios, and designate this annual CO*_2_* equivalent aCO_2_eq^y^.

### Product equivalents

To compute per product unit and per protein emissions equivalents, we divided aCO_2_eq^2100^ for immediate elimination of the product (in kg CO_2_eq/year) by the annual production of the product (in kg production/year) yielding a per product unit emission equivalent measured in kg CO_2_eq per kg production.

For example, assuming, as our model does, that emissions and land use scale with consumption, if annual beef production were reduced by one tonne (1,000 kg) per year, it would result in corresponding annual reductions of -3,476 kg CO_2_, -726 kg CH_4_ and -36 kg N_2_O, and would initiate 30 year biomass recovery of 6,050,000 kg of CO_2_ equivalent carbon on 25.2 ha of land.

The cumulative reduction in RF, through 2100, of such annual emissions reductions and biomass recovery would be equivalent to a CO_2_ emission reduction of 199,000 kg/year. The ratio of these two rates, -199,000 kg CO_2_eq/year over 1,000 kg beef/year yields -199 kg CO_2_eq per kg beef as a measure of the warming impact of one kg of beef. Adjusting this for the dressing percentage of beef (the values reported by FAO, and used in these calculations, are carcass weight, of which only approximately ⅔ ends up as a consumer product) yields the values shown in Figure 6.

For all meat products we scaled the production amount by a typical dressing percentage of ⅔ to convert to consumer product units. For protein unit equivalents we used protein yields from GLEAM. To convert to driving equivalents we used a value of .254 kg CO_2_eq per km driven taken from a full life cycle analysis of 2021 sedans in the United States from (Bieker, 2021).

## Supporting information

Peer Reviews and Responses

---

**Figure 1-S1.**
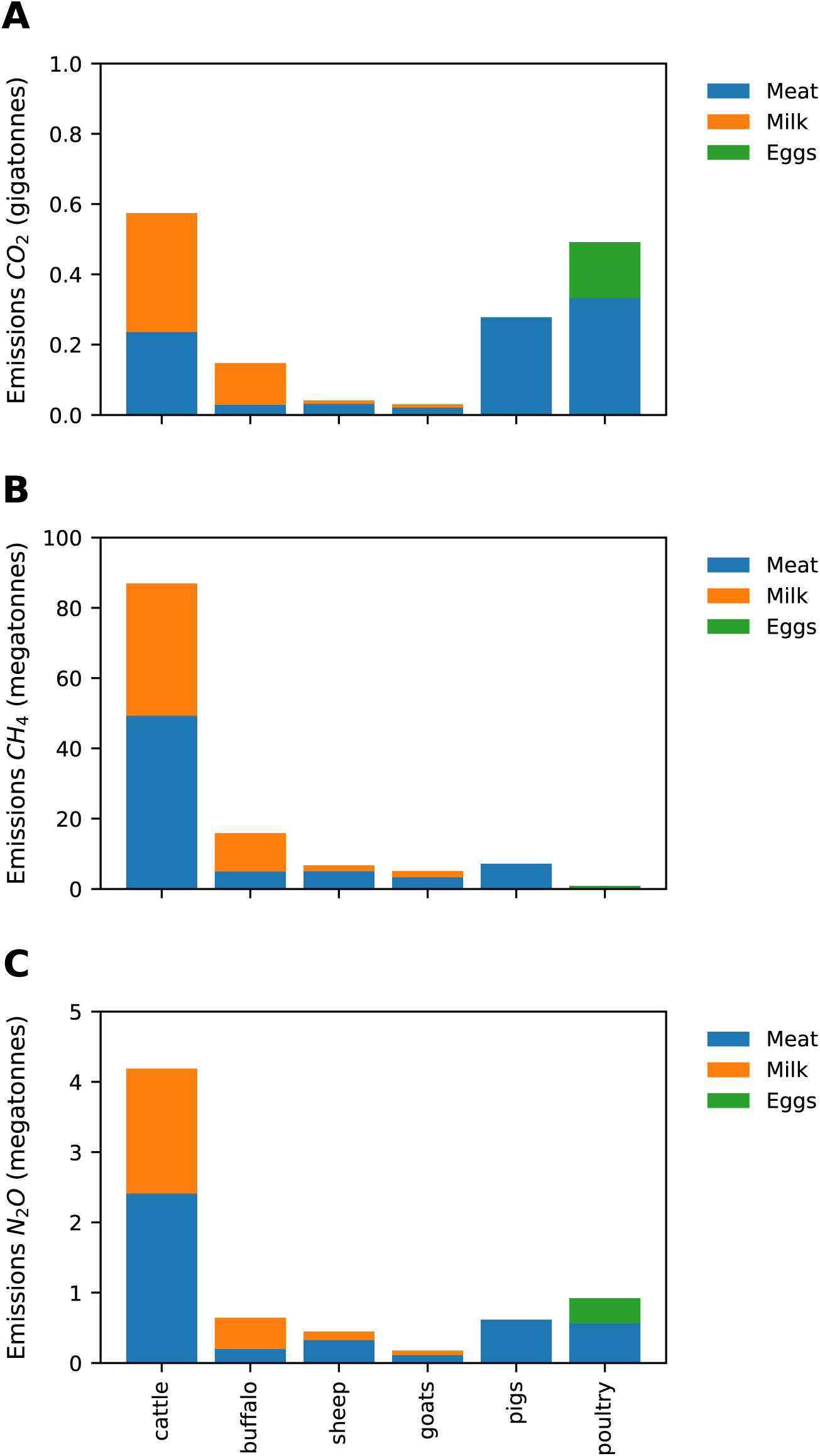
Gas-specific emission footprints of animal agriculture. Assembled from species, product and country-specific production data from FAOSTAT for 2018 and species, product, region and greenhouse gas-specific emissions data from GLEAM (MacLeod et al., 2018).

**Figure 2-S1.**
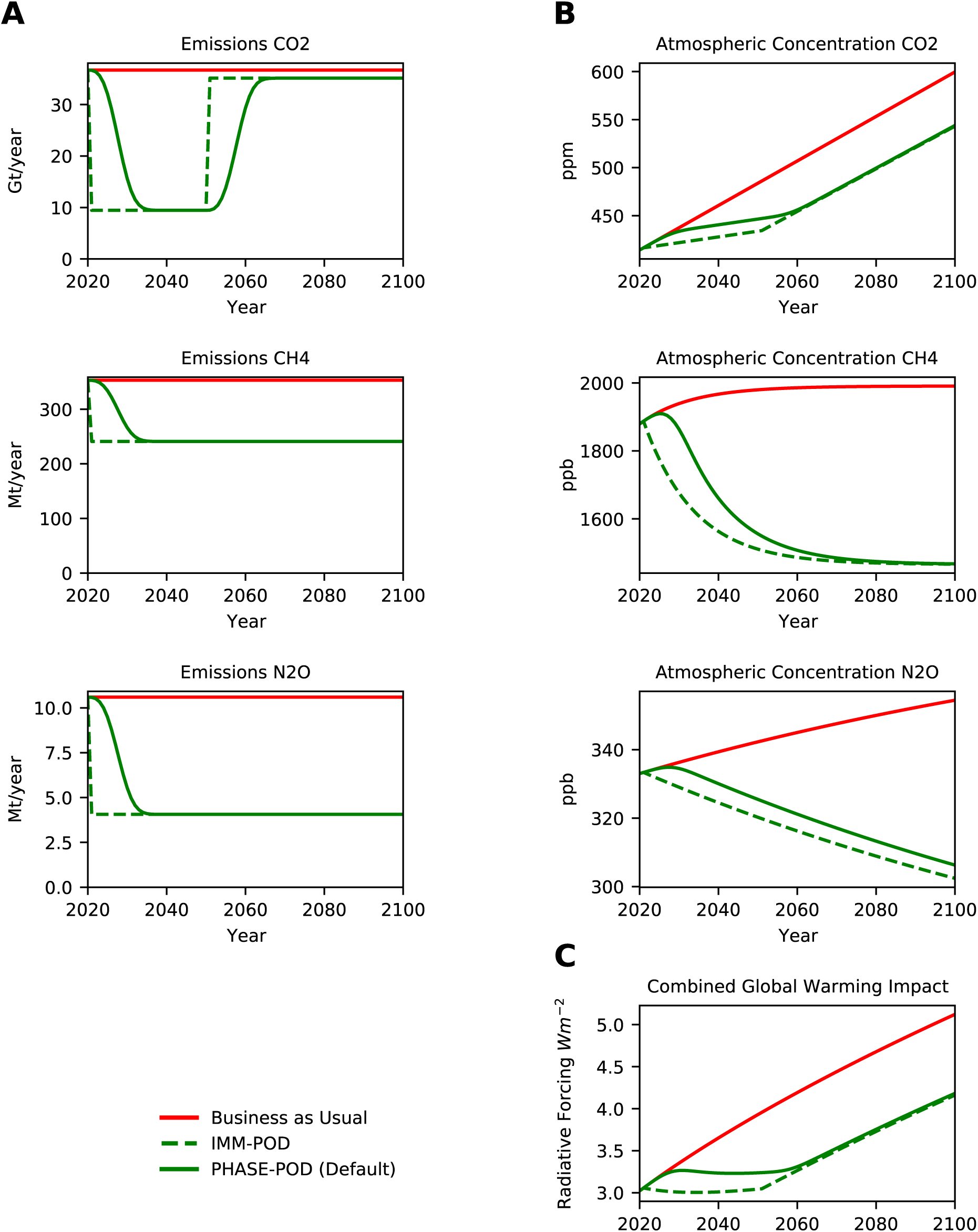
15yr Phaseout vs. Elimination. (A) Projected annual emissions of *CO*_2_, *CH*_4_ and *N*_2_*O* for each scenarios. (B) Projected atmospheric concentrations of *CO*_2_, *CH*_4_ and *N*_2_*O* under each emission scenario. (C) Radiative Forcing (RF) inferred from atmospheric concentrations in (B) by formula of (Myhre et al., 1998; Ramaswamy et al., 2001) as modified in MAGICC6 (Meinshausen et al., 2011). Only differences between PHASE-POD default assumptions (15yr phaseout, 30yr carbon recovery, 100% carbon recovery, BAU non-agriculture emissions, FAO crop replacement, and FAO animal ag emissions) are given.

**Figure 2-S2.**
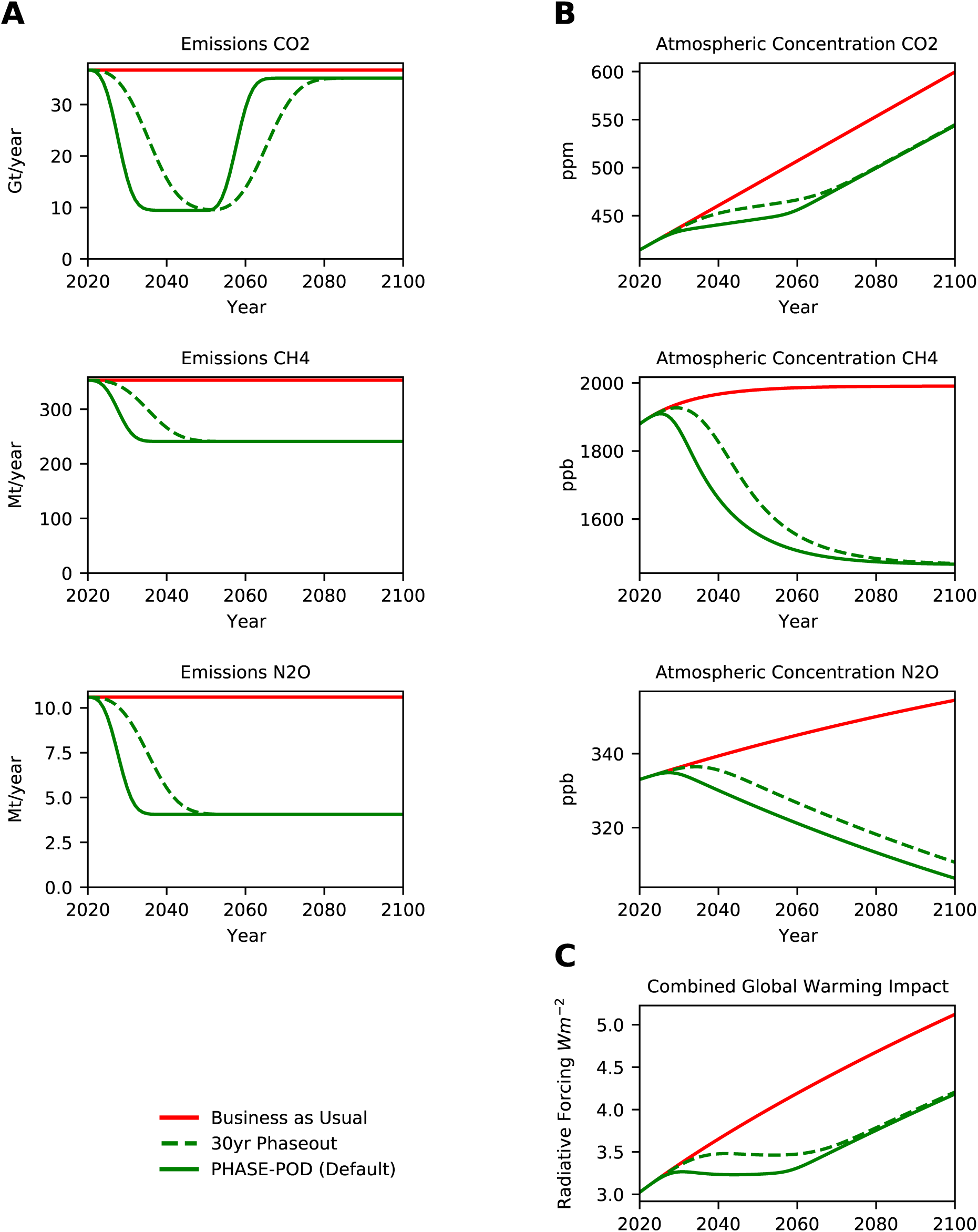
15yr vs. 30yr Phaseout. (A) Projected annual emissions of *CO*_2_, *CH*_4_ and *N*_2_*O* for each scenarios. (B) Projected atmospheric concentrations of *CO*_2_, *CH*_4_ and *N*_2_*O* under each emission scenario. (C) Radiative Forcing (RF) inferred from atmospheric concentrations in (B) by formula of (Myhre et al., 1998; Ramaswamy et al., 2001) as modified in MAGICC6 (Meinshausen et al., 2011). Only differences between PHASE-POD default assumptions (15yr phaseout, 30yr carbon recovery, 100% carbon recovery, BAU non-agriculture emissions, FAO crop replacement, and FAO animal ag emissions) are given.

**Figure 2-S3.**
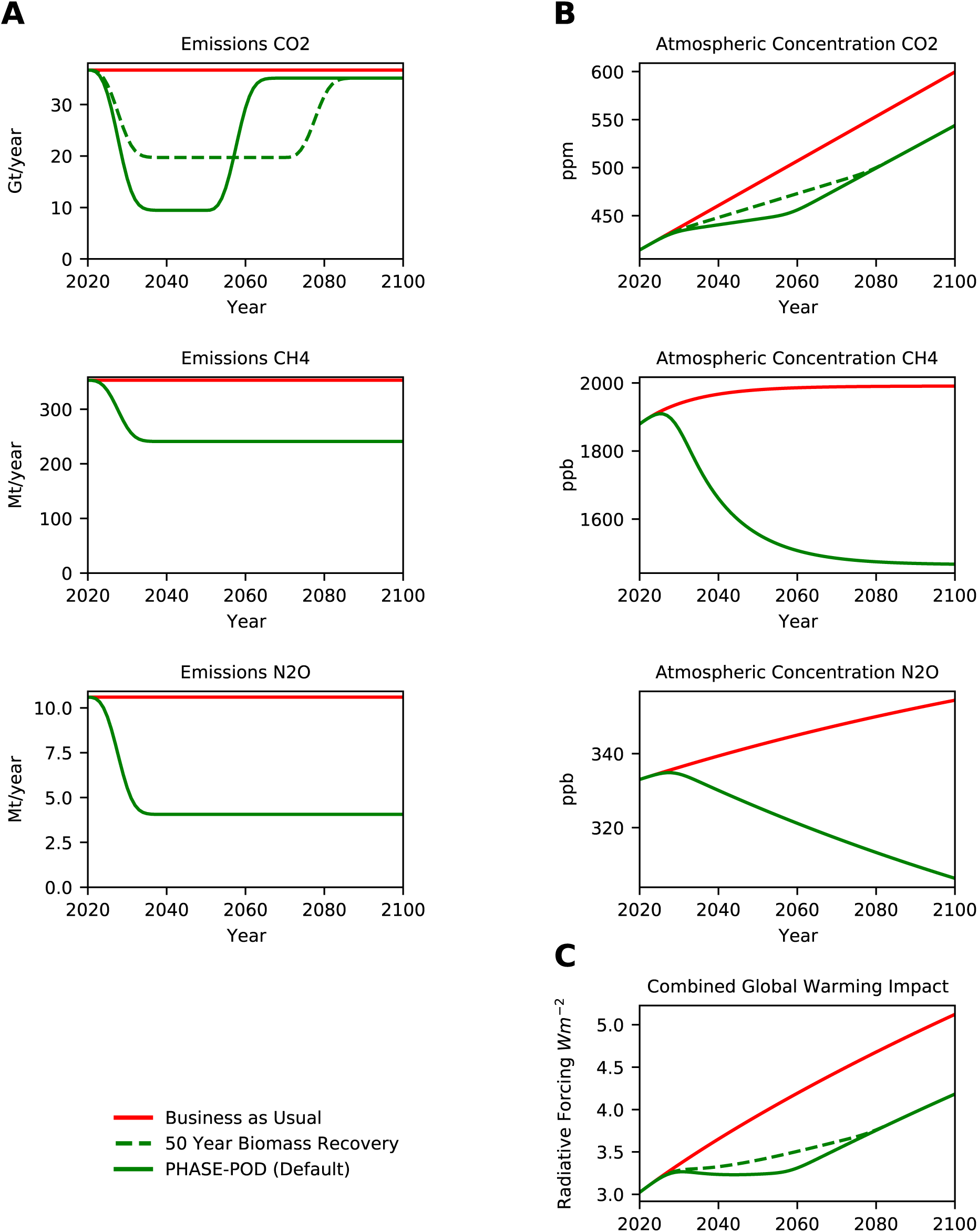
30yr vs. 50yr Biomass Recovery. (A) Projected annual emissions of *CO*_2_, *CH*_4_ and *N*_2_*O* for each scenarios. (B) Projected atmospheric concentrations of *CO*_2_, *CH*_4_ and *N*_2_*O* under each emission scenario. (C) Radiative Forcing (RF) inferred from atmospheric concentrations in (B) by formula of (Myhre et al., 1998; Ramaswamy et al., 2001) as modified in MAGICC6 (Meinshausen et al., 2011). Only differences between PHASE-POD default assumptions (15yr phaseout, 30yr carbon recovery, 100% carbon recovery, BAU non-agriculture emissions, FAO crop replacement, and FAO animal ag emissions) are given.

**Figure 2-S4.**
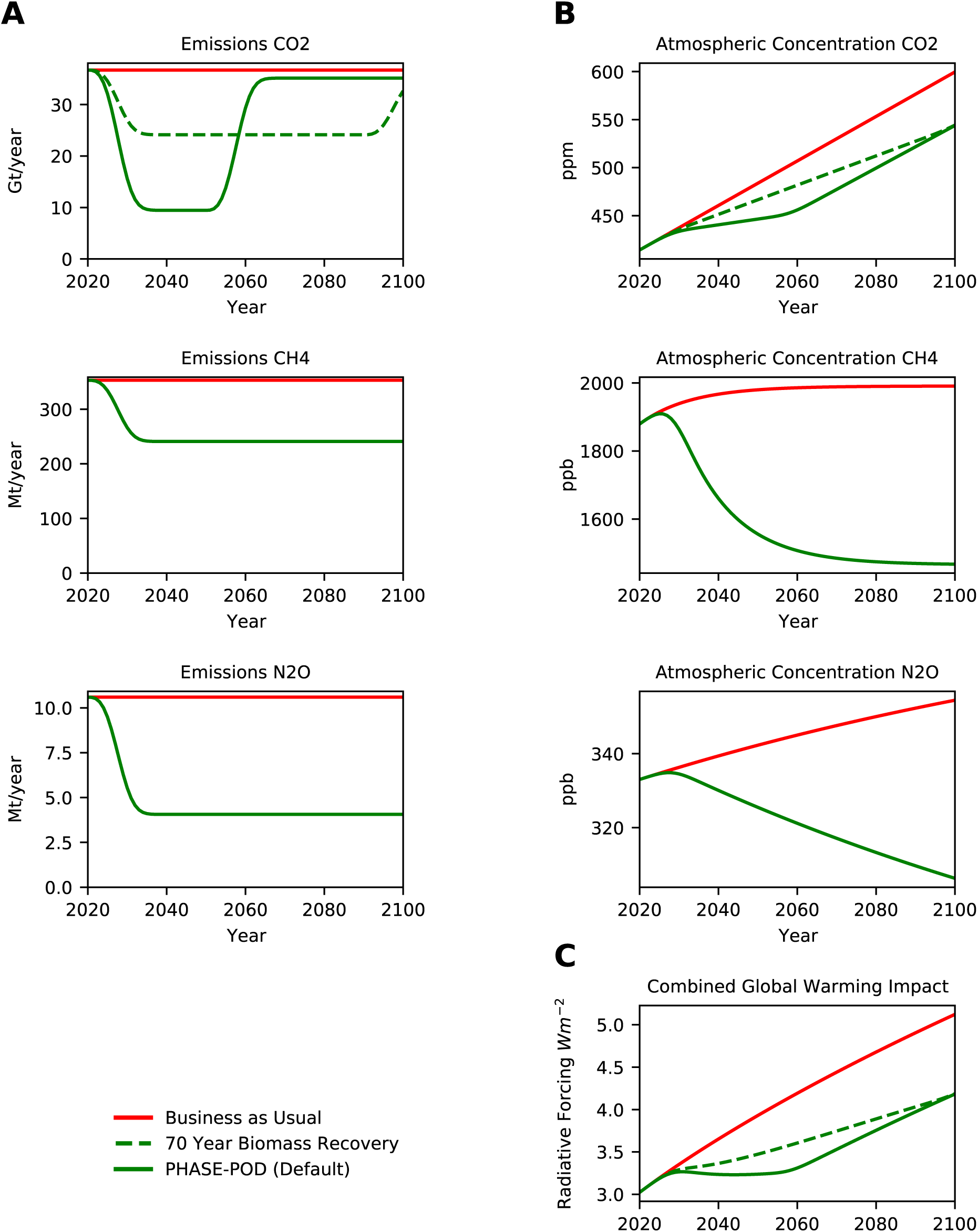
30yr vs. 70yr Biomass Recovery. (A) Projected annual emissions of *CO*_2_, *CH*_4_ and *N*_2_*O* for each scenarios. (B) Projected atmospheric concentrations of *CO*_2_, *CH*_4_ and *N*_2_*O* under each emission scenario. (C) Radiative Forcing (RF) inferred from atmospheric concentrations in (B) by formula of (Myhre et al., 1998; Ramaswamy et al., 2001) as modified in MAGICC6 (Meinshausen et al., 2011). Only differences between PHASE-POD default assumptions (15yr phaseout, 30yr carbon recovery, 100% carbon recovery, BAU non-agriculture emissions, FAO crop replacement, and FAO animal ag emissions) are given.

**Figure 2-S5.**
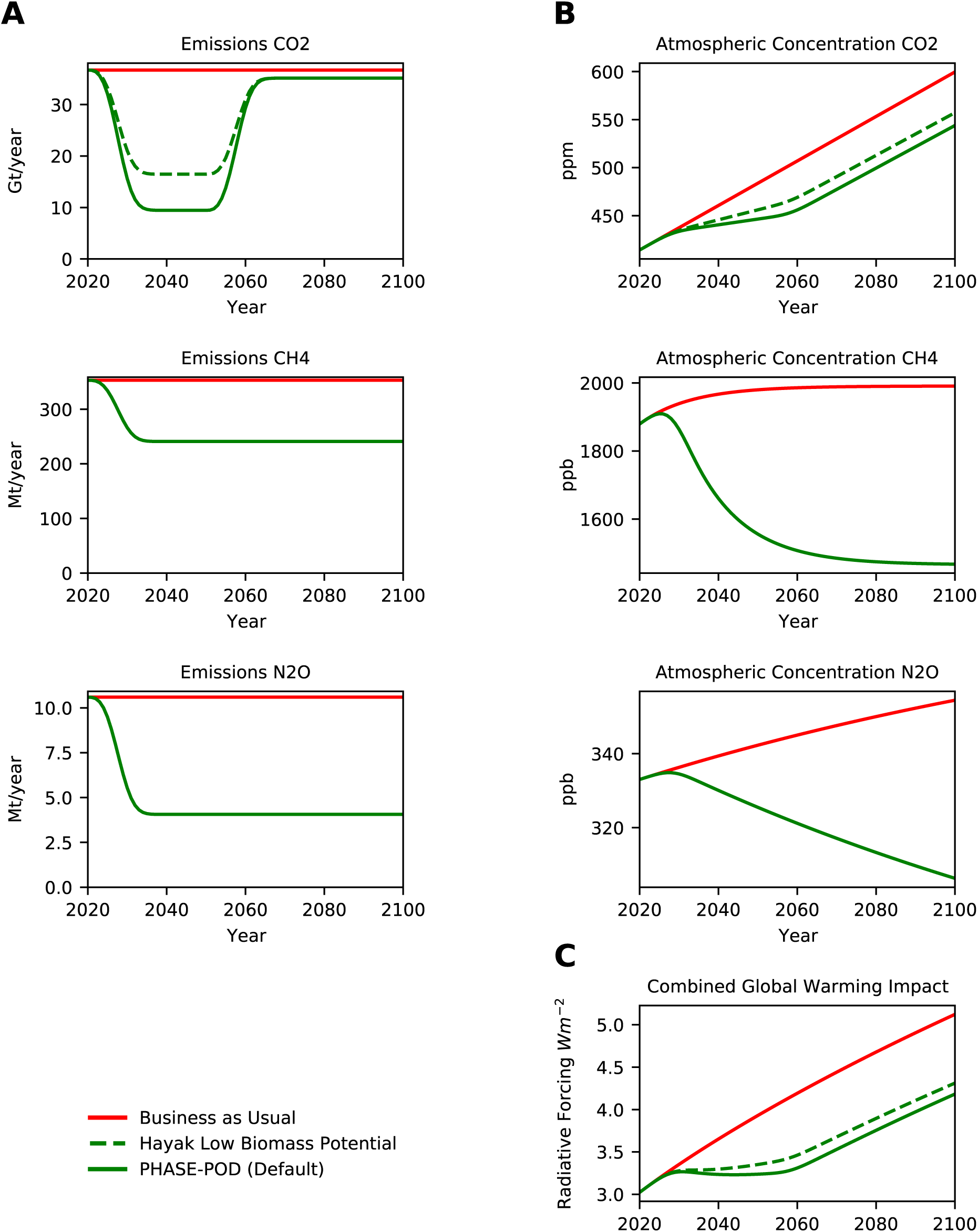
Hayek Median vs. Low Recovery Potential. (A) Projected annual emissions of *CO*_2_, *CH*_4_ and *N*2*O* for each scenarios. (B) Projected atmospheric concentrations of *CO*_2_, *CH*_4_ and *N*_2_*O* under each emission scenario. (C) Radiative Forcing (RF) inferred from atmospheric concentrations in (B) by formula of (Myhre et al., 1998; Ramaswamy et al., 2001) as modified in MAGICC6 (Meinshausen et al., 2011). Only differences between PHASE-POD default assumptions (15yr phaseout, 30yr carbon recovery, 100% carbon recovery, BAU non-agriculture emissions, FAO crop replacement, and FAO animal ag emissions) are given.

**Figure 2-S6.**
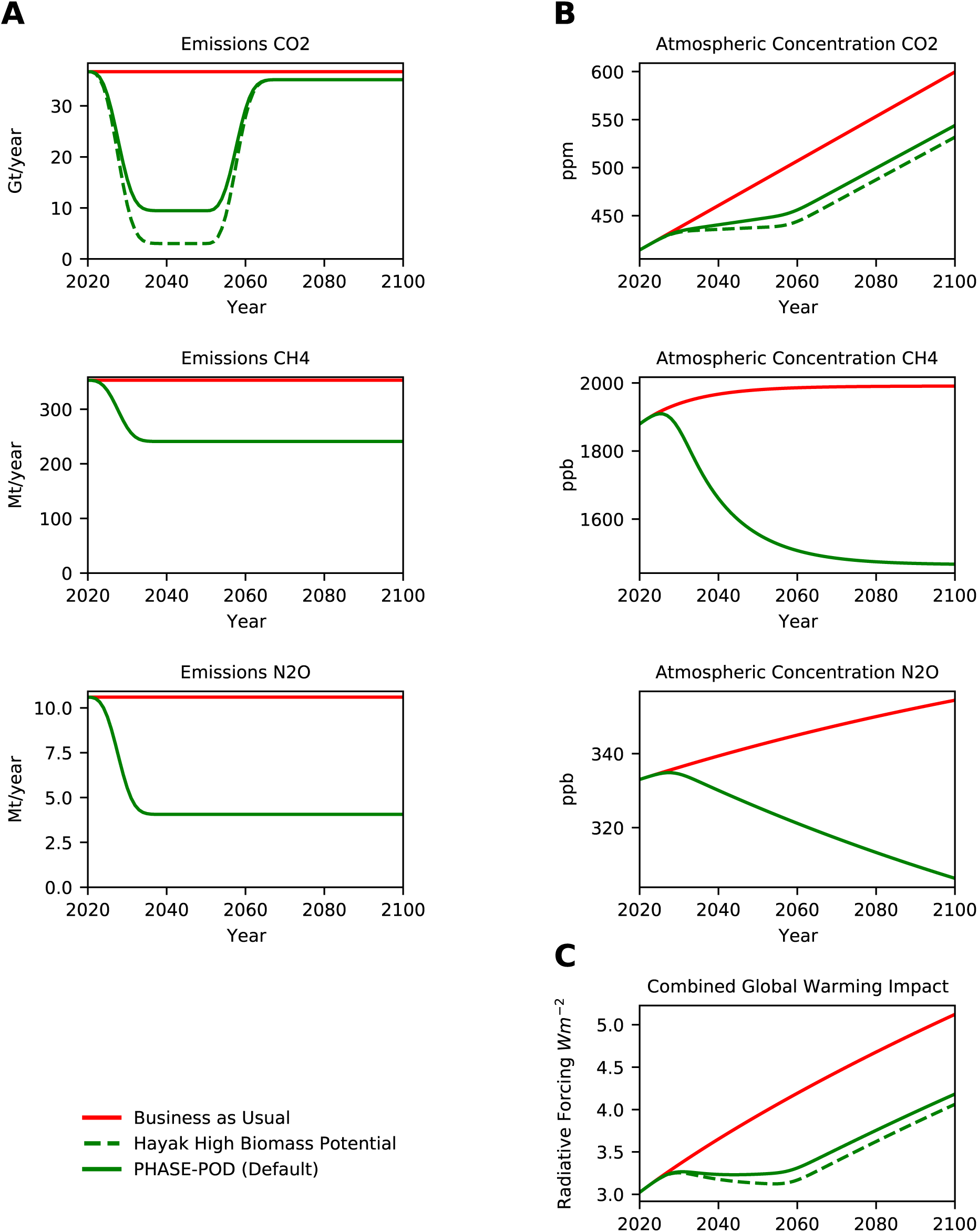
Hayek Median vs. High Recovery Potential. (A) Projected annual emissions of *CO*_2_, *CH*_4_ and *N*_2_*O* for each scenarios. (B) Projected atmospheric concentrations of *CO*_2_, *CH*_4_ and *N*_2_*O* under each emission scenario. (C) Radiative Forcing (RF) inferred from atmospheric concentrations in (B) by formula of (Myhre et al., 1998; Ramaswamy et al., 2001) as modified in MAGICC6 (Meinshausen et al., 2011). Only differences between PHASE-POD default assumptions (15yr phaseout, 30yr carbon recovery, 100% carbon recovery, BAU non-agriculture emissions, FAO crop replacement, and FAO animal ag emissions) are given.

**Figure 2-S7.**
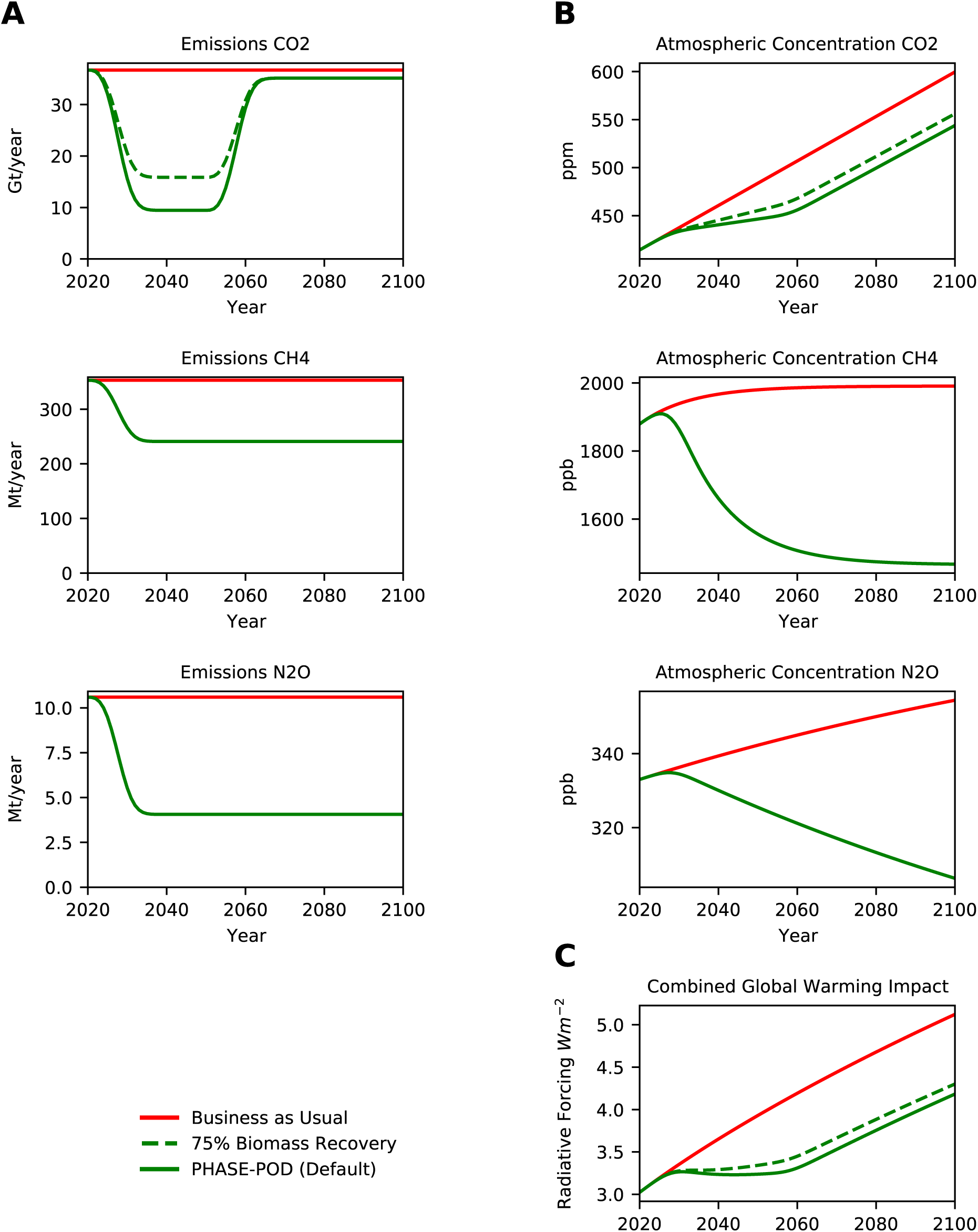
100% vs. 75% Biomass Recovery. (A) Projected annual emissions of *CO*_2_, *CH*_4_ and *N*_2_*O* for each scenarios. (B) Projected atmospheric concentrations of *CO*_2_, *CH*_4_ and *N*_2_*O* under each emission scenario. (C) Radiative Forcing (RF) inferred from atmospheric concentrations in (B) by formula of (Myhre et al., 1998; Ramaswamy et al., 2001) as modified in MAGICC6 (Meinshausen et al., 2011). Only differences between PHASE-POD default assumptions (15yr phaseout, 30yr carbon recovery, 100% carbon recovery, BAU non-agriculture emissions, FAO crop replacement, and FAO animal ag emissions) are given.

**Figure 2-S8.**
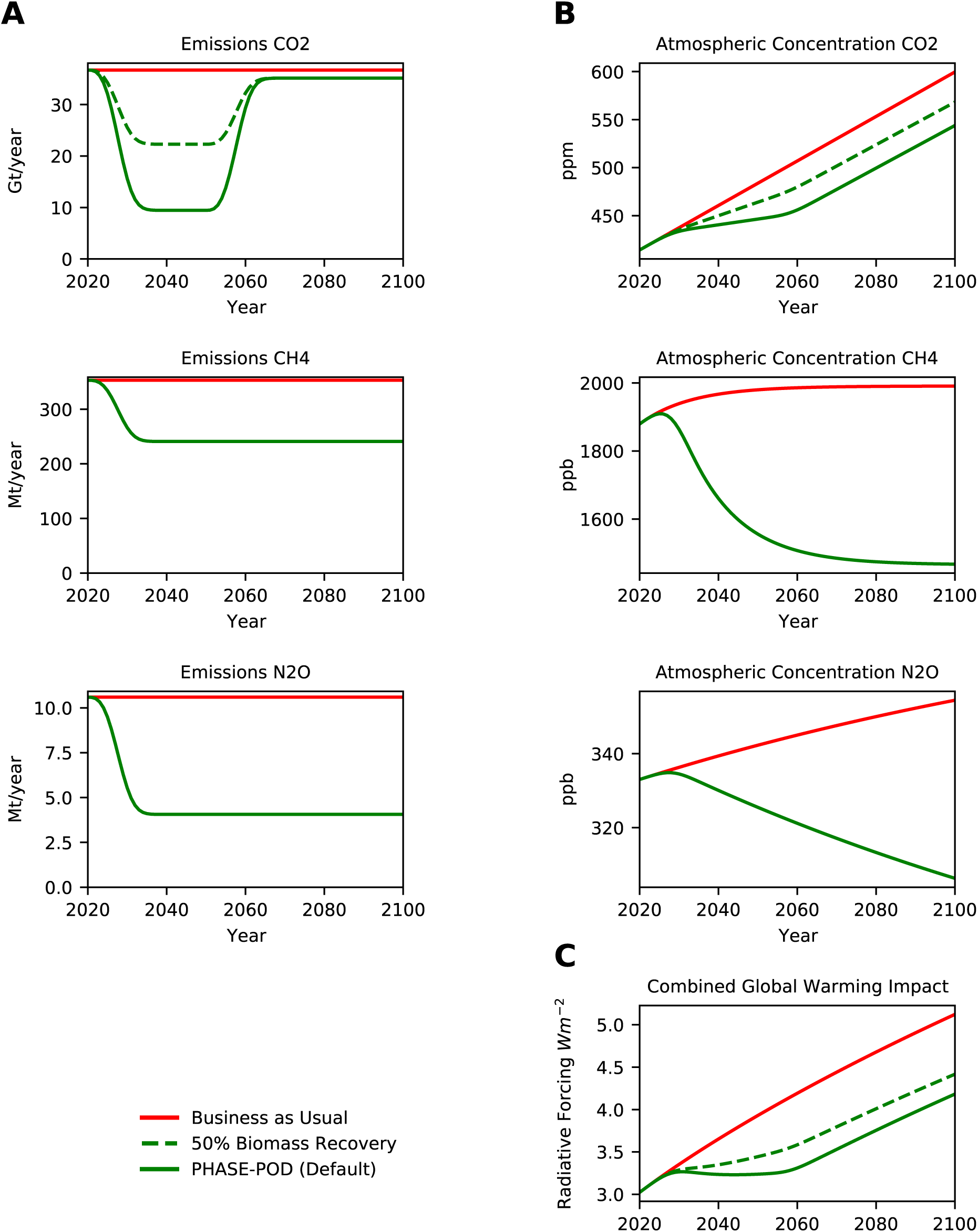
100% vs. 50% Biomass Recovery. (A) Projected annual emissions of *CO*_2_, *CH*_4_ and *N*_2_*O* for each scenarios. (B) Projected atmospheric concentrations of *CO*_2_, *CH*_4_ and *N*_2_*O* under each emission scenario. (C) Radiative Forcing (RF) inferred from atmospheric concentrations in (B) by formula of (Myhre et al., 1998; Ramaswamy et al., 2001) as modified in MAGICC6 (Meinshausen et al., 2011). Only differences between PHASE-POD default assumptions (15yr phaseout, 30yr carbon recovery, 100% carbon recovery, BAU non-agriculture emissions, FAO crop replacement, and FAO animal ag emissions) are given.

**Figure 2-S9.**
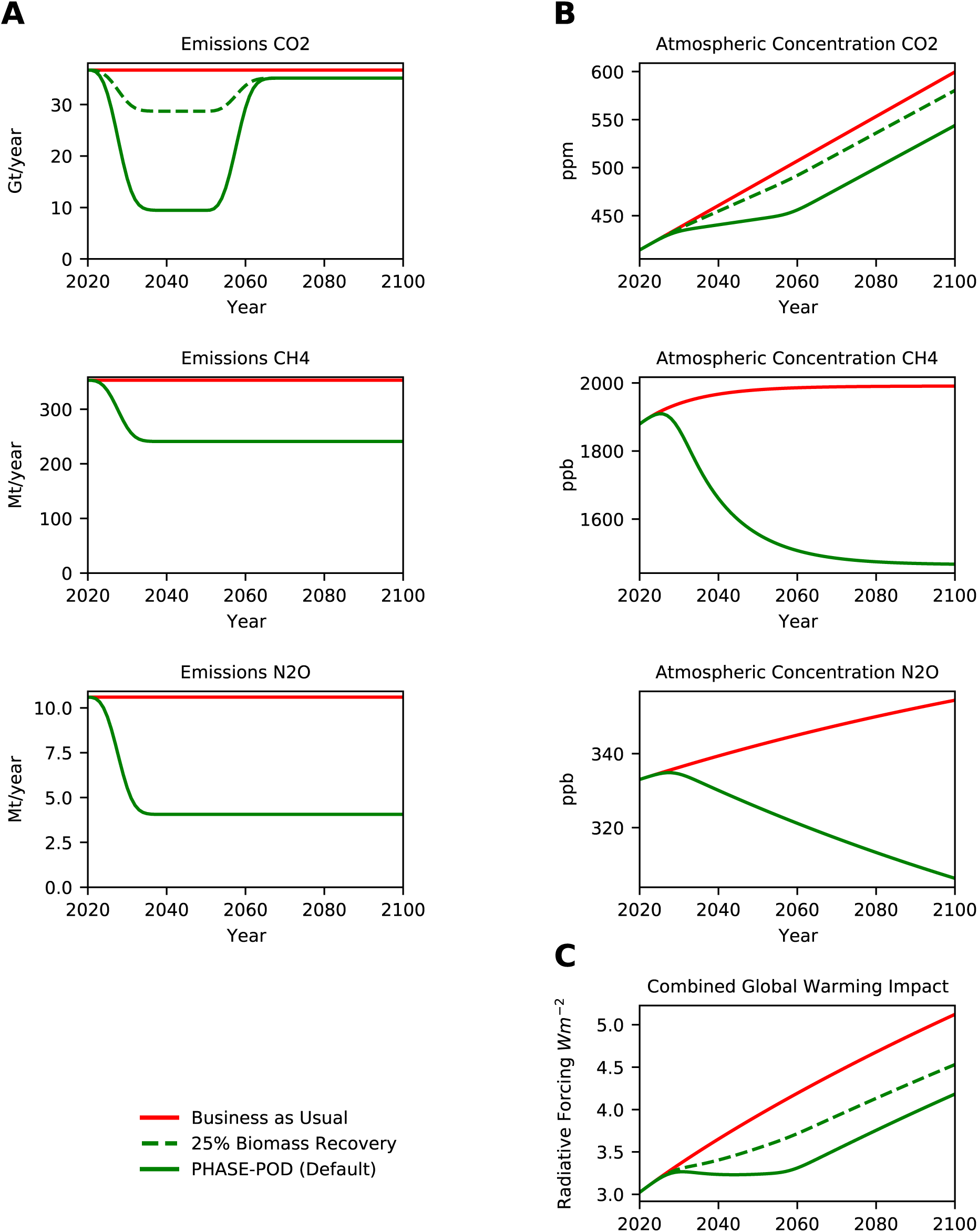
100% vs. 25% Biomass Recovery. (A) Projected annual emissions of *CO*_2_, *CH*_4_ and *N*_2_*O* for each scenarios. (B) Projected atmospheric concentrations of *CO*_2_, *CH*_4_ and *N*_2_*O* under each emission scenario. (C) Radiative Forcing (RF) inferred from atmospheric concentrations in (B) by formula of (Myhre et al., 1998; Ramaswamy et al., 2001) as modified in MAGICC6 (Meinshausen et al., 2011). Only differences between PHASE-POD default assumptions (15yr phaseout, 30yr carbon recovery, 100% carbon recovery, BAU non-agriculture emissions, FAO crop replacement, and FAO animal ag emissions) are given.

**Figure 2-S10.**
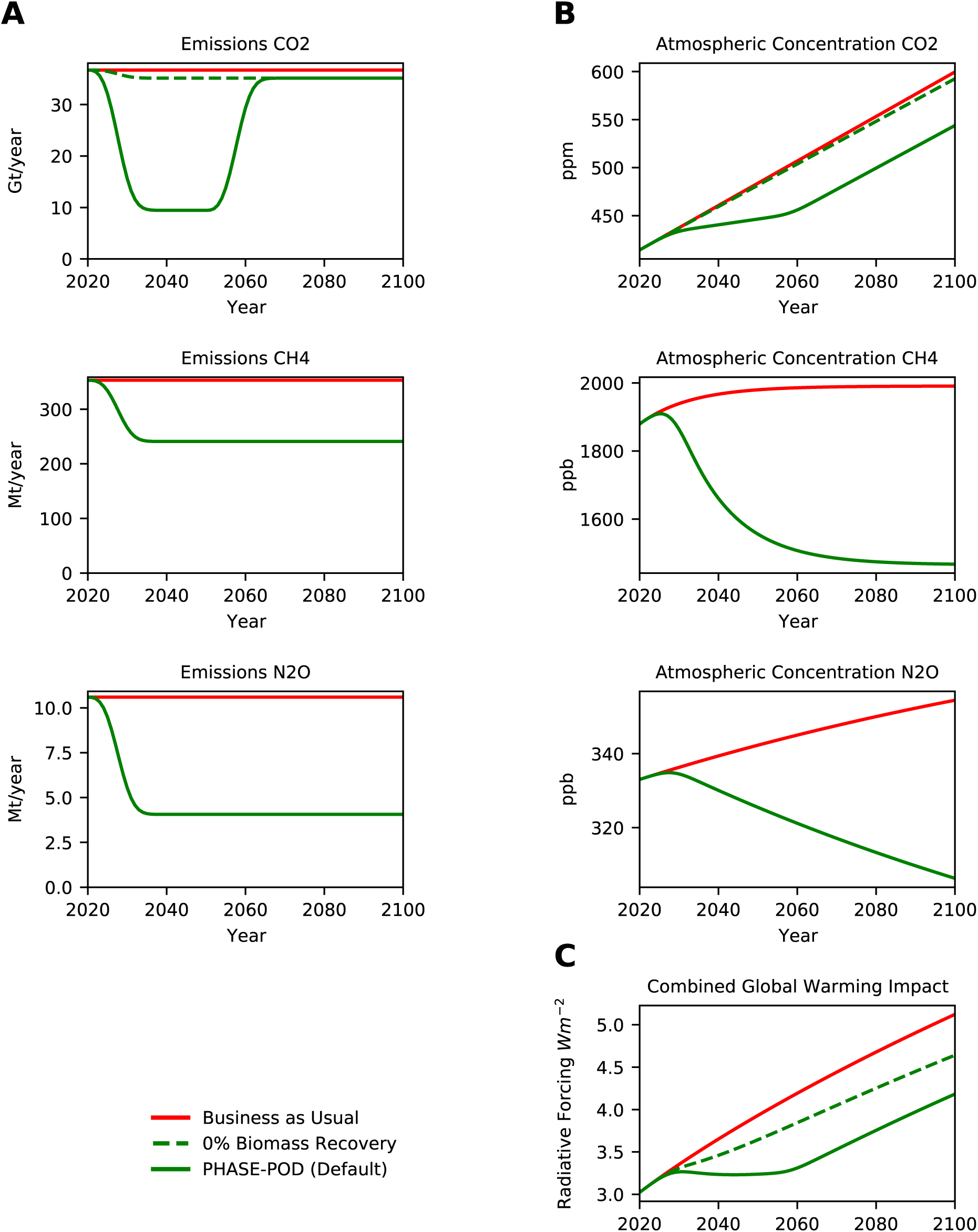
100% vs. 0% Biomass Recovery. (A) Projected annual emissions of *CO*_2_, *CH*_4_ and *N*_2_*O* for each scenarios. (B) Projected atmospheric concentrations of *CO*_2_, *CH*_4_ and *N*_2_*O* under each emission scenario. (C) Radiative Forcing (RF) inferred from atmospheric concentrations in (B) by formula of (Myhre et al., 1998; Ramaswamy et al., 2001) as modified in MAGICC6 (Meinshausen et al., 2011). Only differences between PHASE-POD default assumptions (15yr phaseout, 30yr carbon recovery, 100% carbon recovery, BAU non-agriculture emissions, FAO crop replacement, and FAO animal ag emissions) are given.

**Figure 2-S11.**
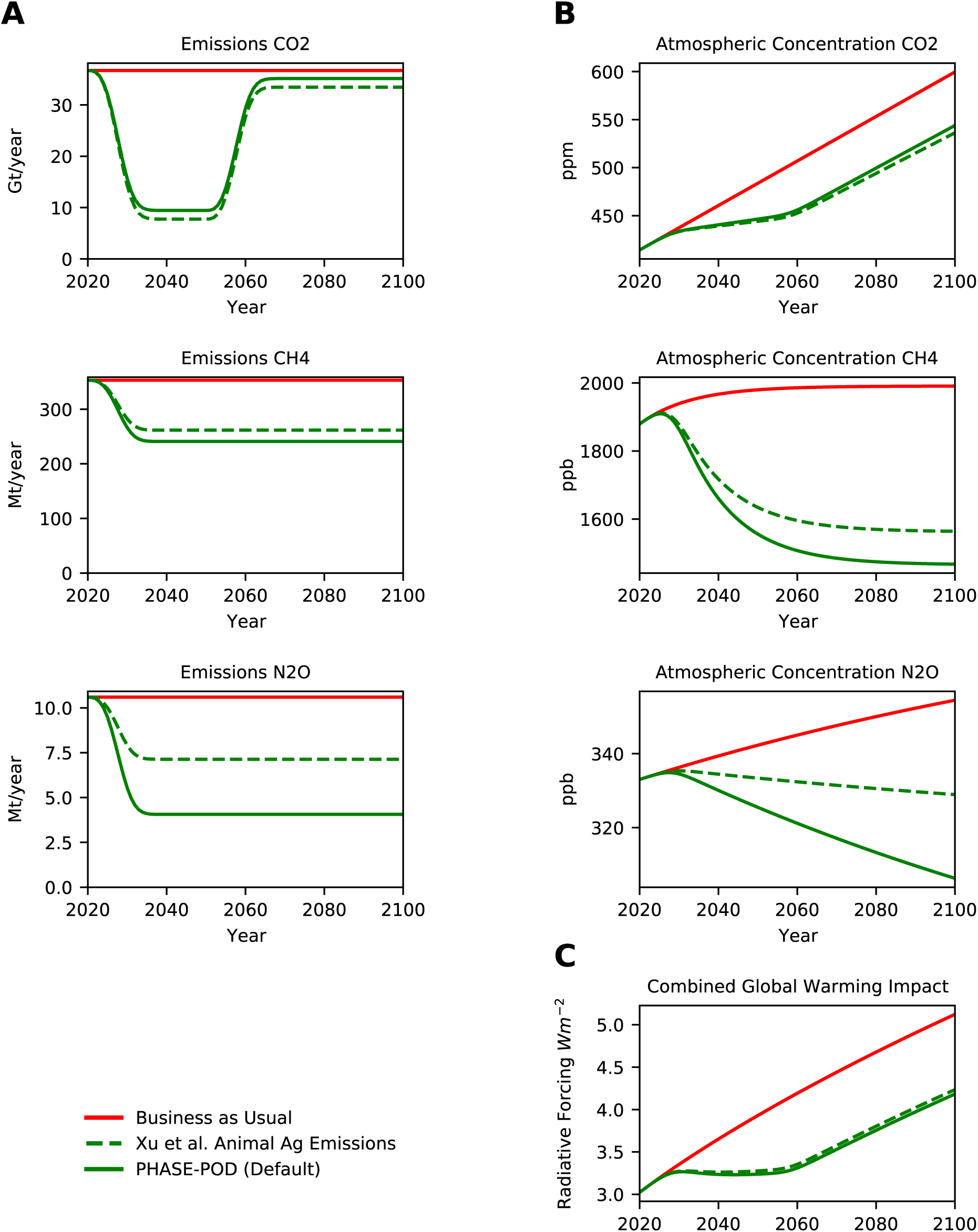
FAO vs. Xu Emissions. (A) Projected annual emissions of *CO*_2_, *CH*_4_ and *N*_2_*O* for each scenarios. (B) Projected atmospheric concentrations of *CO*_2_, *CH*_4_ and *N*_2_*O* under each emission scenario. (C) Radiative Forcing (RF) inferred from atmospheric concentrations in (B) by formula of (Myhre et al., 1998; Ramaswamy et al., 2001) as modified in MAGICC6 (Meinshausen et al., 2011). Only differences between PHASE-POD default assumptions (15yr phaseout, 30yr carbon recovery, 100% carbon recovery, BAU non-agriculture emissions, FAO crop replacement, and FAO animal ag emissions) are given.

**Figure 2-S12.**
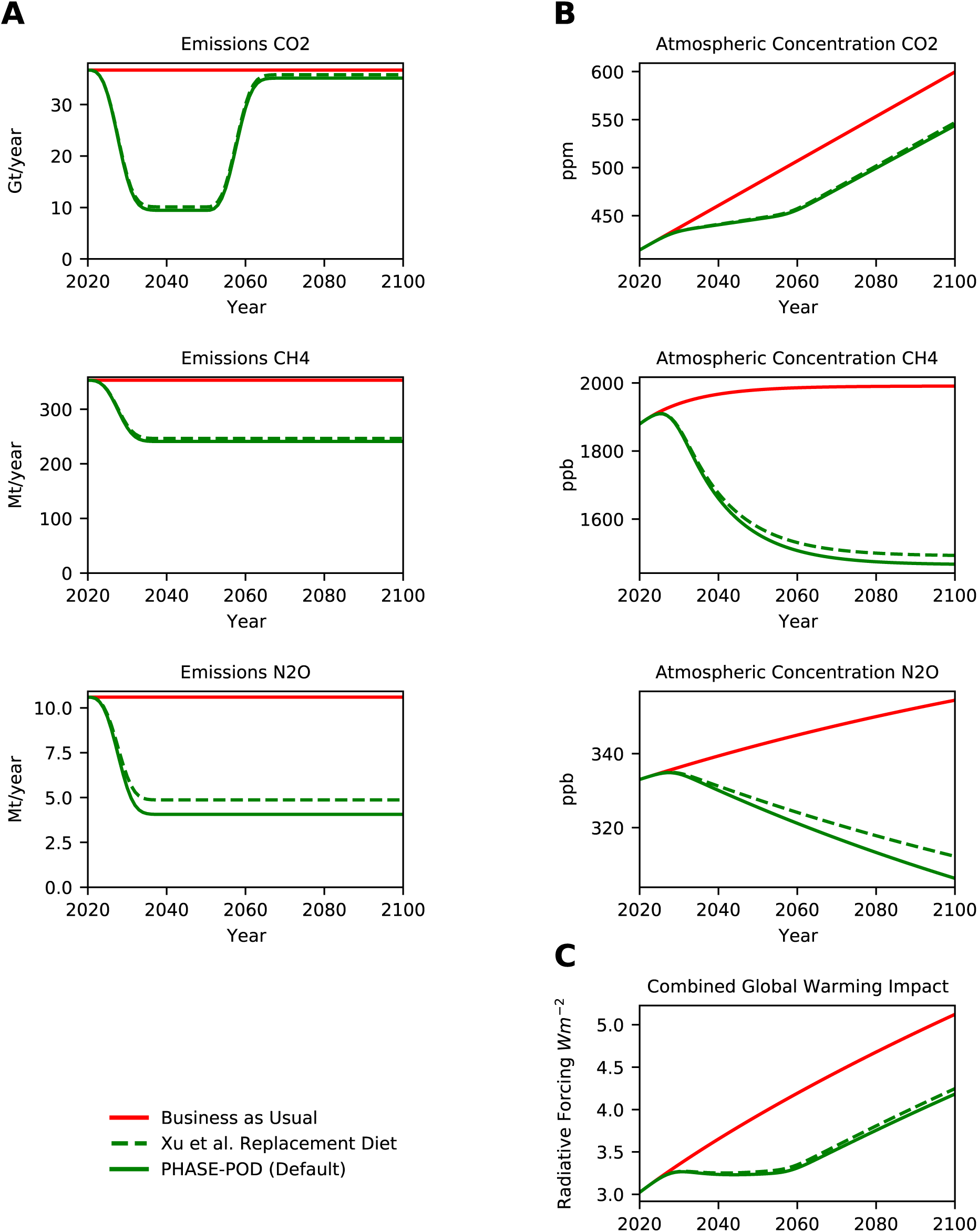
FAO vs. Xu Replacement Diet. (A) Projected annual emissions of *CO*_2_, *CH*_4_ and *N*_2_*O* for each scenarios. (B) Projected atmospheric concentrations of *CO*_2_, *CH*_4_ and *N*_2_*O* under each emission scenario. (C) Radiative Forcing (RF) inferred from atmospheric concentrations in (B) by formula of (Myhre et al., 1998; Ramaswamy et al., 2001) as modified in MAGICC6 (Meinshausen et al., 2011). Only differences between PHASE-POD default assumptions (15yr phaseout, 30yr carbon recovery, 100% carbon recovery, BAU non-agriculture emissions, FAO crop replacement, and FAO animal ag emissions) are given.

**Figure 2-S13.**
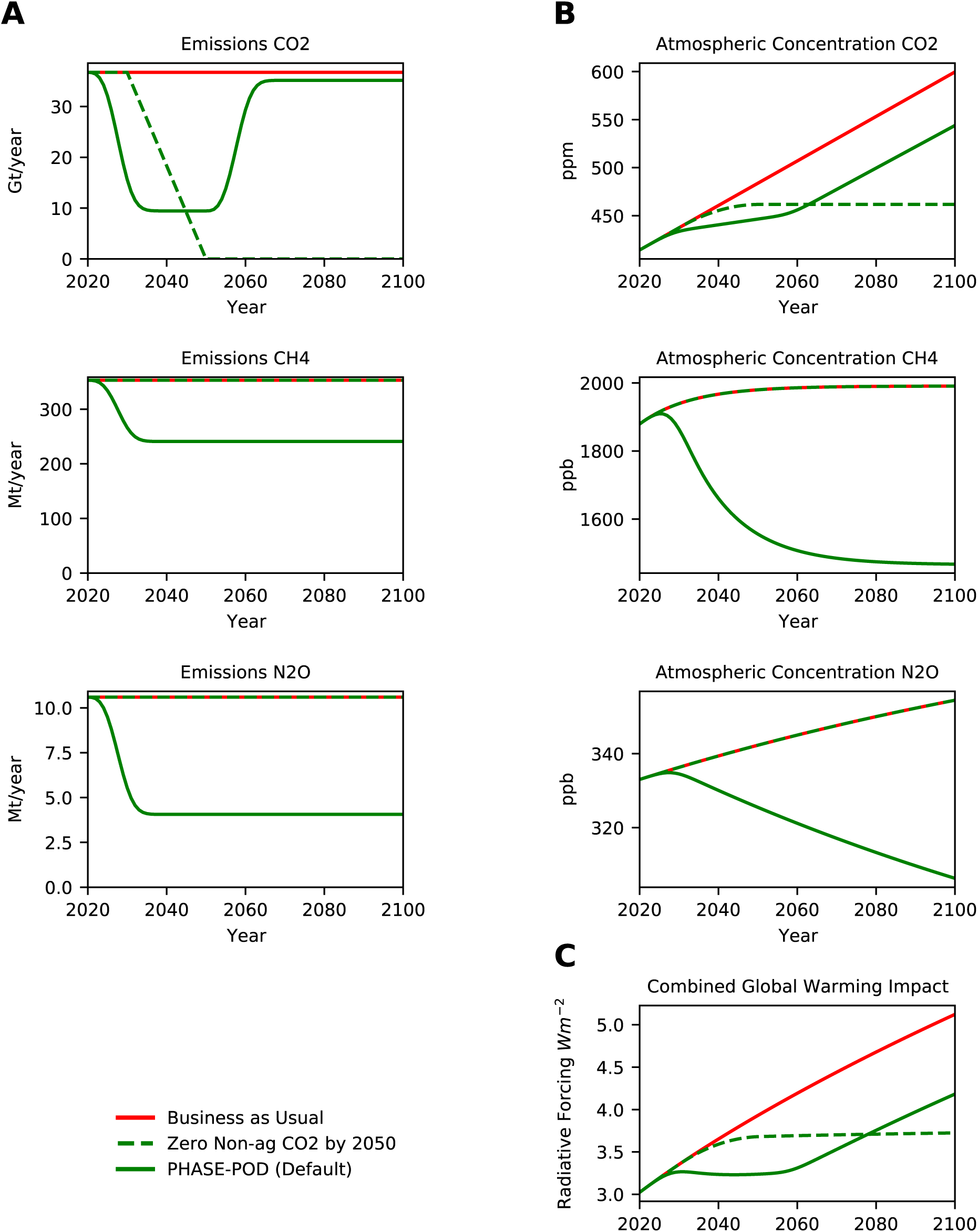
15yr Phaseout vs. Net Zero CO2. (A) Projected annual emissions of *CO*_2_, *CH*_4_ and *N*_2_*O* for each scenarios. (B) Projected atmospheric concentrations of *CO*_2_, *CH*_4_ and *N*_2_*O* under each emission scenario. (C) Radiative Forcing (RF) inferred from atmospheric concentrations in (B) by formula of (Myhre et al., 1998; Ramaswamy et al., 2001) as modified in MAGICC6 (Meinshausen et al., 2011). Only differences between PHASE-POD default assumptions (15yr phaseout, 30yr carbon recovery, 100% carbon recovery, BAU non-agriculture emissions, FAO crop replacement, and FAO animal ag emissions) are given.

**Figure 2-S14.**
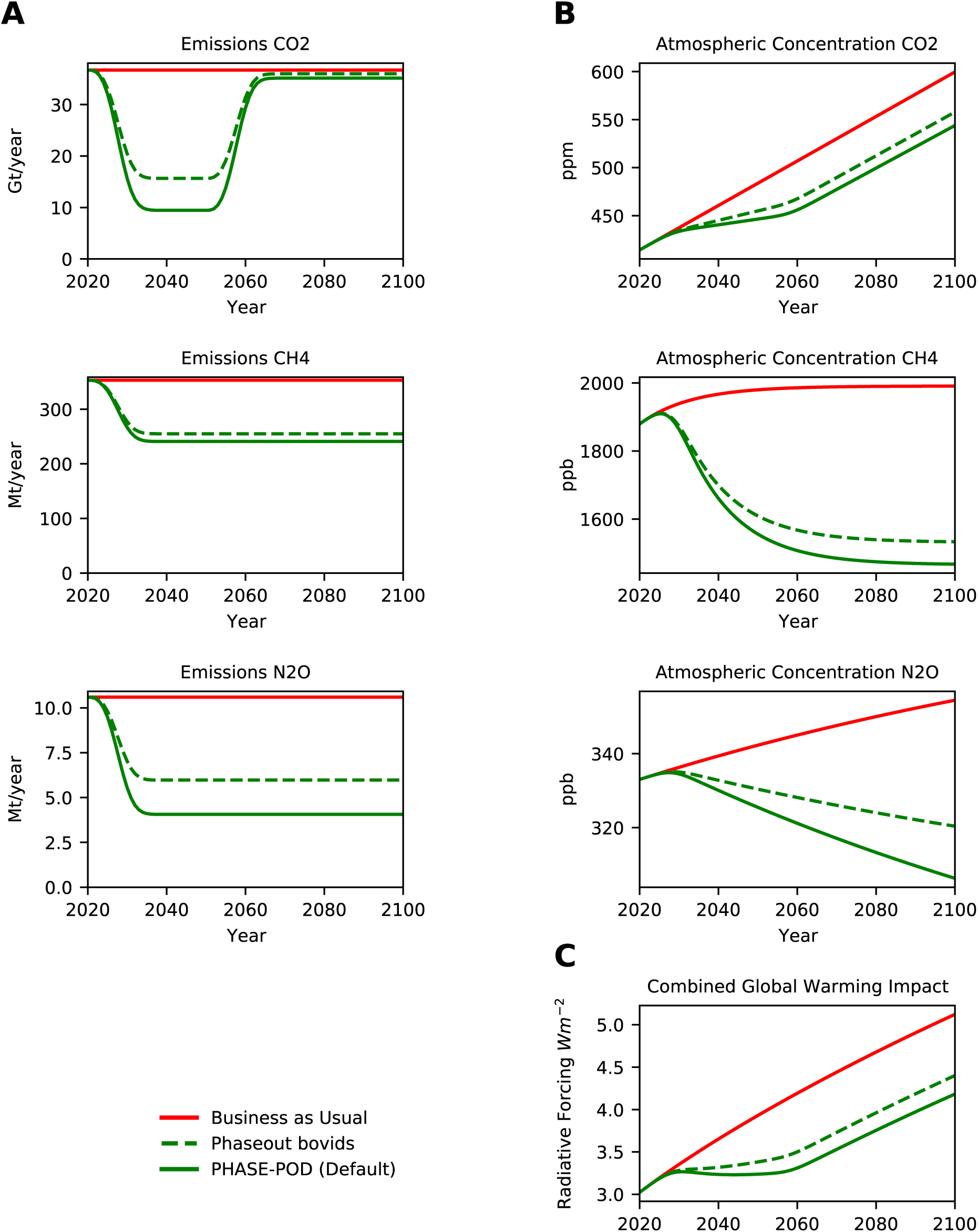
Phaseout of Bovids. (A) Projected annual emissions of *CO*_2_, *CH*_4_ and *N*_2_*O* for each scenarios. (B) Projected atmospheric concentrations of *CO*_2_, *CH*_4_ and *N*_2_*O* under each emission scenario. (C) Radiative Forcing (RF) inferred from atmospheric concentrations in (B) by formula of (Myhre et al., 1998; Ramaswamy et al., 2001) as modified in MAGICC6 (Meinshausen et al., 2011). Only differences between PHASE-POD default assumptions (15yr phaseout, 30yr carbon recovery, 100% carbon recovery, BAU non-agriculture emissions, FAO crop replacement, and FAO animal ag emissions) are given.

**Figure 2-S15.**
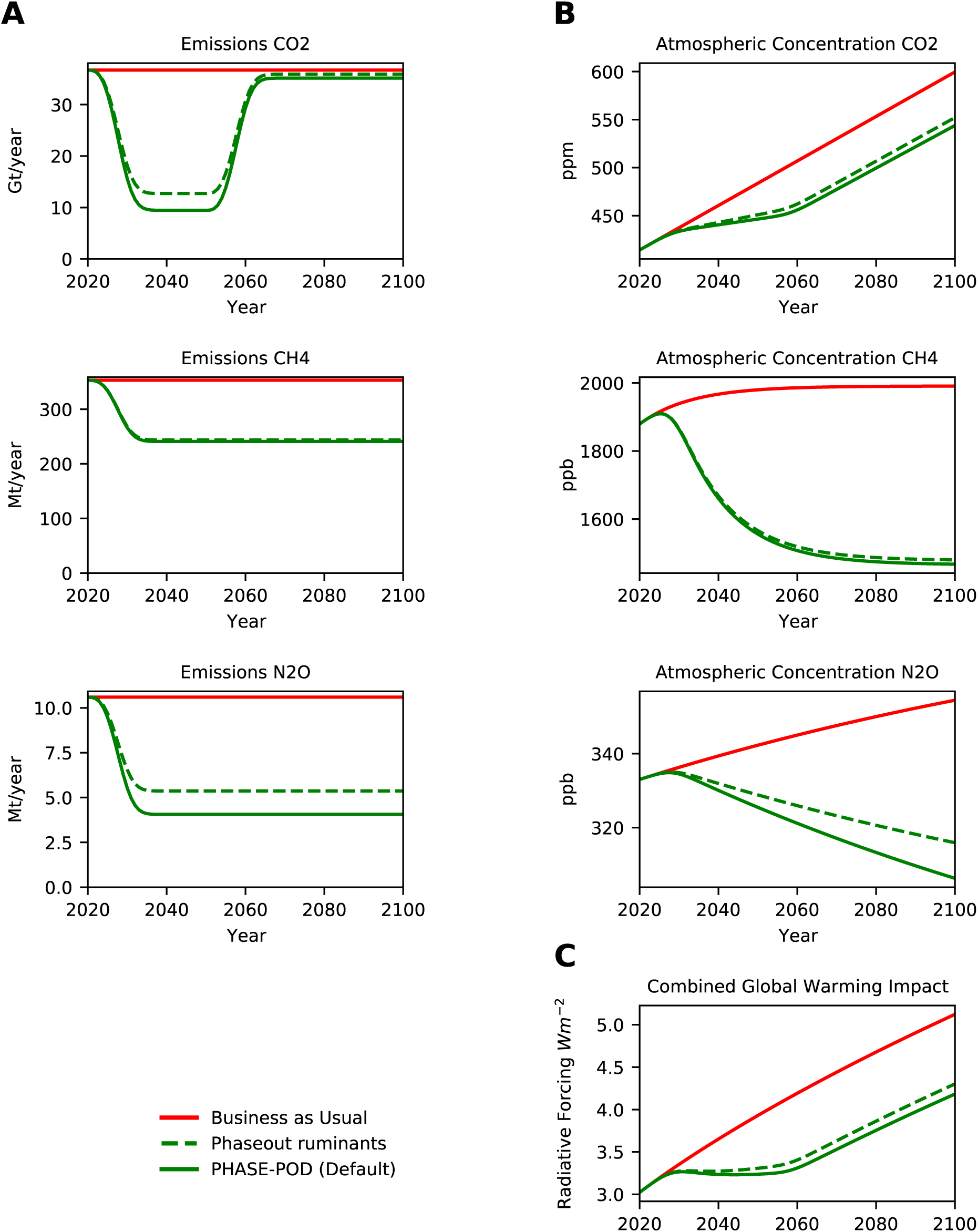
Phaseout of Ruminants. (A) Projected annual emissions of *CO*_2_, *CH*_4_ and *N*_2_*O* for each scenarios. (B) Projected atmospheric concentrations of *CO*_2_, *CH*_4_ and *N*_2_*O* under each emission scenario. (C) Radiative Forcing (RF) inferred from atmospheric concentrations in (B) by formula of (Myhre et al., 1998; Ramaswamy et al., 2001) as modified in MAGICC6 (Meinshausen et al., 2011). Only differences between PHASE-POD default assumptions (15yr phaseout, 30yr carbon recovery, 100% carbon recovery, BAU non-agriculture emissions, FAO crop replacement, and FAO animal ag emissions) are given.

**Figure 2-S16.**
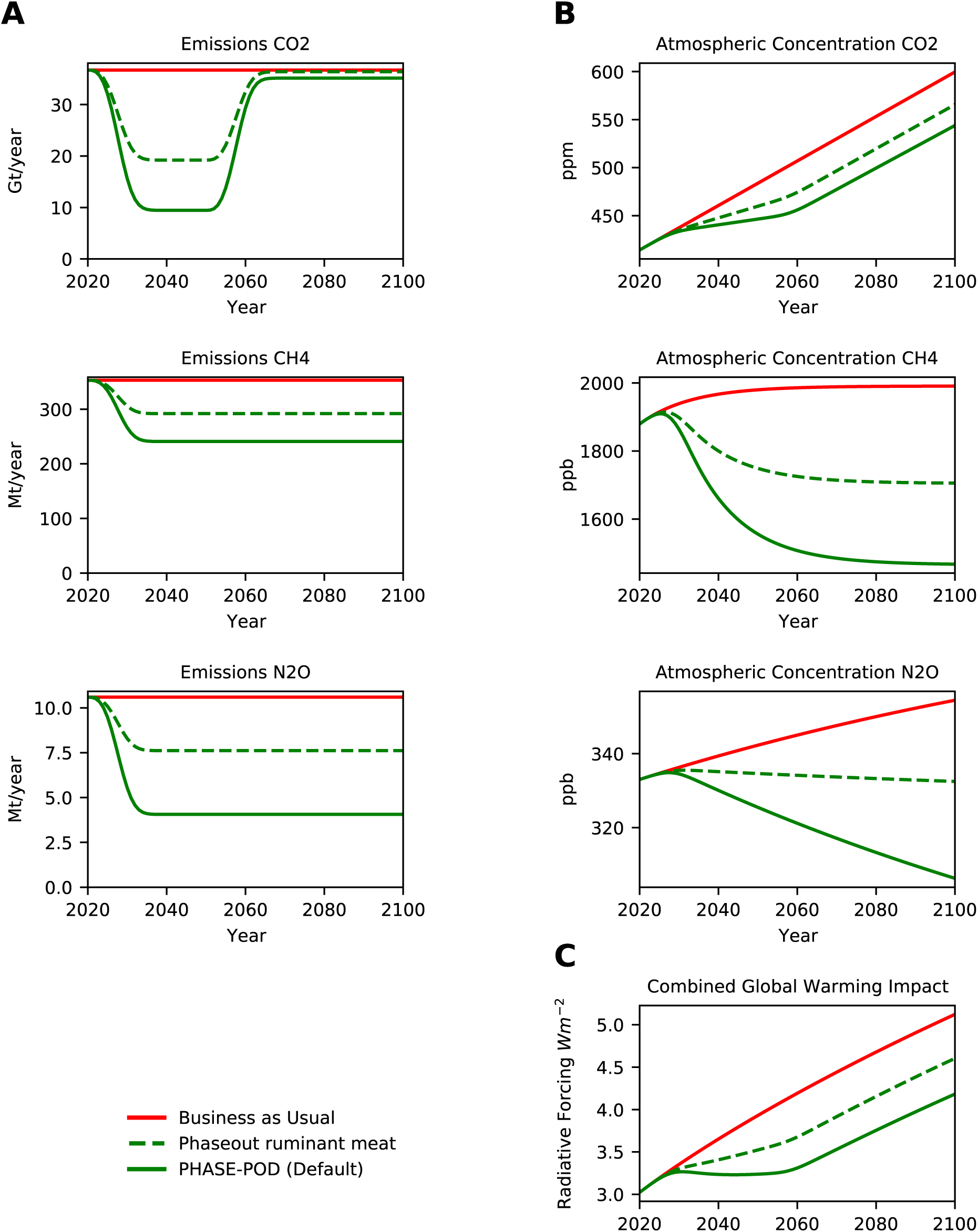
Phaseout of Ruminant Meat. (A) Projected annual emissions of *CO*_2_, *CH*_4_ and *N*_2_*O* for each scenarios. (B) Projected atmospheric concentrations of *CO*_2_, *CH*_4_ and *N*_2_*O* under each emission scenario. (C) Radiative Forcing (RF) inferred from atmospheric concentrations in (B) by formula of (Myhre et al., 1998; Ramaswamy et al., 2001) as modified in MAGICC6 (Meinshausen et al., 2011). Only differences between PHASE-POD default assumptions (15yr phaseout, 30yr carbon recovery, 100% carbon recovery, BAU non-agriculture emissions, FAO crop replacement, and FAO animal ag emissions) are given.

**Figure 2-S17.**
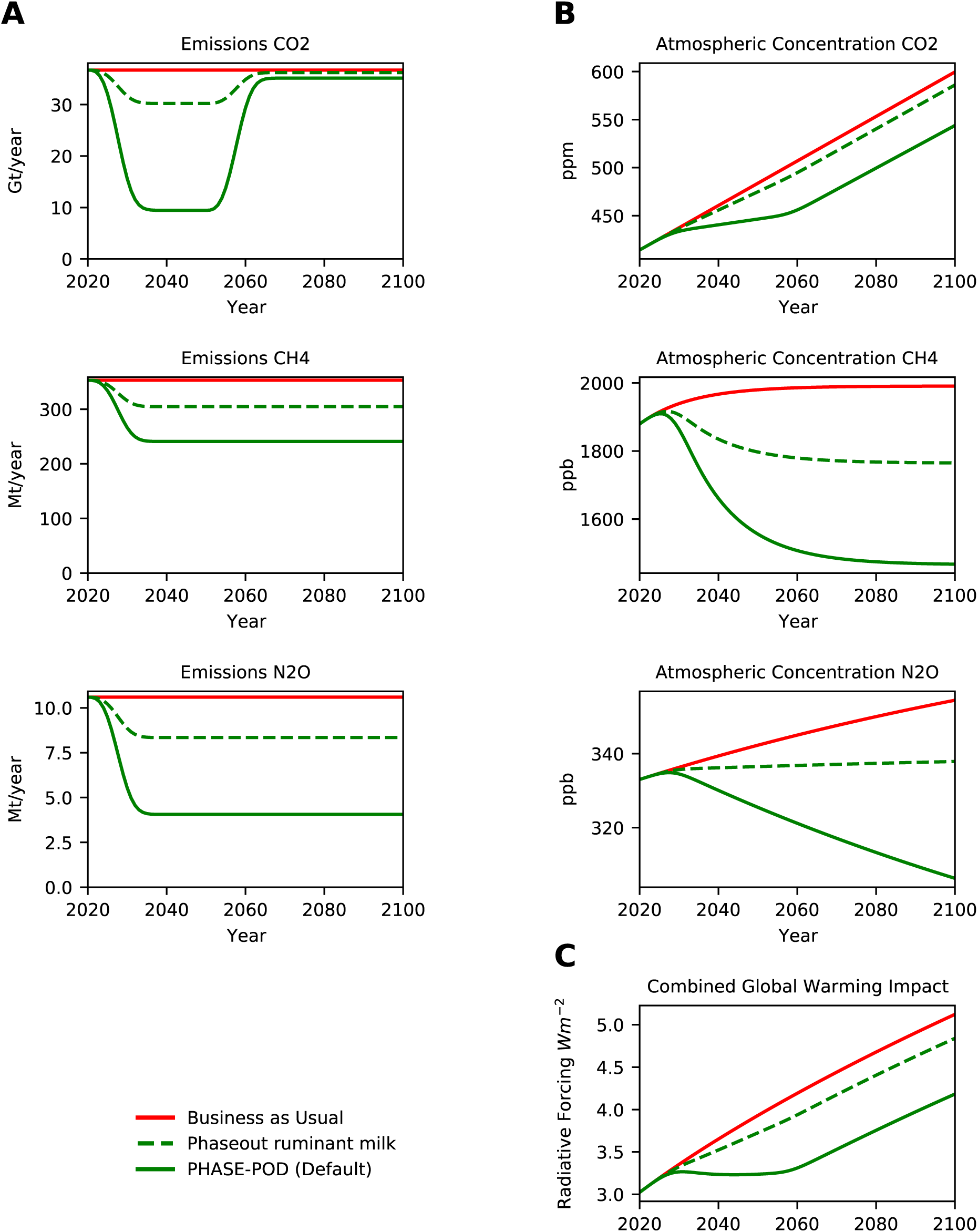
Phaseout of Ruminant Milk. (A) Projected annual emissions of *CO*_2_, *CH*_4_ and *N*_2_*O* for each scenarios. (B) Projected atmospheric concentrations of *CO*_2_, *CH*_4_ and *N*_2_*O* under each emission scenario. (C) Radiative Forcing (RF) inferred from atmospheric concentrations in (B) by formula of (Myhre et al., 1998; Ramaswamy et al., 2001) as modified in MAGICC6 (Meinshausen et al., 2011). Only differences between PHASE-POD default assumptions (15yr phaseout, 30yr carbon recovery, 100% carbon recovery, BAU non-agriculture emissions, FAO crop replacement, and FAO animal ag emissions) are given.

**Figure 2-S18.**
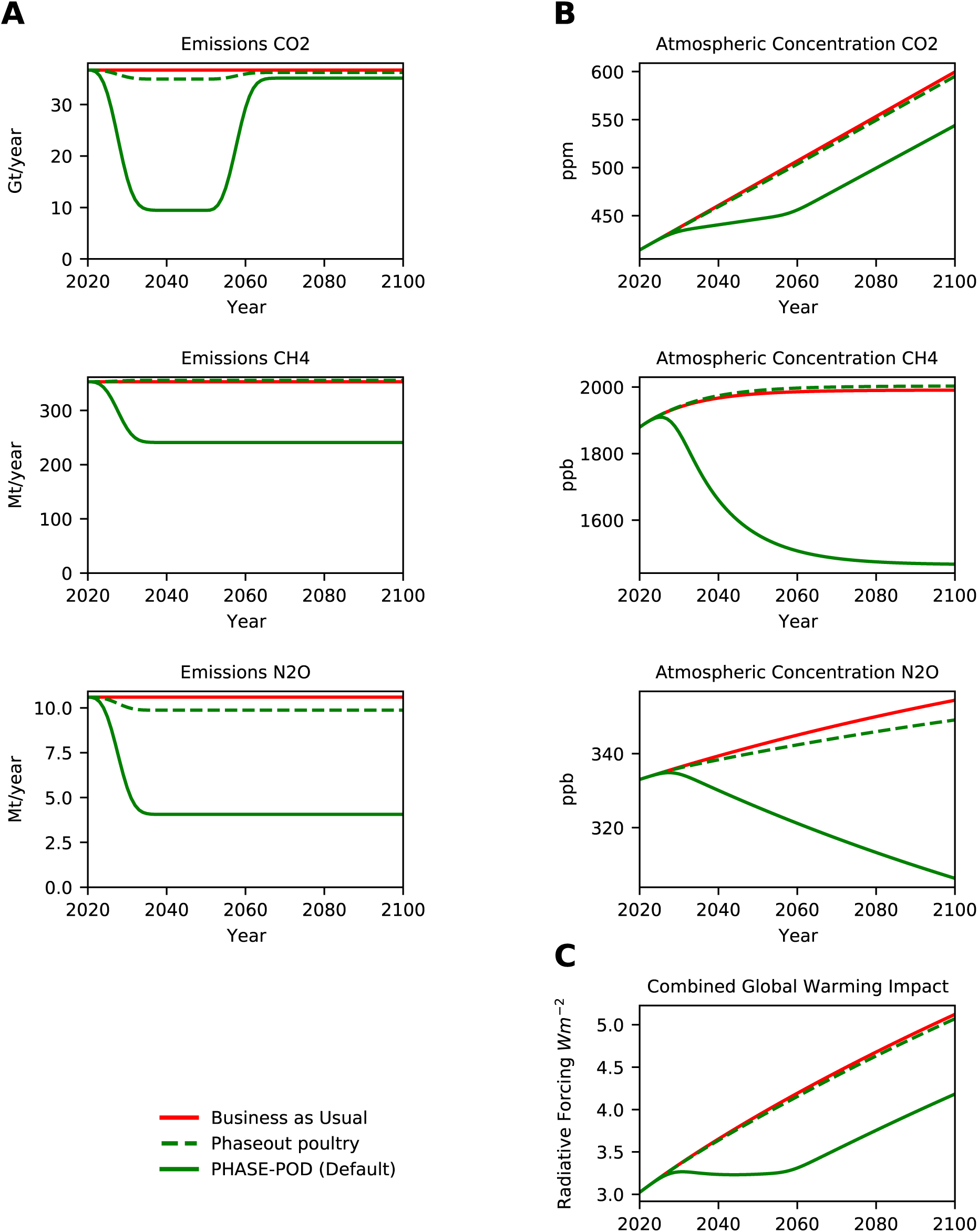
Phaseout of Poultry. (A) Projected annual emissions of *CO*_2_, *CH*_4_ and *N*_2_*O* for each scenarios. (B) Projected atmospheric concentrations of *CO*_2_, *CH*_4_ and *N*_2_*O* under each emission scenario. (C) Radiative Forcing (RF) inferred from atmospheric concentrations in (B) by formula of (Myhre et al., 1998; Ramaswamy et al., 2001) as modified in MAGICC6 (Meinshausen et al., 2011). Only differences between PHASE-POD default assumptions (15yr phaseout, 30yr carbon recovery, 100% carbon recovery, BAU non-agriculture emissions, FAO crop replacement, and FAO animal ag emissions) are given.

**Figure 2-S19.**
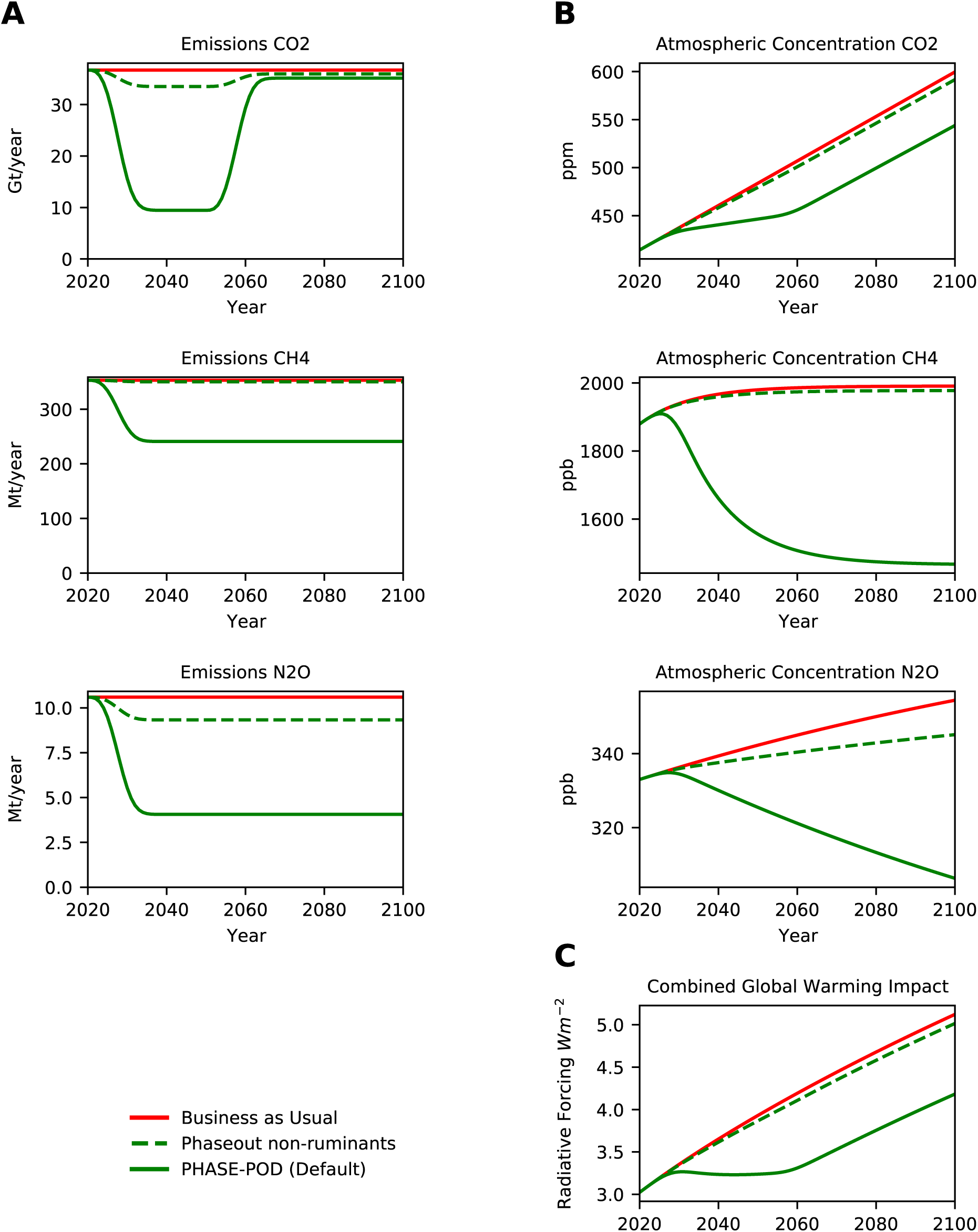
Phaseout of Non-Ruminants. (A) Projected annual emissions of *CO*_2_, *CH*_4_ and *N*_2_*O* for each scenarios. (B) Projected atmospheric concentrations of *CO*_2_, *CH*_4_ and *N*_2_*O* under each emission scenario. (C) Radiative Forcing (RF) inferred from atmospheric concentrations in (B) by formula of (Myhre et al., 1998; Ramaswamy et al., 2001) as modified in MAGICC6 (Meinshausen et al., 2011). Only differences between PHASE-POD default assumptions (15yr phaseout, 30yr carbon recovery, 100% carbon recovery, BAU non-agriculture emissions, FAO crop replacement, and FAO animal ag emissions) are given.

**Figure 2-S20.**
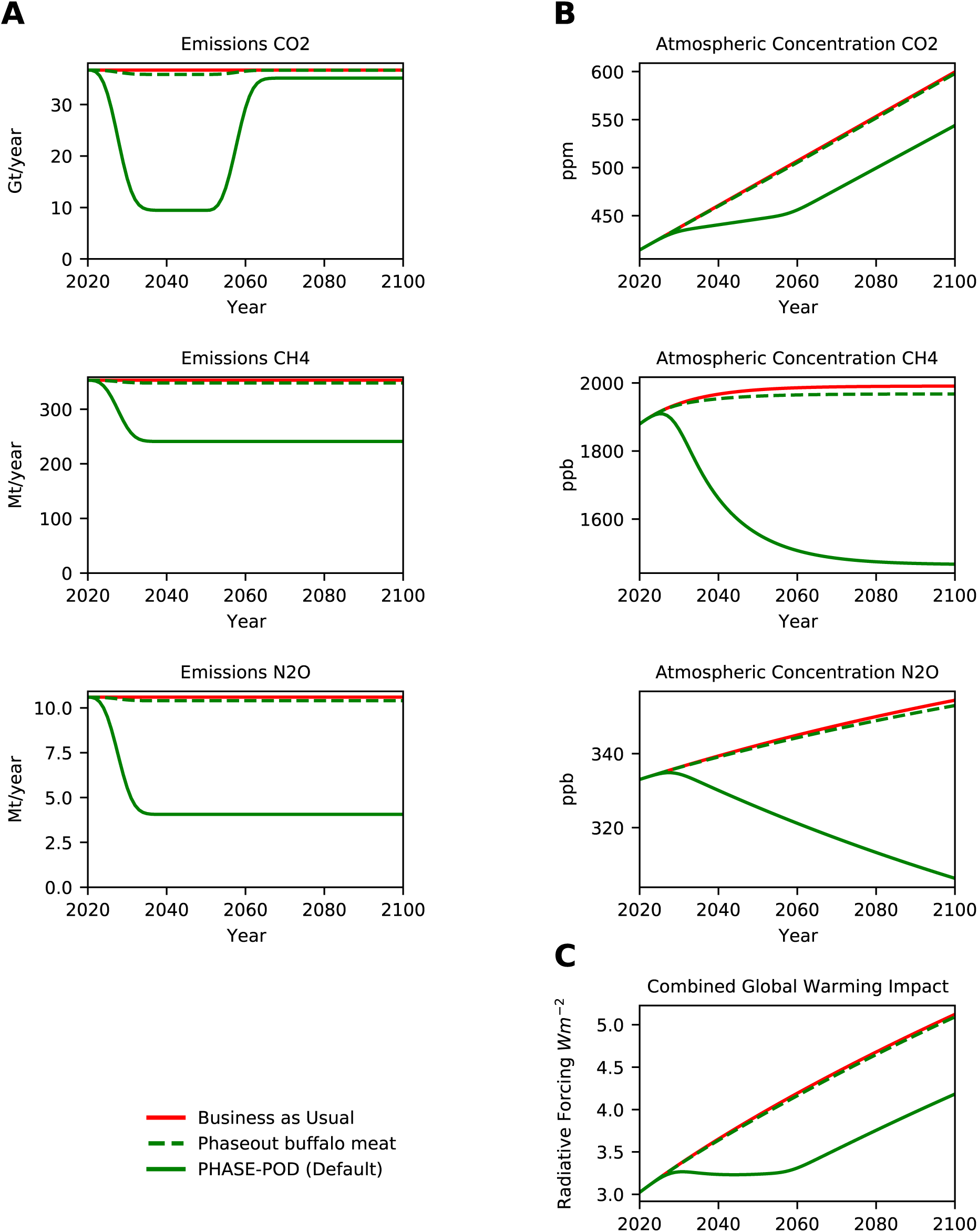
Phaseout of Buffalo Meat. (A) Projected annual emissions of *CO*_2_, *CH*_4_ and *N*_2_*O* for each scenarios. (B) Projected atmospheric concentrations of *CO*_2_, *CH*_4_ and *N*_2_*O* under each emission scenario. (C) Radiative Forcing (RF) inferred from atmospheric concentrations in (B) by formula of (Myhre et al., 1998; Ramaswamy et al., 2001) as modified in MAGICC6 (Meinshausen et al., 2011). Only differences between PHASE-POD default assumptions (15yr phaseout, 30yr carbon recovery, 100% carbon recovery, BAU non-agriculture emissions, FAO crop replacement, and FAO animal ag emissions) are given.

**Figure 2-S21.**
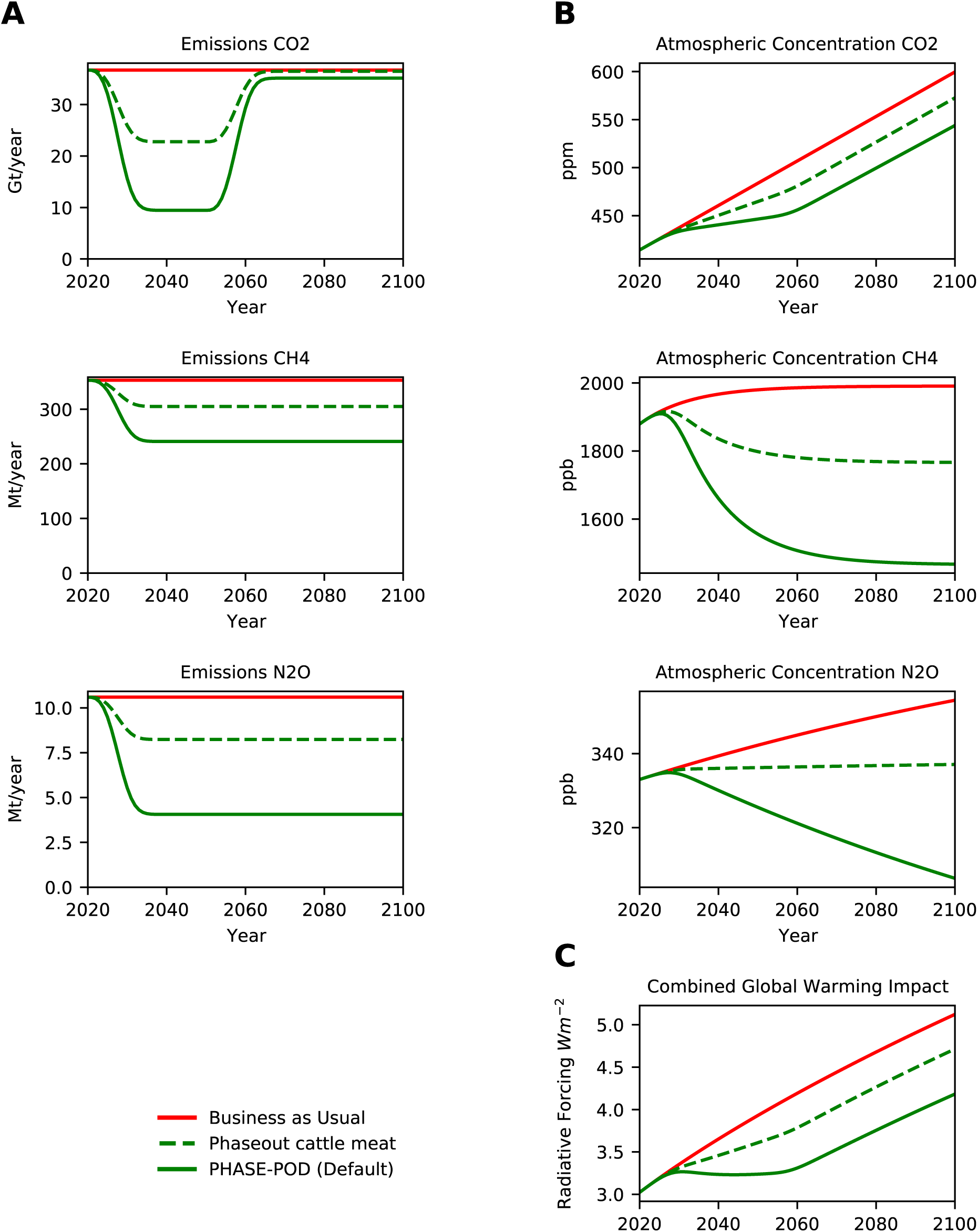
Phaseout of Cattle Meat. (A) Projected annual emissions of *CO*_2_, *CH*_4_ and *N*_2_*O* for each scenarios. (B) Projected atmospheric concentrations of *CO*_2_, *CH*_4_ and *N*_2_*O* under each emission scenario. (C) Radiative Forcing (RF) inferred from atmospheric concentrations in (B) by formula of (Myhre et al., 1998; Ramaswamy et al., 2001) as modified in MAGICC6 (Meinshausen et al., 2011). Only differences between PHASE-POD default assumptions (15yr phaseout, 30yr carbon recovery, 100% carbon recovery, BAU non-agriculture emissions, FAO crop replacement, and FAO animal ag emissions) are given.

**Figure 2-S22.**
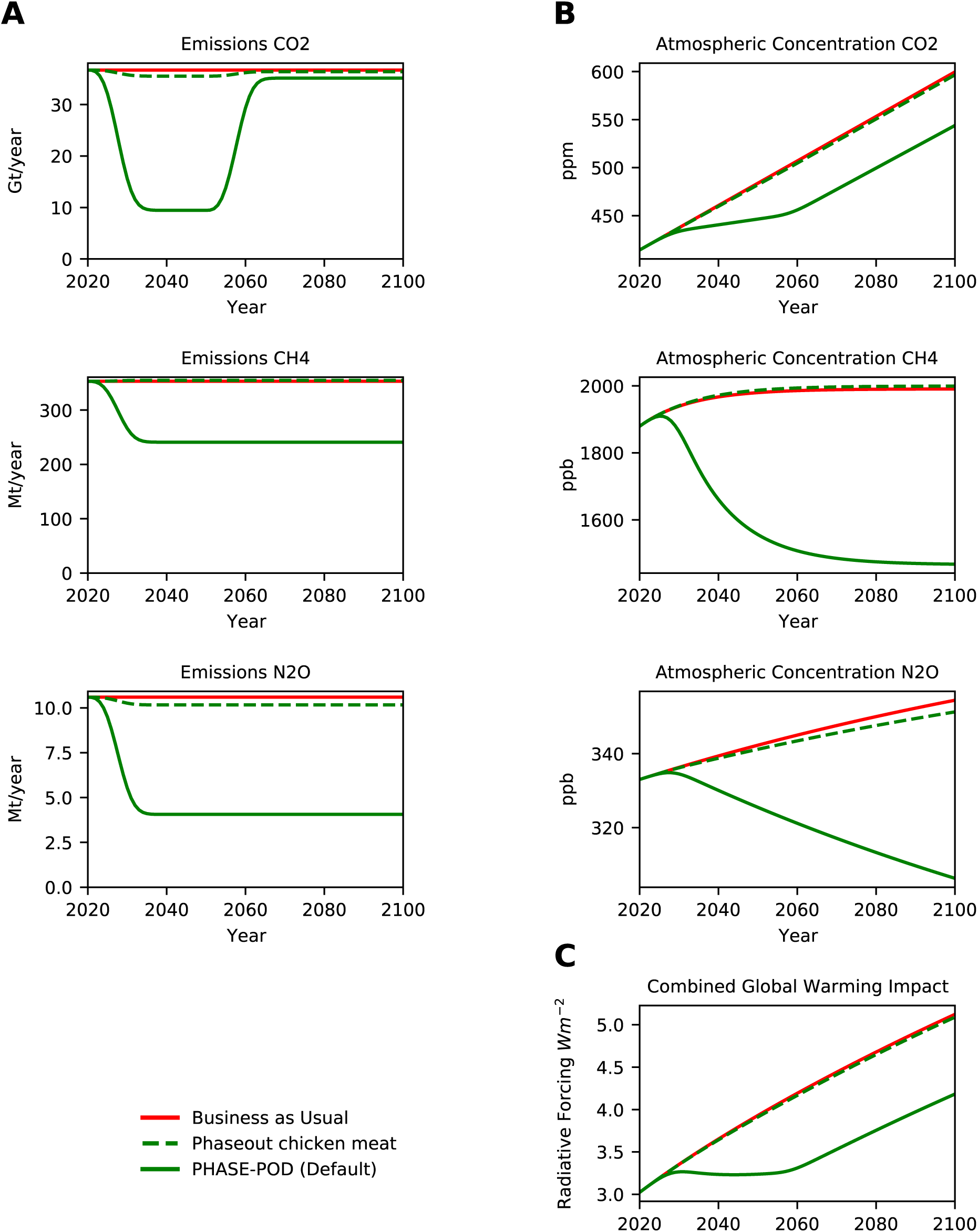
Phaseout of Chicken Meat. (A) Projected annual emissions of *CO*_2_, *CH*_4_ and *N*_2_*O* for each scenarios. (B) Projected atmospheric concentrations of *CO*_2_, *CH*_4_ and *N*_2_*O* under each emission scenario. (C) Radiative Forcing (RF) inferred from atmospheric concentrations in (B) by formula of (Myhre et al., 1998; Ramaswamy et al., 2001) as modified in MAGICC6 (Meinshausen et al., 2011). Only differences between PHASE-POD default assumptions (15yr phaseout, 30yr carbon recovery, 100% carbon recovery, BAU non-agriculture emissions, FAO crop replacement, and FAO animal ag emissions) are given.

**Figure 2-S23.**
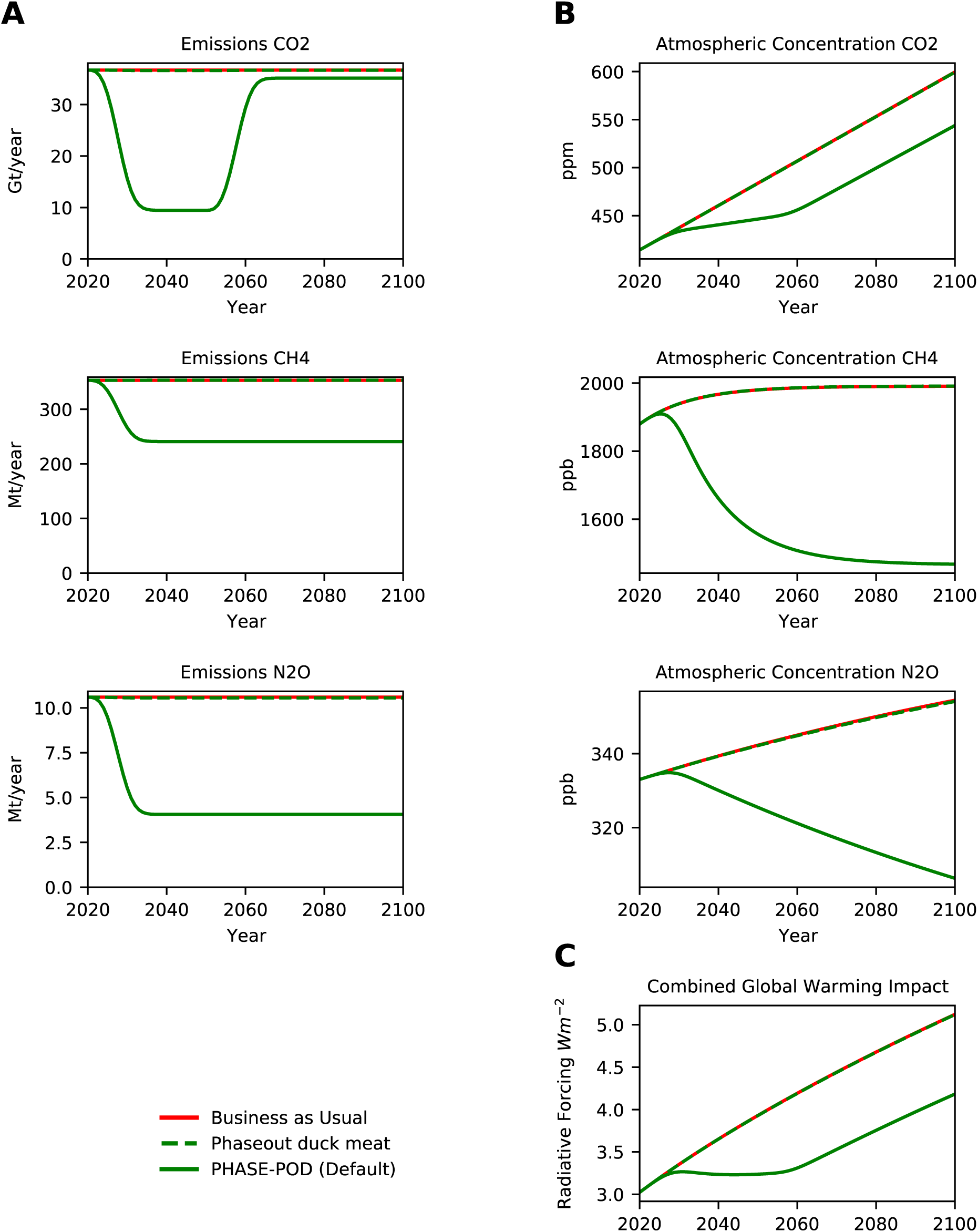
Phaseout of Duck Meat. (A) Projected annual emissions of *CO*_2_, *CH*_4_ and *N*_2_*O* for each scenarios. (B) Projected atmospheric concentrations of *CO*_2_, *CH*_4_ and *N*_2_*O* under each emission scenario. (C) Radiative Forcing (RF) inferred from atmospheric concentrations in (B) by formula of (Myhre et al., 1998; Ramaswamy et al., 2001) as modified in MAGICC6 (Meinshausen et al., 2011). Only differences between PHASE-POD default assumptions (15yr phaseout, 30yr carbon recovery, 100% carbon recovery, BAU non-agriculture emissions, FAO crop replacement, and FAO animal ag emissions) are given.

**Figure 2-S24.**
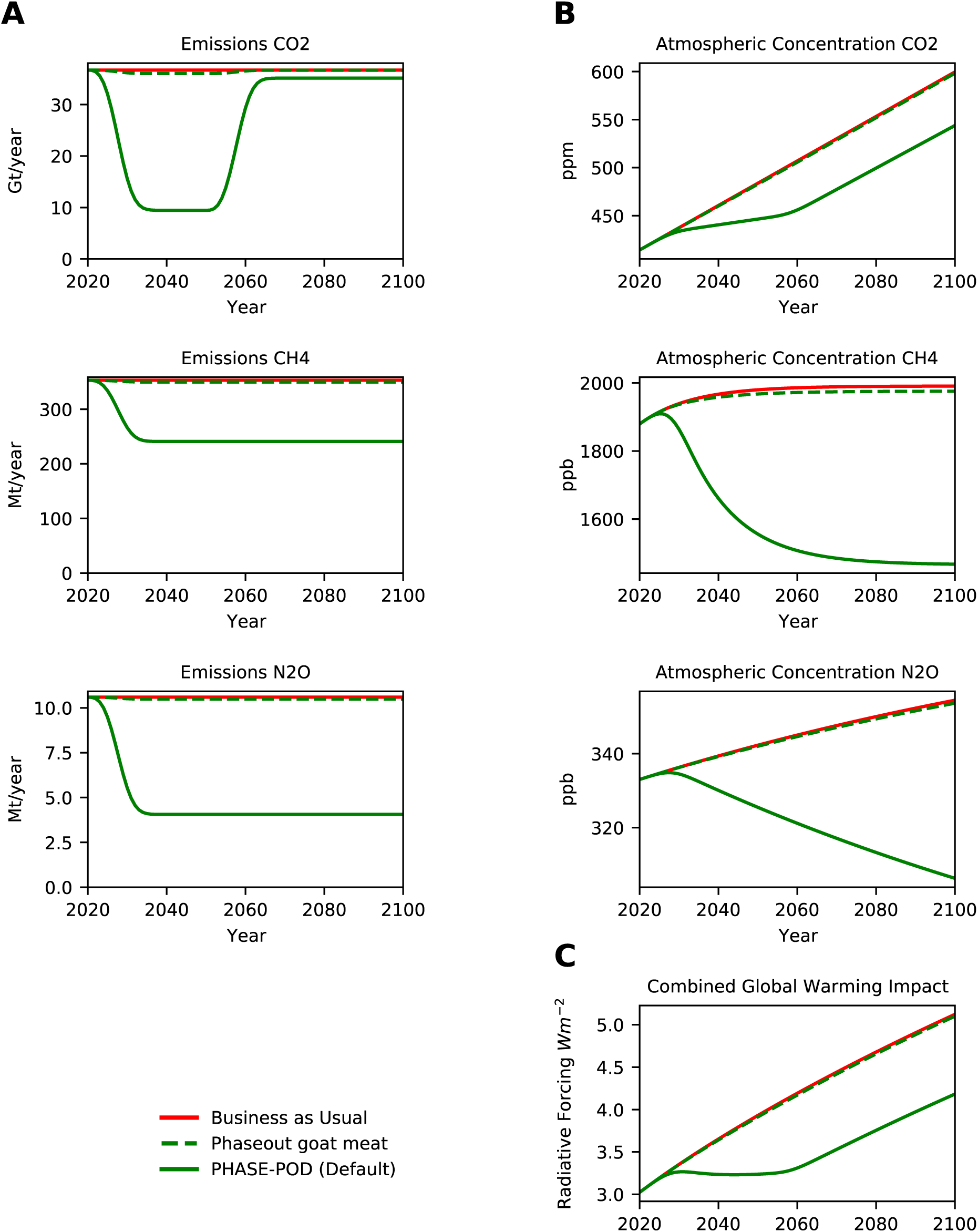
Phaseout of Goat Meat. (A) Projected annual emissions of *CO*_2_, *CH*_4_ and *N*_2_*O* for each scenarios. (B) Projected atmospheric concentrations of *CO*_2_, *CH*_4_ and *N*_2_*O* under each emission scenario. (C) Radiative Forcing (RF) inferred from atmospheric concentrations in (B) by formula of (Myhre et al., 1998; Ramaswamy et al., 2001) as modified in MAGICC6 (Meinshausen et al., 2011). Only differences between PHASE-POD default assumptions (15yr phaseout, 30yr carbon recovery, 100% carbon recovery, BAU non-agriculture emissions, FAO crop replacement, and FAO animal ag emissions) are given.

**Figure 2-S25.**
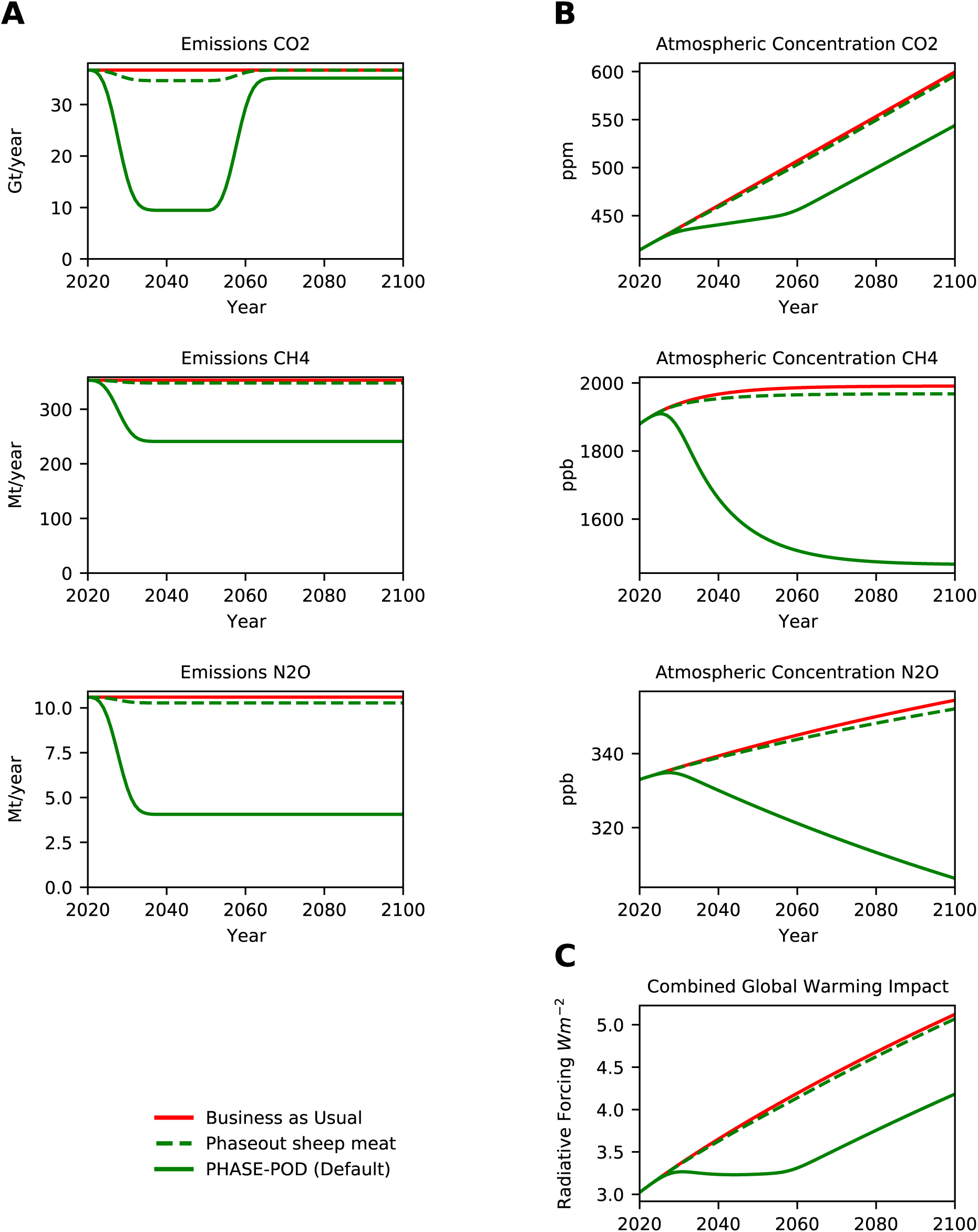
Phaseout of Sheep Meat. (A) Projected annual emissions of *CO*_2_, *CH*_4_ and *N*_2_*O* for each scenarios. (B) Projected atmospheric concentrations of *CO*_2_, *CH*_4_ and *N*_2_*O* under each emission scenario. (C) Radiative Forcing (RF) inferred from atmospheric concentrations in (B) by formula of (Myhre et al., 1998; Ramaswamy et al., 2001) as modified in MAGICC6 (Meinshausen et al., 2011). Only differences between PHASE-POD default assumptions (15yr phaseout, 30yr carbon recovery, 100% carbon recovery, BAU non-agriculture emissions, FAO crop replacement, and FAO animal ag emissions) are given.

**Figure 2-S26.**
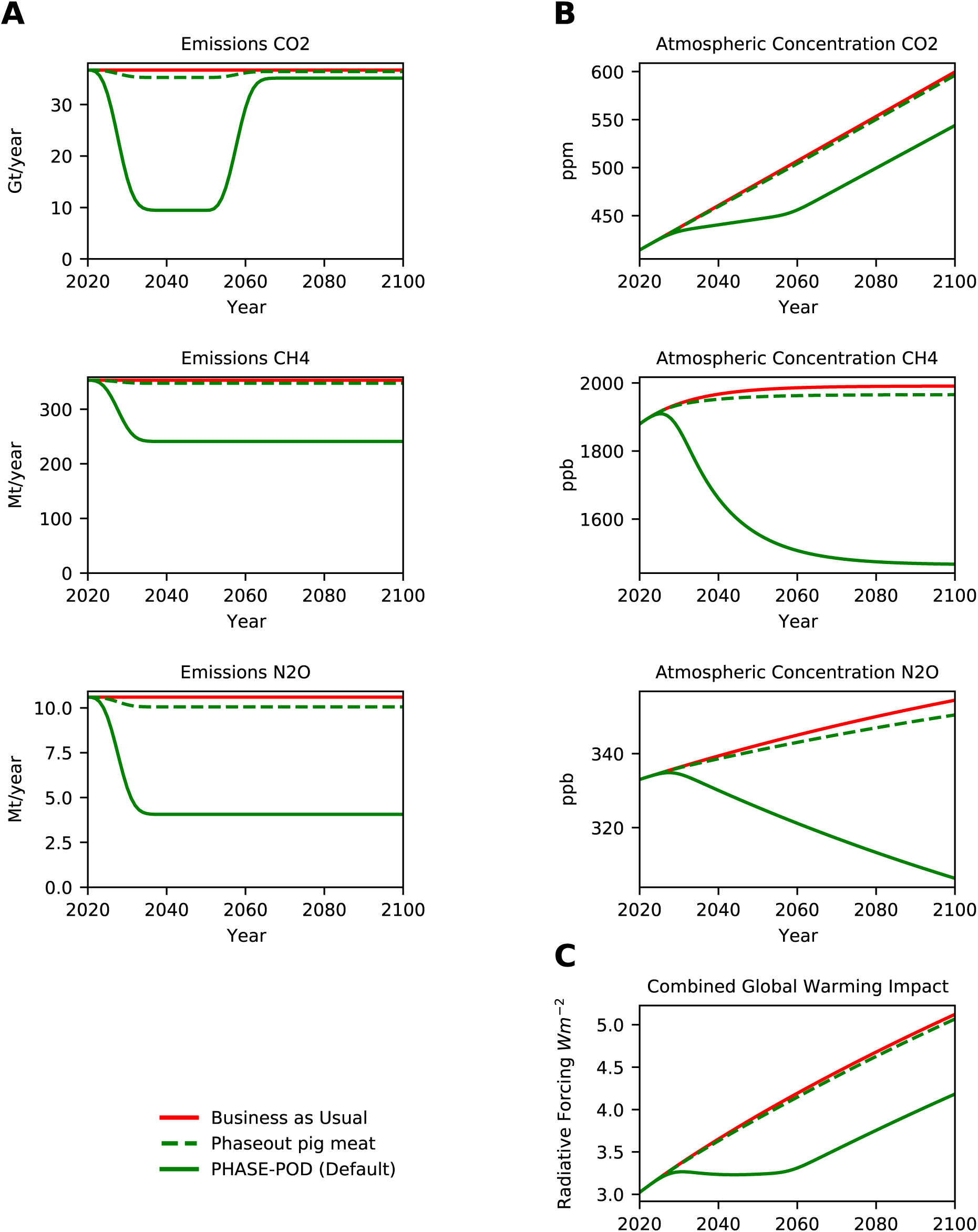
Phaseout of Pig Meat. (A) Projected annual emissions of *CO*_2_, *CH*_4_ and *N*_2_*O* for each scenarios. (B) Projected atmospheric concentrations of *CO*_2_, *CH*_4_ and *N*_2_*O* under each emission scenario. (C) Radiative Forcing (RF) inferred from atmospheric concentrations in (B) by formula of (Myhre et al., 1998; Ramaswamy et al., 2001) as modified in MAGICC6 (Meinshausen et al., 2011). Only differences between PHASE-POD default assumptions (15yr phaseout, 30yr carbon recovery, 100% carbon recovery, BAU non-agriculture emissions, FAO crop replacement, and FAO animal ag emissions) are given.

**Figure 2-S27.**
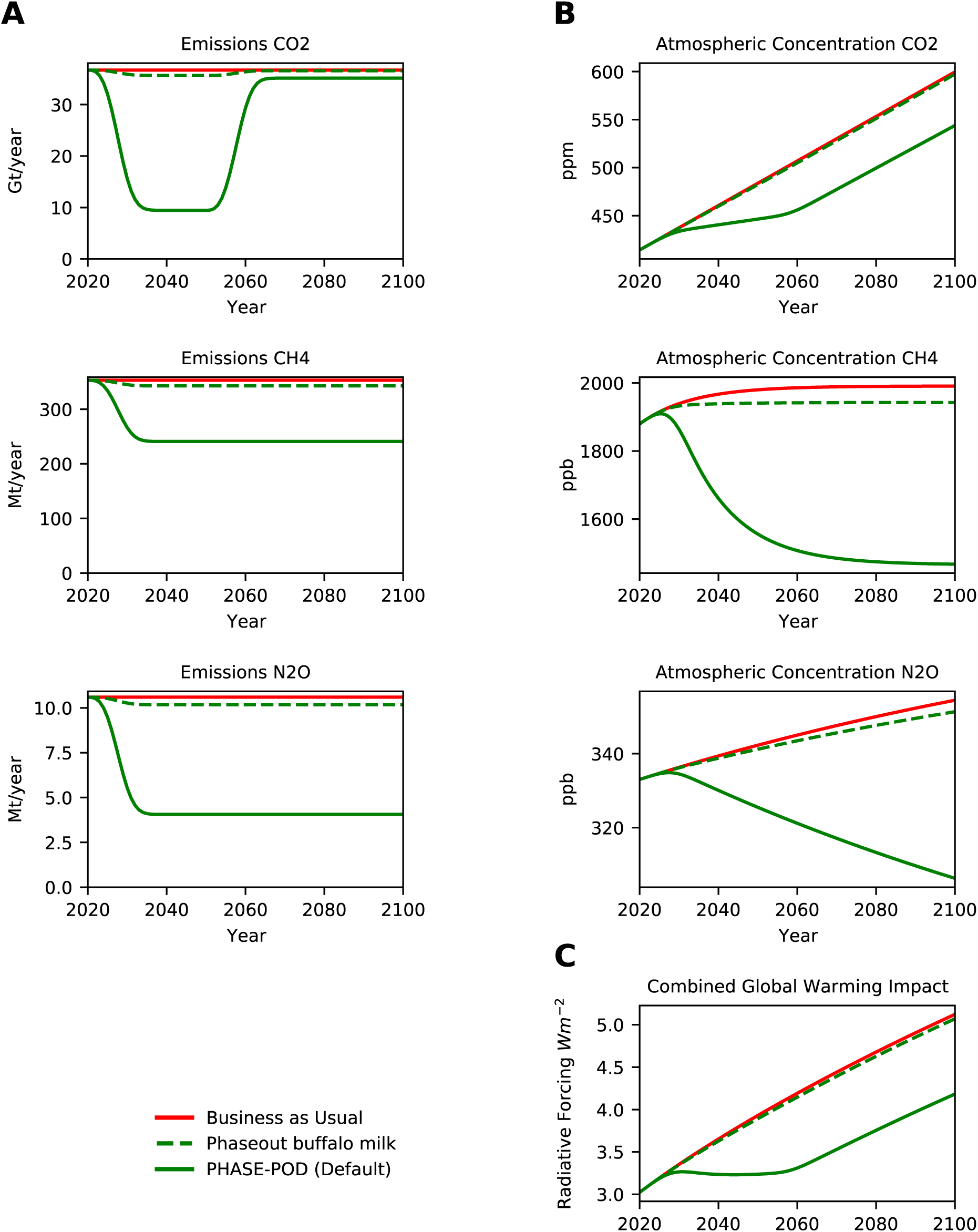
Phaseout of Buffalo Milk. (A) Projected annual emissions of *CO*_2_, *CH*_4_ and *N*_2_*O* for each scenarios. (B) Projected atmospheric concentrations of *CO*_2_, *CH*_4_ and *N*_2_*O* under each emission scenario. (C) Radiative Forcing (RF) inferred from atmospheric concentrations in (B) by formula of (Myhre et al., 1998; Ramaswamy et al., 2001) as modified in MAGICC6 (Meinshausen et al., 2011). Only differences between PHASE-POD default assumptions (15yr phaseout, 30yr carbon recovery, 100% carbon recovery, BAU non-agriculture emissions, FAO crop replacement, and FAO animal ag emissions) are given.

**Figure 2-S28.**
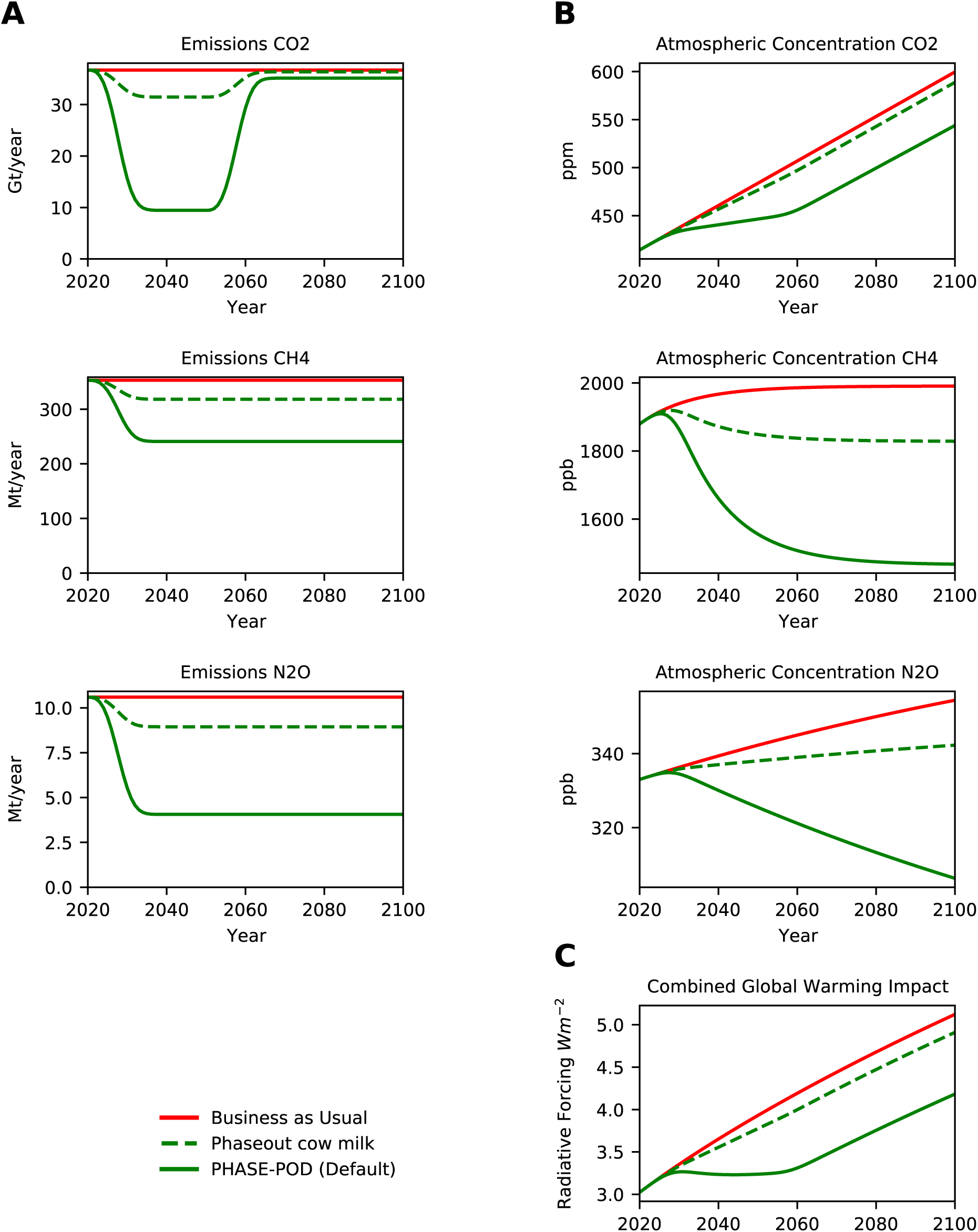
Phaseout of Cow Milk. (A) Projected annual emissions of *CO*_2_, *CH*_4_ and *N*_2_*O* for each scenarios. (B) Projected atmospheric concentrations of *CO*_2_, *CH*_4_ and *N*_2_*O* under each emission scenario. (C) Radiative Forcing (RF) inferred from atmospheric concentrations in (B) by formula of (Myhre et al., 1998; Ramaswamy et al., 2001) as modified in MAGICC6 (Meinshausen et al., 2011). Only differences between PHASE-POD default assumptions (15yr phaseout, 30yr carbon recovery, 100% carbon recovery, BAU non-agriculture emissions, FAO crop replacement, and FAO animal ag emissions) are given.

**Figure 2-S29.**
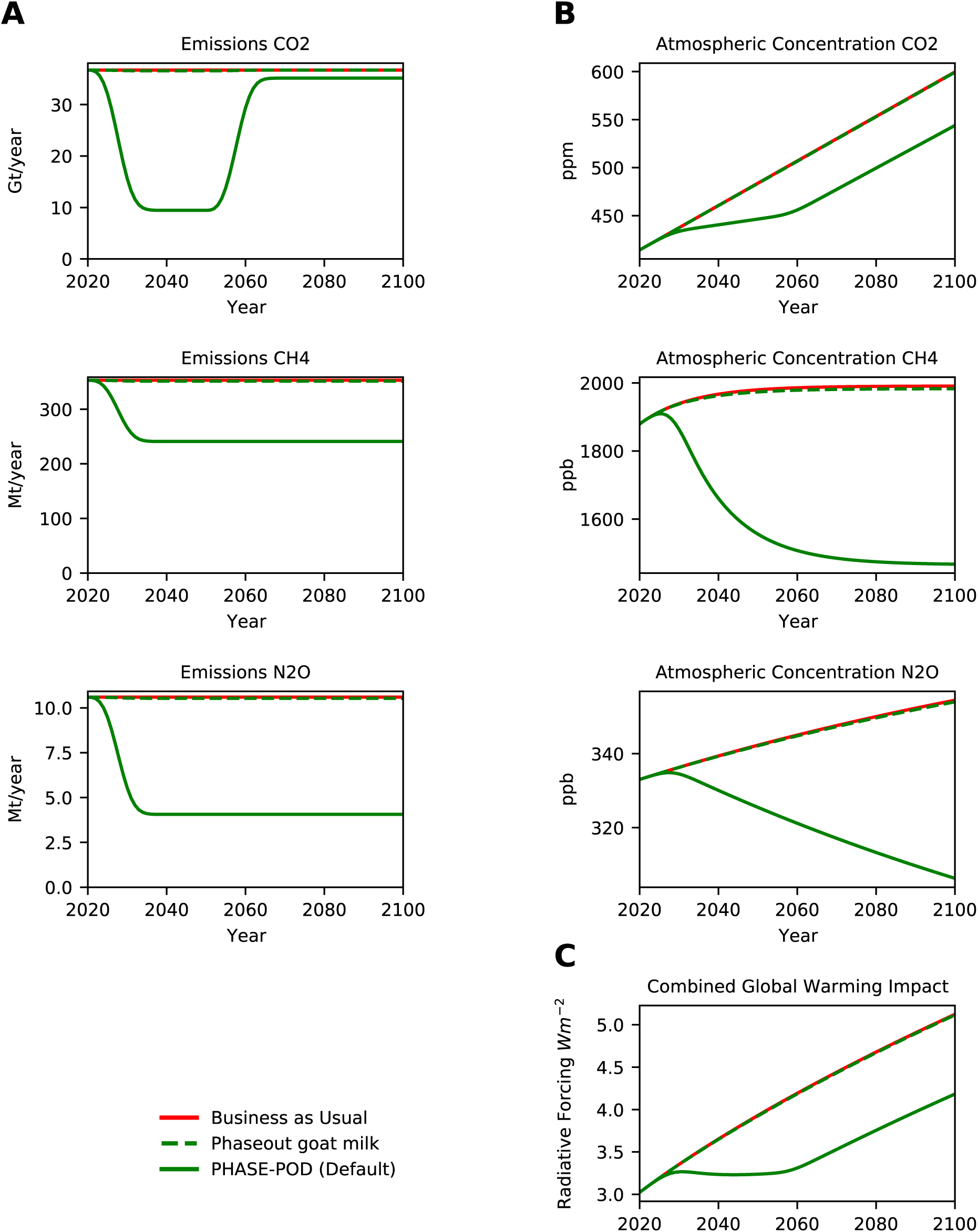
Phaseout of Goat Milk. (A) Projected annual emissions of *CO*_2_, *CH*_4_ and *N*_2_*O* for each scenarios. (B) Projected atmospheric concentrations of *CO*_2_, *CH*_4_ and *N*_2_*O* under each emission scenario. (C) Radiative Forcing (RF) inferred from atmospheric concentrations in (B) by formula of (Myhre et al., 1998; Ramaswamy et al., 2001) as modified in MAGICC6 (Meinshausen et al., 2011). Only differences between PHASE-POD default assumptions (15yr phaseout, 30yr carbon recovery, 100% carbon recovery, BAU non-agriculture emissions, FAO crop replacement, and FAO animal ag emissions) are given.

**Figure 2-S30.**
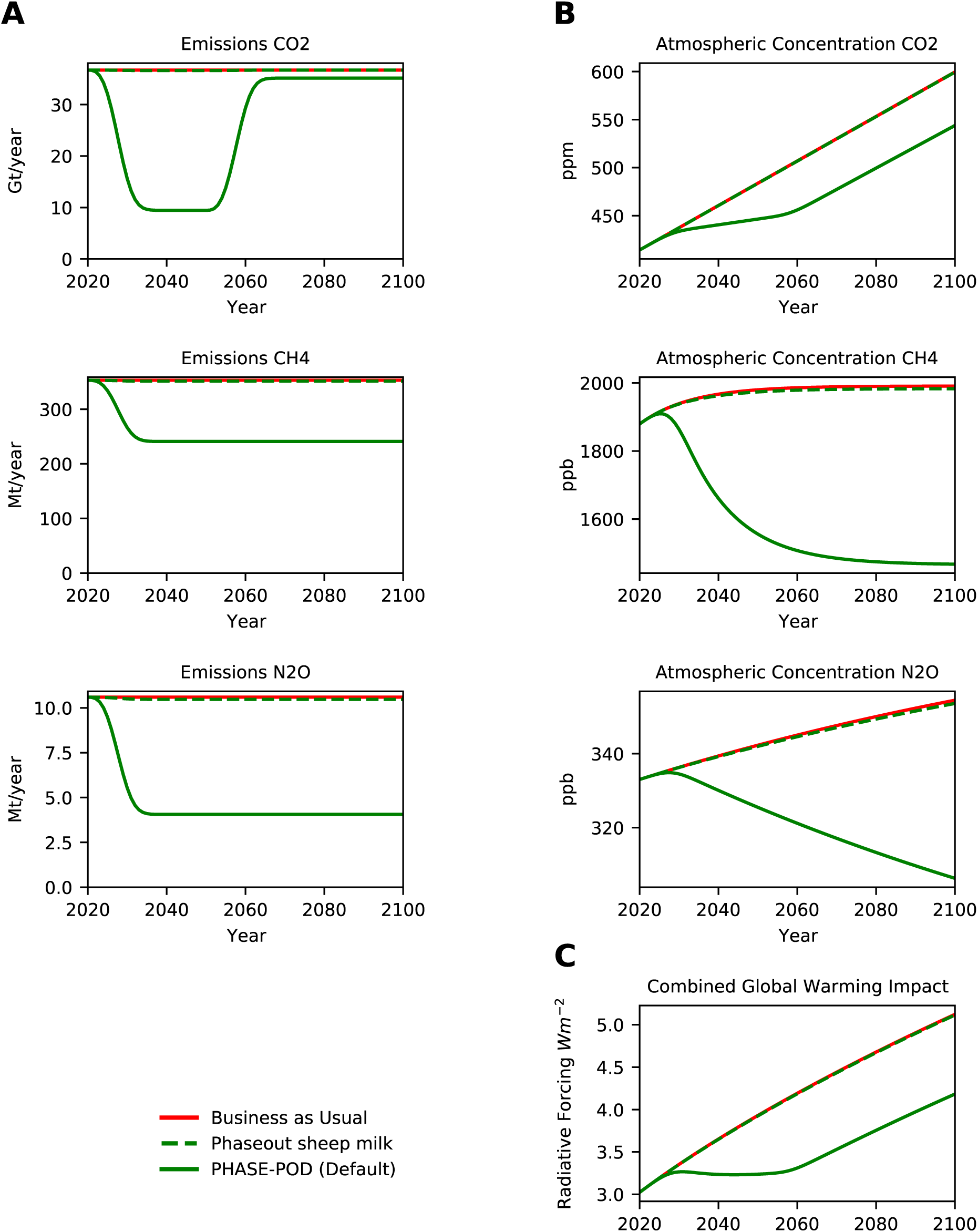
Phaseout of Sheep Milk. (A) Projected annual emissions of *CO*_2_, *CH*_4_ and *N*_2_*O* for each scenarios. (B) Projected atmospheric concentrations of *CO*_2_, *CH*_4_ and *N*_2_*O* under each emission scenario. (C) Radiative Forcing (RF) inferred from atmospheric concentrations in (B) by formula of (Myhre et al., 1998; Ramaswamy et al., 2001) as modified in MAGICC6 (Meinshausen et al., 2011). Only differences between PHASE-POD default assumptions (15yr phaseout, 30yr carbon recovery, 100% carbon recovery, BAU non-agriculture emissions, FAO crop replacement, and FAO animal ag emissions) are given.

**Figure 2-S31.**
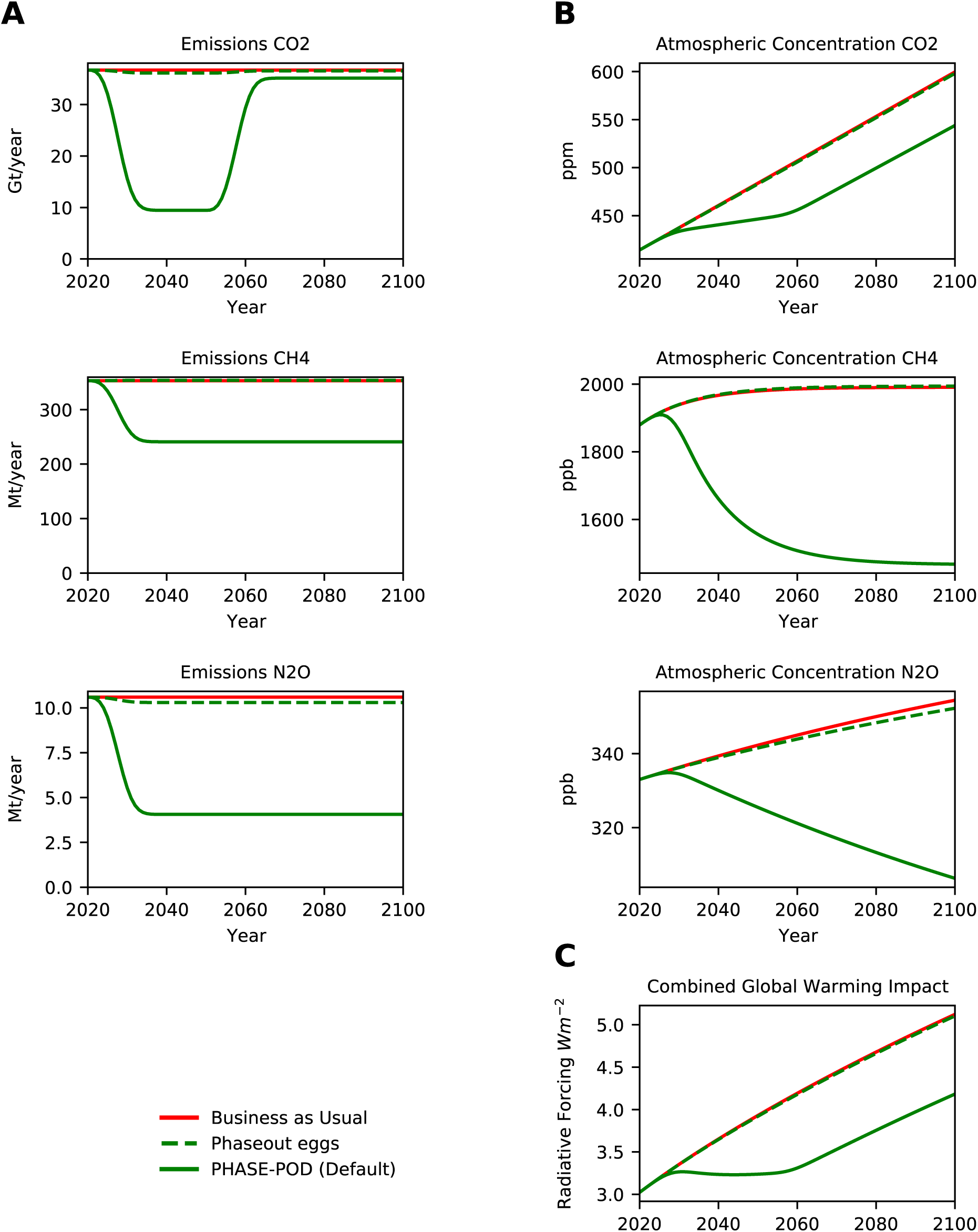
Phaseout of Eggs. (A) Projected annual emissions of *CO*_2_, *CH*_4_ and *N*_2_*O* for each scenarios. (B) Projected atmospheric concentrations of *CO*_2_, *CH*_4_ and *N*_2_*O* under each emission scenario. (C) Radiative Forcing (RF) inferred from atmospheric concentrations in (B) by formula of (Myhre et al., 1998; Ramaswamy et al., 2001) as modified in MAGICC6 (Meinshausen et al., 2011). Only differences between PHASE-POD default assumptions (15yr phaseout, 30yr carbon recovery, 100% carbon recovery, BAU non-agriculture emissions, FAO crop replacement, and FAO animal ag emissions) are given.

**Figure 3-S1.**
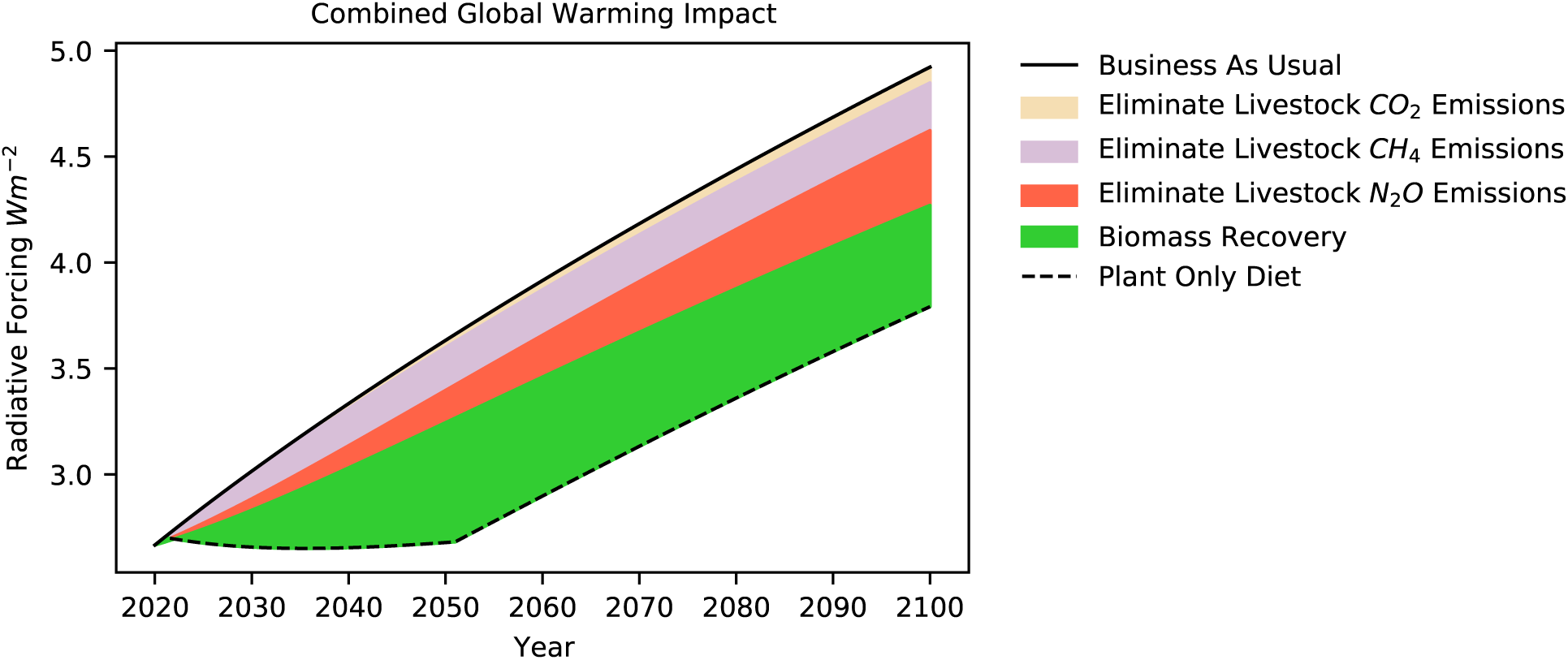
Immediate elimination of animal agriculture reduces global warming impact of atmosphere. Effect of eliminating emissions linked to animal agriculture and of biomass recovery on land currently used in animal agriculture on Radiative Forcing (RF), a measure of the instantaneous warming potential of the atmosphere. RF values computed from atmospheric concentrations in by formula of (Myhre et al., 1998; Ramaswamy et al., 2001) as modified in MAGICC6 (Meinshausen et al., 2011) with adjustment for gasses other than *CO*_2_, *CH*_4_ and *N*_2_*O* as described in text.

**Figure 3-S2.**
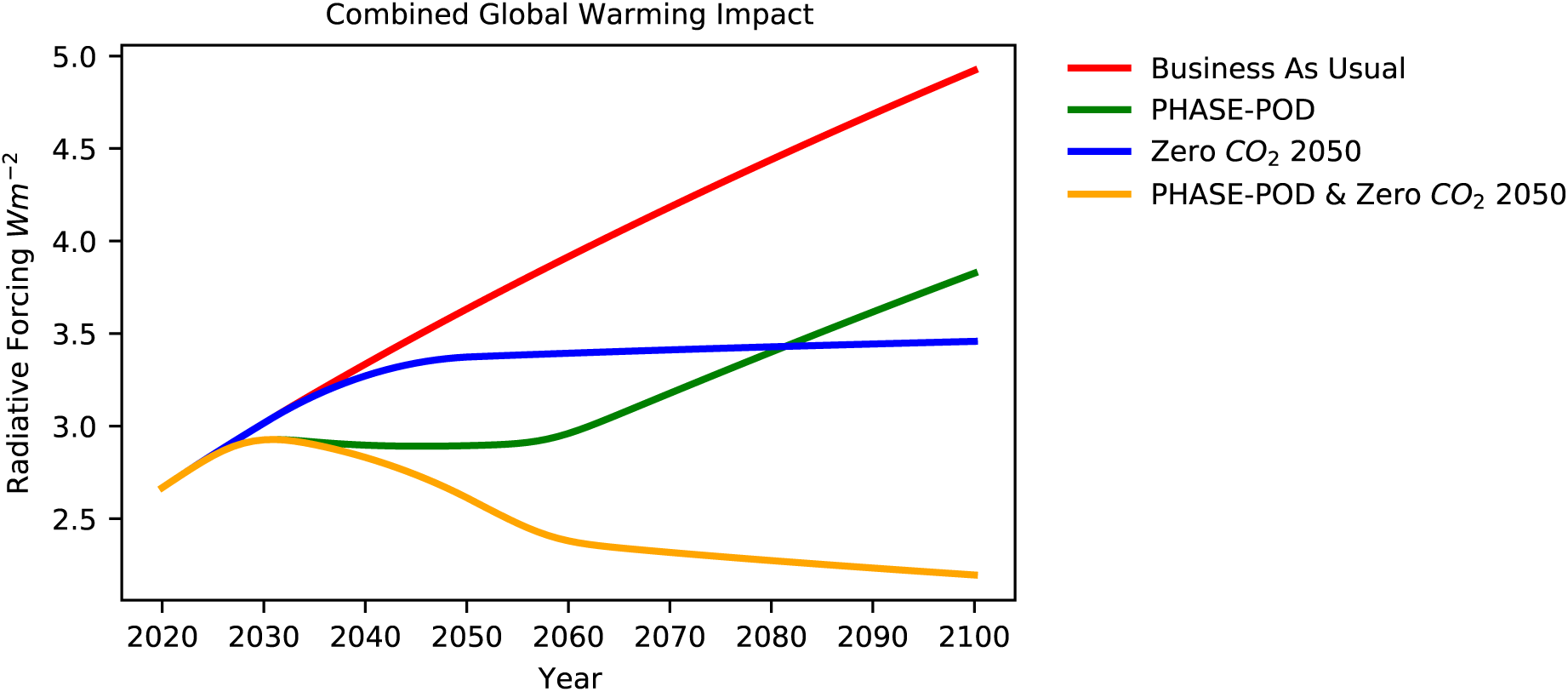
Similar effects of phaseout of animal ag and drawdown of *CO*_2_ emissions. Comparison of effects of PHASE-POD (a 15 year phaseout of animal agriculture) and a linear drawdown of all anthropogenic *CO*_2_ emissions between 2030 and 2050, and the two combined, on Radiative Forcing (RF), a measure of the instantaneous warming potential of the atmosphere. RF values computed from atmospheric concentrations in by formula of (Myhre et al., 1998; Ramaswamy et al., 2001) as modified in MAGICC6 (Meinshausen et al., 2011) with adjustment for gasses other than *CO*_2_, *CH*_4_ and *N*_2_*O* as described in text.

**Figure 4-S1.**
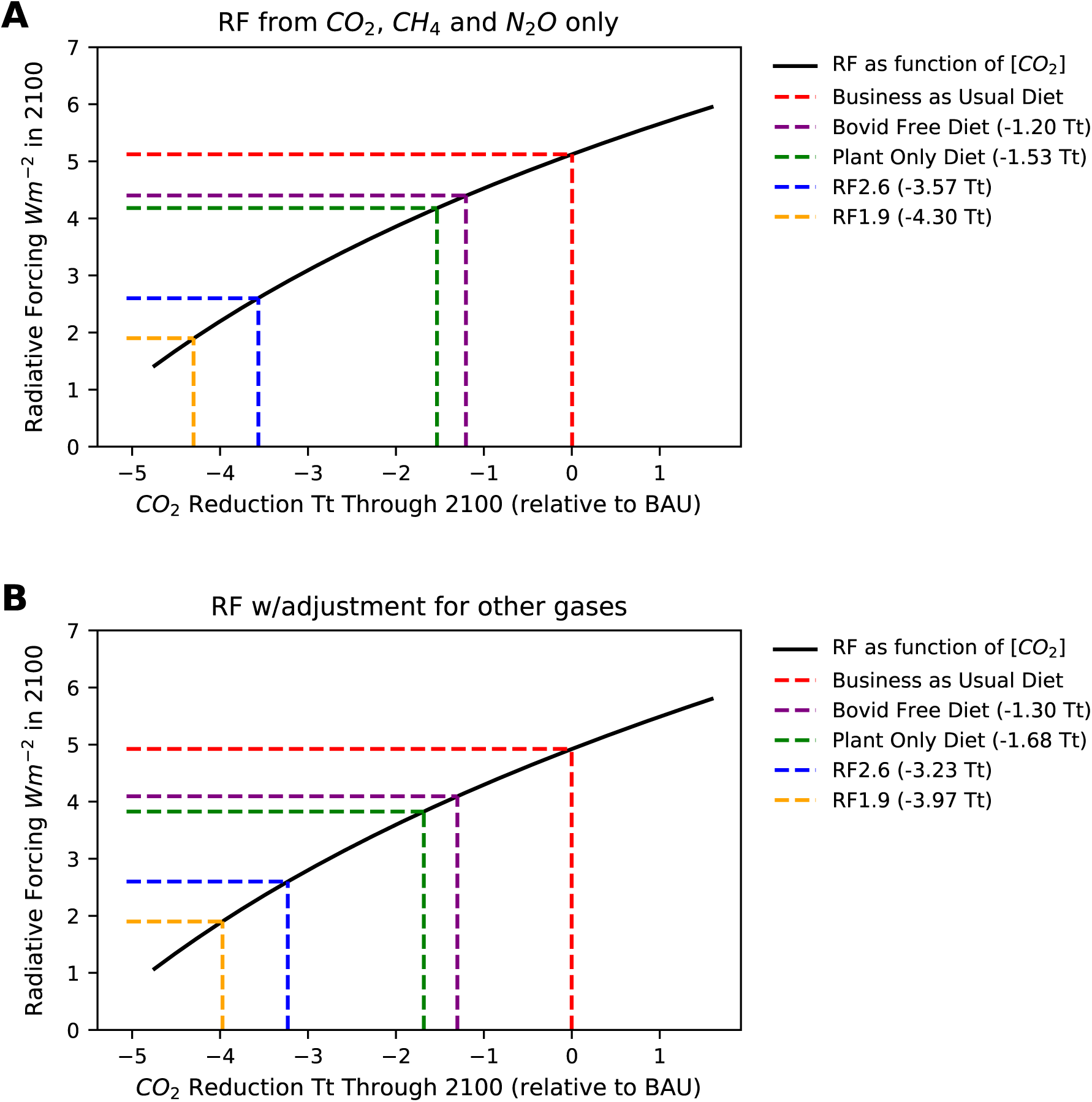
Full carbon opportunity cost of animal agriculture. We define the Emission and Land Carbon Opportunity Cost of animal agriculture as the total *CO*_2_ reduction necessary to lower the RF in 2100 from the level estimated for a business as usual (BAU) diet to the level estimated for a plant only diet (POD). For these calculations we fix the *CH*_4_ and *N*_2_*O* levels in the RF calculation at those estimated for the BAU diet in 2100 and adjust *CO*_2_ levels to reach the target RF. We also calculate ELCOC for just bovid sourced foods and determine the emission reductions necessary to reach RF s of 2.6 and 1.9, often cited as targets for limiting warming to 2.0 C and 1.5 C respectively. (A) Shows the results for RF directly calculated from *CO*_2_, *CH*_4_ and *N*_2_*O*, while (B) shows an RF adjusted for other gases using multivariate linear regression on MAGICC6 output downloaded from the SSP database.

**Figure 5-S1.**
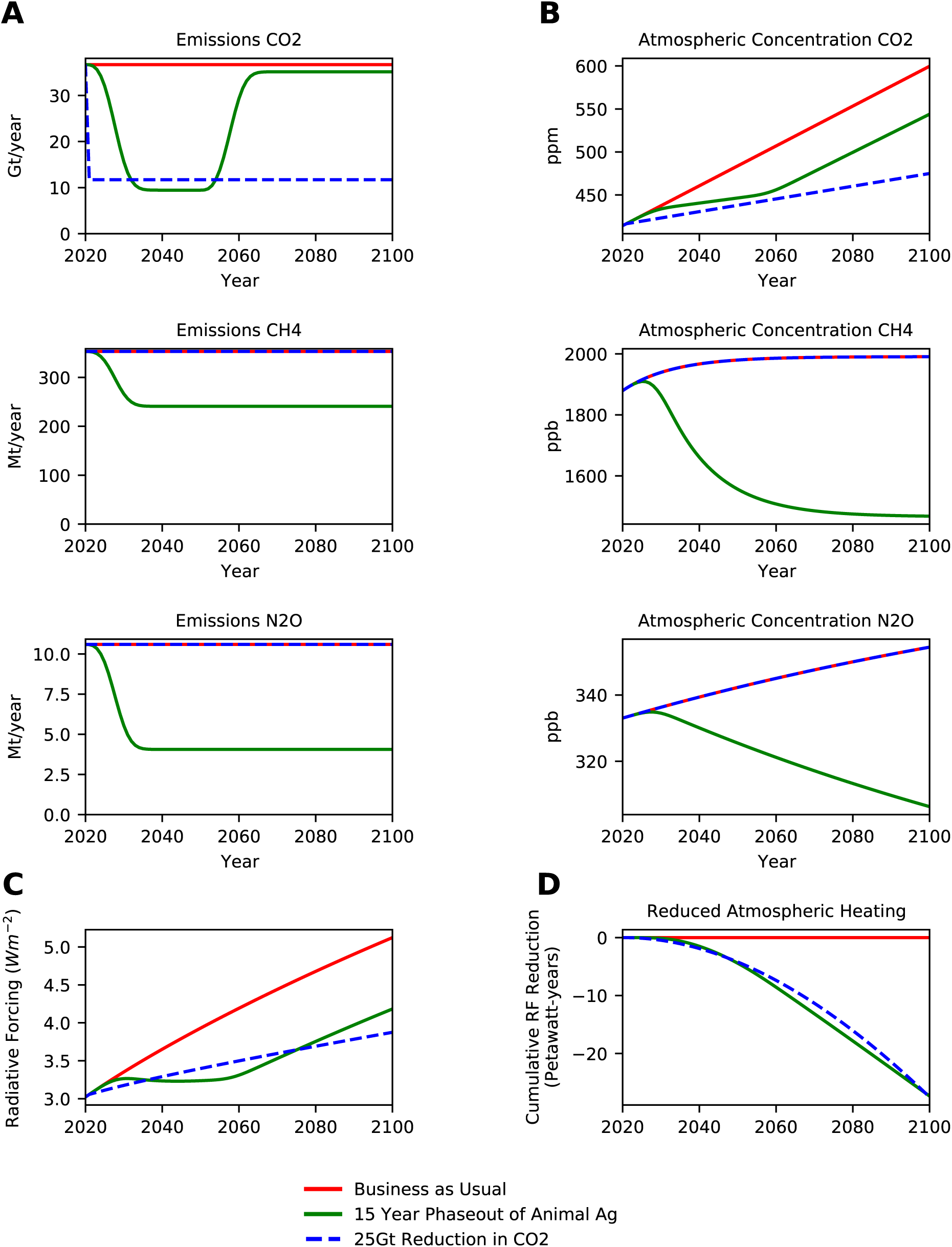
*ACO*_2_*eq* Calibration for PHASE-POD in 2100. (A) Projected annual emissions of *CO*_2_, *CH*_4_ and *N*_2_*O* for shown scenarios. (B) Projected atmospheric concentrations of *CO*_2_, *CH*_4_ and *N*_2_*O* under each emission scenario. (C) Radiation Forcing. (D) Cumulative difference between scenario and BAU of Radiative Forcing.

**Figure 5-S2.**
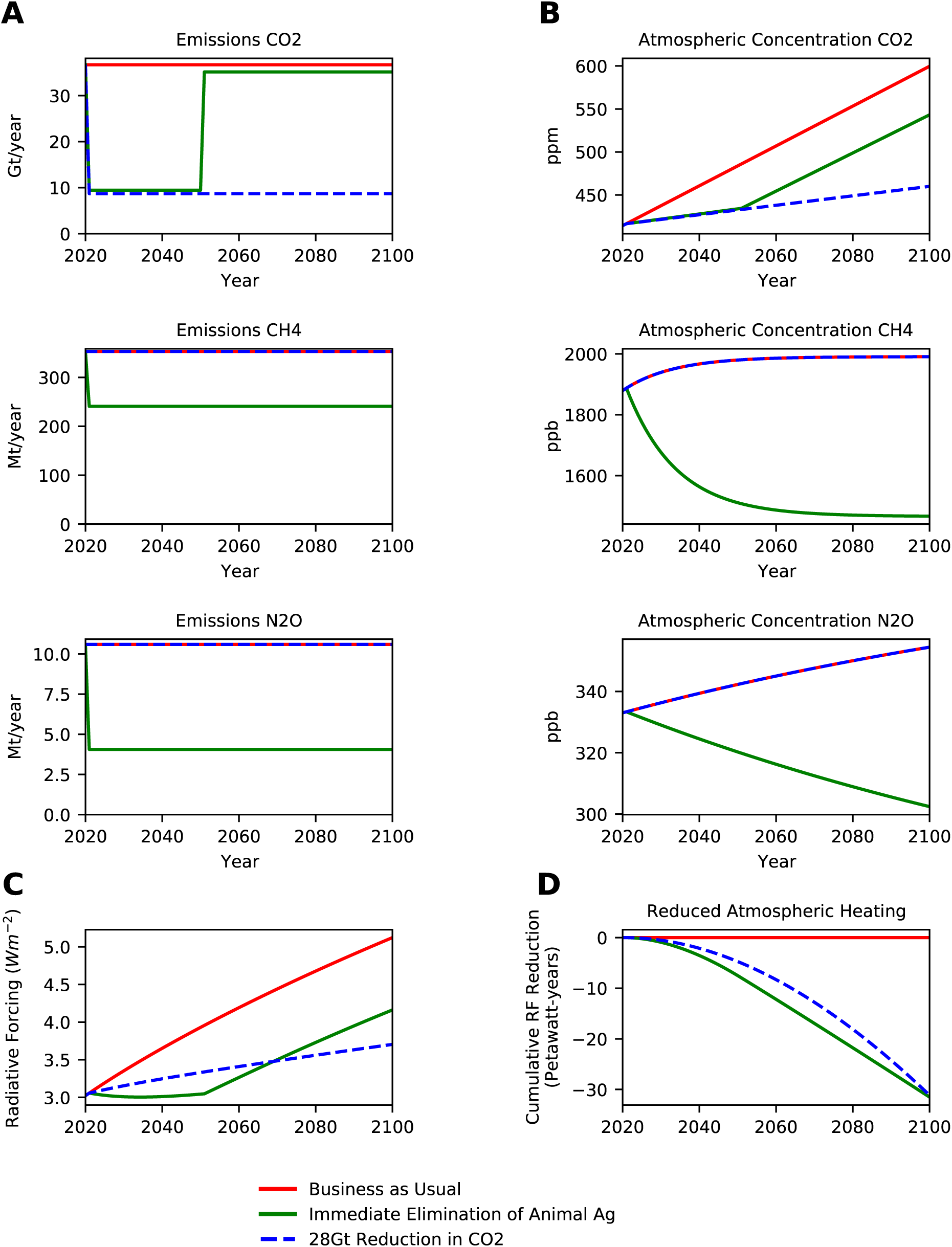
*ACO*_2_*eq* Calibration for IMM-POD in 2100. (A) Projected annual emissions of *CO*_2_, *CH*_4_ and *N*_2_*O* for shown scenarios. (B) Projected atmospheric concentrations of *CO*_2_, *CH*_4_ and *N*_2_*O* under each emission scenario. (C) Cumulative difference between scenario and BAU of Radiative Forcing.

**Figure 5-S3.**
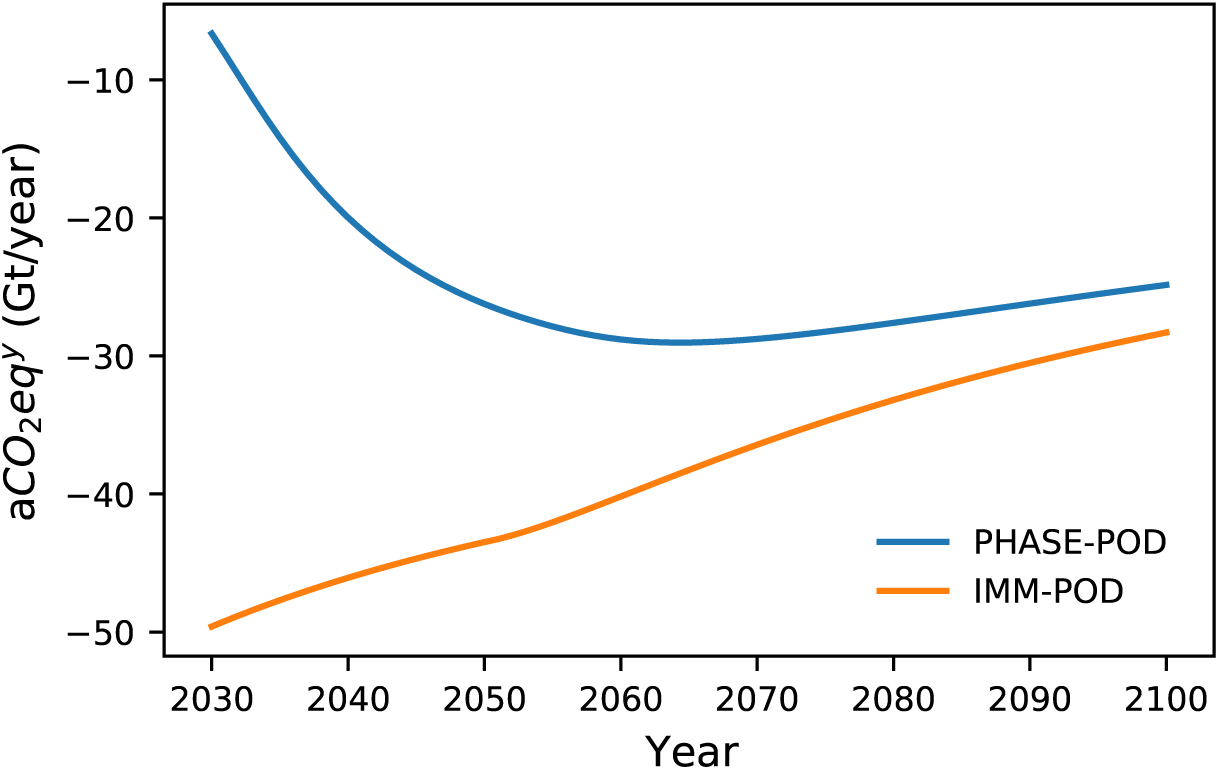
Emissions reduction equivalents of ending animal agriculture. The equivalent *CO*_2_ emission reductions associated with different interventions in animal agriculture, *aCO*_2_*eq*, vary with the time window over which cumulative warming impact is evaluated. These plots show, for immediate elimination of animal agriculture (IMM-POD) and a 15-year phaseout (PHASE-POD) how *aCO*_2_*eq^y^* which is the *aCO*_2_*eq* from 2021 to year y, varies with y. Because all of the changes in IMM-POD are implemented immediately, its effect is biggest as it is implemented and declines over longer time horizons (the decline in the first 30 years, when biomass recovery is occurring at a constant high right, is due to the slowing of annual decreases in atmospheric *CH*_4_ and *N*_2_*O* levels as they asymptotically approach new equilibria). In contrast, PHASE-POD builds slowly, reaching a maximum around 2060 when biomass recovery peaks.

**Figure 6-S1.**
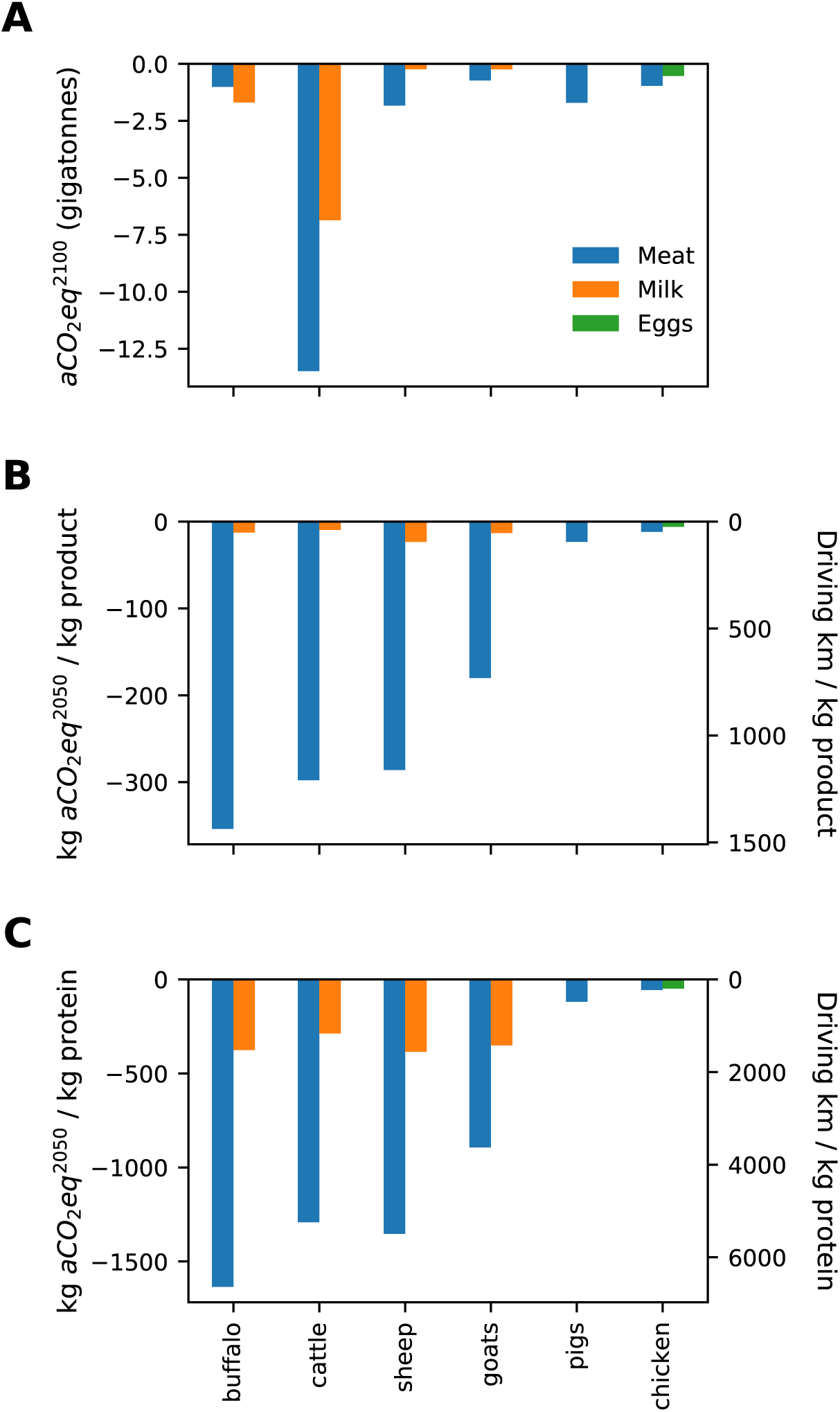
Emission equivalents of livestock products through 2100. We calculated the (A) total annualized CO_2_ equivalents through 2100, *aCO*_2_*eq*^2100^, for all tracked animal products, and the *aCO*_2_*eq*^2100^ per unit production (B) or per unit protein (C). For (B) and (C) we also convert the values to driving equivalents, assuming cars that get 10.6 km per liter of gas (the average of new cars in the United States).

**Figure 7-S1.**
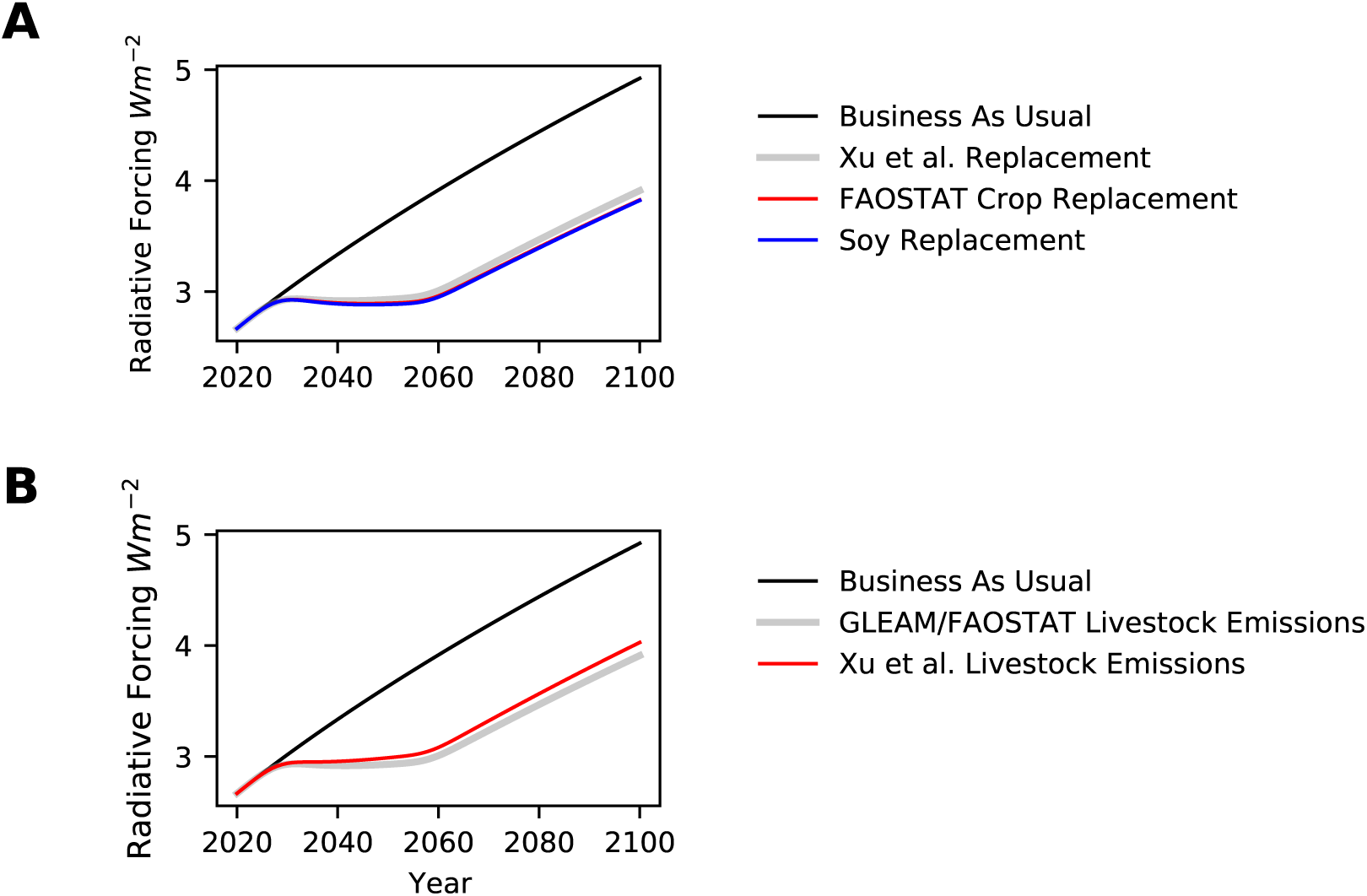
Sensitivity of impact of phaseout of animal agriculture to model assumptions. The grey line in each plot is PHASE-POD, the default scenario of 15 year phaseout, 30 year carbon recovery, livestock emissions from FAOSTAT, and a diverse plant replacement diet based on (Xu et al., 2021). (A) Effect of substituting the default plant based replacement diet from (Xu et al., 2021) with a diet based on all current human consumed crops using data from FAOSTAT, or a soy only replacment diet. (B) Effect of substituting default combined emissions of animal agriculture estimated via GLEAM and FAOSTAT with those from (Xu et al., 2021).

