## Supplementary material for "Rapid global phaseout of animal agriculture has the potential to stabilize greenhouse gas levels for 30 years and offset 68 percent of CO_2_ emissions this century": Peer Reviews and Responses

### Peer Review History

The correspondence has been redacted as follows to conform to bioRxiv rules:

The name of the journal has been replaced by JOURNAL.
The name of the editor has been replaced by EDITOR.
URLs that would reveal the name of the journal have been replaced by WEBSITE.

#### Version #1 Submitted to JOURNAL on July 19, 2021

Version 1 is materially similar to https://www.biorxiv.org/content/10.1101/2021.04.15.440019v3

#### Decision #1 Received from JOURNAL on September 9, 2021

Text in black is from the journal. Text in red is our response which as transmitted to the journal during resubmission.

Dear Dr. Eisen,

Thank you for submitting your manuscript to JOURNAL. After careful consideration, we feel that it has merit but does not fully meet JOURNAL's publication criteria as it currently stands. Therefore, we invite you to submit a revised version of the manuscript that addresses the points raised during the review process.

This paper has a clear potential to become a strong and impactful paper. The main issues that need to be addressed include a more rigorous discussion in relation to sensitivity tests for key assumptions that are fundamental to drive the results of the paper and the addition of corroboration and nuance in relation to the way that the results are presented. The latter includes discussing some of the trade-offs, barriers, and real plausibility in relation to eliminating animal agriculture.

Please submit your revised manuscript by Oct 17 2021 11:59PM. If you will need more time than this to complete your revisions, please reply to this message or contact the journal office at JOURNAL. When you're ready to submit your revision, log on to WEBSITE and select the 'Submissions Needing Revision' folder to locate your manuscript file.

Please include the following items when submitting your revised manuscript:

- A rebuttal letter that responds to each point raised by the editor and reviewer(s). You should upload this letter as a separate file labeled 'Response to Reviewers'.
- A marked-up copy of your manuscript that highlights changes made to the original version. You should upload this as a separate file labeled 'Revised Manuscript with Track Changes'.
- An unmarked version of your revised paper without tracked changes. You should upload this as a separate file labeled 'Manuscript'.

Guidelines for resubmitting your figure files are available below the reviewer comments at the end of this letter.

We look forward to receiving your revised manuscript.

Kind regards,

EDITOR

#### Review Comments to the Author

Please use the space provided to explain your answers to the questions above. You may also include additional comments for the author, including concerns about dual publication, research ethics, or publication ethics. (Please upload your review as an attachment if it exceeds 20,000 characters)

Reviewer #1: The manuscript is technically sound, and the data supports the conclusions. However, as I have detailed in my reviewer report, I think the authors need to more rigorously discuss/sensitivity test several of the assumptions in their methodology, and this is why I believe the manuscript currently only partly meets publication criteria. For this reason, and several other more minor comments detailed in my report where I think data used needs to be better justified or changed (e.g their analogy to driving, aquaculture), the statistical analysis is also partly satisfied at present. I have tried to be as helpful as possible in my comments regarding what the authors need to do to meet these criteria, and I am confident that once this is complete, they will have a strong and impactful paper.

We have made multiple changes in response to the reviewers suggestions, as detailed below. Specifically, we have added a section exploring the sensitivity of our results to assumptions made in our analyses. The assumptions explored include how long it takes to phase out animal agriculture, the carbon displaced by animal agriculture, the duration and extent of carbon recovery, the scale of animal-linked emissions, and the choice of replacement diet. Key results are now shown in Figure 7.

Reviewer #2: The topic of the paper, how to tackle the climate emergency and how reduced or eliminated animal agriculture can contribute, is highly relevant.

The singular focus of the paper on animal agriculture as THE climate change solution should be nuanced, as many sectors and processes contribute to GHG emissions. It is not possible to tackle climate change by focusing on one single solution (or even one single sector). In addition, some of the trade-offs and barriers/challenges in relation to eliminating animal agriculture need to be brought more clearly to the readers' attention.

Although the focus of this paper is on animal agriculture, it was not our intention to suggest that it is the only solution. Indeed in both the text and analyses we emphasized that even if the full extent of the potential we highlight here were realized, it would still not be enough to solve the climate crisis. We have made this point clearer at multiple points in the manuscript. We also explore the challenges to eliminating animal agriculture in more detail.

Most importantly, however, the results as presented (e.g. offset of 70% possible by eliminating animal agriculture) that could justify this strong focus, need to be corroborated and presented with much more nuance. The methodology followed in the manuscript combines many different calculations based on many assumptions and with large separate uncertainties associated with it.

As discussed above, we have added a section exploring the sensitivity of our results to assumptions made in our analyses so that readers can get a better sense of the degree of uncertainty in our calculations and its sources.

More detailed comments (about methodology as well as the other parts of the manuscript) are attached.

We have responded below to the reviewer's comments and modified the manuscript where appropriate.

#### Reviewer 1

##### Reviewer Report

The role of animal agriculture in driving climate change is an important and often complex topic, where science can help guide societal understanding and policy makers. This study contributes to this scientific basis, and is novel in combining the impact of livestock emissions reductions with potential biomass carbon recovery, to provide a 'climate opportunity cost' of animal agriculture. The manuscript is generally very well-written and clear regarding how the study was carried out, the assumptions and data used, and the conclusions are hard-hitting.

My comments are long: I have tried to be as thorough as possible to help the authors strengthen this manuscript before publication, and mostly the comments should be straight-forward to address, relating to clarifications, consistencies, and suggestions for improved referencing.

I do have two more significant comments relating to the assumptions used in the modelling which need addressing (see below, major comments 1) and 2)). The authors are generally clear and up-front about their assumptions, and do emphasize that this is a simple study, however, I think further clarity is needed, and a discussion in particular of two of the assumptions and how they have shaped the results (soy as a replacement of protein and potential biomass carbon recovery). I think at the least it needs to be acknowledged that these two assumptions likely overstate the magnitude of the results, and why. The authors could also provide results of a simple sensitivity analysis (perhaps in the SI) of these assumptions: this would strengthen the main conclusions of their study, that a phased transition from animal agriculture to plant-based diets would lead to significant emission reduction and biomass carbon recovery, delivering a major contribution towards the Paris targets.

##### Major Comments

1) The assumption that soy protein is used to replace all animal protein. If the authors don't provide results of sensitivity analysis on this assumption, they need to at least acknowledge that the impact would be different under other plant proteins. This assumption could be discussed in the Discussion, when the biomass recovery assumption is discussed. In reality, a range of plant

proteins are being used now and would be used under further shifts to plant-based diets. There are of course many regions where other plant protein crops would be favored, biophysically, culturally and socio-economically, over soy protein. Additionally, a proportion of plant-based protein consumed is processed, and would be under a future plant-based scenario, and this will also influence the emissions of a plant-based diet, e.g Santo et al. (2020, Frontiers in Sustainable Food Systems) compares environmental impact of processed plant-based foods to wholefood plant foods. In reality, the assumption that soy protein would replace all animal protein likely overstates the emission reduction of this phased out animal agriculture scenario somewhat.

We agree that this assumption is on the low end of potential emissions from replacement diets. We felt this was in the spirit of the paper's effort to quantify the potential gains due to elimination of animal agriculture, but agree that this is probably too aggressive an assumption. So in the revised manuscript we instead include emissions from non-animal foods scaled to replace the protein from animal agriculture. In the sensitivity section we compare this to emissions from a diverse plant-based diet based on data from a recent paper from Xu et al. that used LCA to estimate global emissions from plant and animal foods (see Figure 7A-S1).

2) Assumption on biomass recovery. This assumption and uncertainty around it are already discussed (page 13) but this needs expansion, particularly because this is the major driver of the study's results. The authors should acknowledge other rising pressures on land besides animal agriculture: energy (solar, wind, biomass), urbanization, and climate change (wildfires, drought), all of which are expected to increase this century. My understanding of the methods of Hayek et al. (2021) – and the authors can correct me if they think this is wrong – is that these other pressures on land were not accounted for (though a rising population was). As the purpose here is a self-described simple analysis with clear assumptions, I don't think the authors necessarily need to model these changes from other land-use pressures - which of course would be very challenging and full of uncertainty - but they should acknowledge that the likely biomass recovery would in reality fall below the potential level they use because of these other pressures, and potentially by a quite a sizeable amount. They could consider a simple sensitivity analysis to show they did consider this in their study, and that the main conclusions remain unchanged, even if, say, a third of that biomass carbon recovery wasn't realized.

The reviewer is correct that neither Hayek nor Strassburg model these other pressures on land use, although the data we use from Hayek did account for the land needed to grow plant-based foods for a rising population. It isn't feasible to accurately model how those additional land pressures might manifest, but to account for the wide range of possible outcomes, we have modeled a range of carbon recovery fractions and included these numbers in the new section on Sensitivity Analysis, and show them in Figure 7D. We have also modeled the low and high estimates of above ground biomass recovery from Hayek, shown in Figure 7C. While neither of these answers the question of how much carbon recovery we can reasonably expect, it does provide a framework for understanding the value of different levels of carbon recovery.

We also note that the likely land impact of urban expansion, solar and wind are relatively small compared to the land footprint of animal agriculture.  Moreover since, in general their impact is relatively independent of the business as usual vs  animal agriculture phase out scenarios, those impacts would not affect our conclusions. A major potential competitor for land currently used for animal agriculture is BECCS. As this would also remove carbon, presumably at a higher rate than regrowth of native biomass or why do it, this would further increase the climate value of eliminating animal agriculture.

#### Other comments

**Abstract:**

1) This is a study on global phaseout of animal agriculture but 'global' isn't used in the abstract – I think it would add clarity to put it in e.g "a 15 year phaseout of animal agriculture globally".

We have changed the title to reflect the global nature of our study and emphasized this in the manuscript.

2) 'Via' on first line, and elsewhere in manuscript – personally, I find this too informal/colloquial – 'through' instead?

Through is better and we have changed it.

3) Comma after 'However' on 3rd line.

All howevers are now appropriately punctuated.

4) Freeze in radiative forcing for 30 years ***from*** 2030 (for clarity).

Done.

5) Offset 70% of ***current*** anthropogenic emissions (for clarity).

This sentence was modified.

**Significance statement:**

1) Use of 'global-warming' here but 'global warming' elsewhere – consistency needed.

Done.

2) "70% reduction in the use of fossil fuels in energy and transportation" – I didn't come across the methods or references for this calculation in the manuscript?

We have updated this statement for clarity and now write "would be equivalent to a 68% reduction in carbon dioxide emissions", which is explained in the text on p. 12 as follows:

Figure 5 shows the aCO2eq for different scenarios for reference years 2050 (to capture short term impacts) and 2100 (Figure 5-S3 shows the full dependence of aCO2eq on the reference year). The aCO2eq2100 for PHASE-POD is -24.8 Gt/year. As global anthropogenic CO2 emissions are currently approximately 36 Gt/year, that PHASE-POD would have the same effect, through the end of the century, as a 68% reduction of CO2 emissions.

**Main Text:**

1) 2nd paragraph (references Hristov et al. 2013a ….): other good reference to add: the recent meta-analysis on LCAs of different beef management practices, Cusack et al. (2021, Global

Change Biology); Springmann et al. (2018, Nature), who compare multiple means of reducing environmental pressures of the food system; Poore and Nemecek (2018).

Done.

'Sizeable' is a more appropriate word than 'some' at the start of this sentence: other means of achieving food-system emission reductions are not as large as plant-based diets, but they are sizeable and important. I also think the sentence should include "plant-based diets" as an option because it makes more sense when in the next sentence you write "of these options", and you can then cite the important Poore and Nemecek meta-analysis on plant and animal food emissions in that first sentence.

Agreed. Rewritten for clarity.

2) 3rd paragraph: MacLeod et al. (2020) is a study on aquaculture emissions, but this is a point about the GHG reduction of plant-based diets – reference used in error in this sentence? Please also check use of MacLeod et al. (2018) here. I also don't think the Long Shadow report (Steinfeld et al., 2006) is best used here: it covers the environmental impact of animal agriculture well, but doesn't quantify the GHG benefits of shifts towards plant-based diets (e.g as done in Poore and Nemecek, 2018, Nature).

This phrase was removed in a rewrite for clarity and flow.

3) Livestock emissions are stated as 15% of anthropogenic emissions: the authors cite the Long Shadow report here (Steinfeld et al. 2006) but this report actually cited an 18% figure. However, the methodology used to reach that 18% figure was criticized. Until recently the more commonly accepted figure has been the FAO's 14.5%, however, Twine (2021, Sustainability) provides a more up-to-date estimate of a minimum of 16.5%. The authors should use the Twine estimate, perhaps also acknowledging the FAO's 14.5% prior estimate.

Done.

The authors calculate that livestock emissions equate to 4 % CO2, 35 % CH4, 66 % N20: I'm curious to know what their own calculations here translate to in terms of a % total CO2eq?

We have now included this in the text. It is 6.3 Gt, with the difference relative to other (generally higher) estimates coming from our not including $CO_2$ emissions due to ongoing land use change.

4) End of fifth paragraph: use of 'rapidly': this is certainly true of the short-lived atmospheric emissions. I did consider if it was true of biomass recovery: Strassburg et al. (2020) do state that carbon stocks of tropical forests (which make up most of the area of restored ecosystems in their modelling) return to >50% of reference ecosystem carbon stocks in the first 20 years – so I think the authors are justified in the use of 'rapidly' here.

It is true that 'rapidly' is very context dependent, and believe, as you have concluded as well, that given the time frames over which people think about global warming, it is appropriate here.

5) Page 10: linking of 1.9 and 2.6 Wm-2 to 1.5 DC and 2 DC: this is an important linkage and I think it would be beneficial to the reader if the specific chapter of the IPCC report was cited here.

Done.

6) Sentence on page 13 “As slower biomass recovery…..” – this sentence sounds incomplete/doesn’t make sense. Is there supposed to be a comma between the previous sentence and this one perhaps?

Fixed.

7) It is good to see on page 14 consideration to the uneven social and economic impact of this transition.

Thank you. It is one of the major challenges we all face in making dietary change happen with minimum social damage, and it is something we are beginning to work on.

8) Use of CRFD: this is an important way to look at warming/cooling potential. It isn’t my area of expertise, but I believe this approach is similar/synonymous with the recently introduced GWP* metric (e.g Allen et al., 2018, Climate and Atmospheric Science)? Given the familiarity with the GWP* metric, particularly in recent discussions of the warming impact of livestock, I think it would add clarity to the reader if the authors referred to GWP* when discussing their CRFD approach, and noted the similarities/differences.

Our aCO2eq is similar in motivation, and shares some aspects of approach, with GWP* but differs in not attempting to provide a direct equivalence to traditional GWPs, which makes our calculation simpler and we believe easier to understand, and in comparing the results of arbitrarily complex interventions to sustained reductions in CO2 emissions rather than pulses. We have added a paragraph noting the relationship and explaining the differences.

9) Use of both ’15 year’ and ’15-year’ in the text (consistency).

Fixed.

10) In the text the authors switch between using ‘%’ and ‘percent’ I’d advise using ‘%’ only.

Fixed.

11) Bottom of page 11, regarding “19% of protein in the human diet” – this needs a reference.

Fixed.

12) Emissions impact comparison to driving: the authors need to be careful making this comparison. In a previous similar comparison, the Long Shadow report was criticized for comparing ‘tailpipe’ transport emissions to emissions from a comprehensive LCA of livestock. A similar comparison is done here, whereby the authors are not considering LCA emissions of driving (car construction, road infrastructure, etc.). I agree it is helpful to make comparisons people are familiar with, and no comparison will be perfect, but the one given here is at risk of being challenged in the same way the Long Shadow report was.

We replaced this calculation with the results of a recent LCA analysis, using values for 2021 sedans in the US. The results are virtually identical, as the increased efficiency of 2021 vehicles compensates for the inclusion of emissions from fuel and vehicle production.

13) Linking back to the major comment 2) in this section, the results shown on page 12 (kg beef -> 470kg CO2eq) are very large, and driven by the biomass recovery estimates: the food

meta-analysis of Poore and Nemecek (2018, Nature) found median beef emissions of 50kg $CO_2$eq, without considering biomass recovery. Have the authors looked for any other similar results to compare to their own? And given the very large influence and uncertainty of the biomass recovery (as discussed already in major comment 2)) it may be worth highlighting the contribution of each of these two components on the 470kg $CO_2$eq and related numbers more clearly.

The 470 kg $CO_2$eq is a combination of effects from emissions and biomass regrowth, which is one reason it is higher than Poore & Nemecek's. That number is also based on a 30 year time horizon, while P&N use 100 year GWPs. Upon further reflection we think it is confusing to use 2100 up to this point and then switch to 2050 (which we did to emphasize short-term impacts of consumer choices), so we have unified all the time horizons in the paper to 2100. Thus reduces the EI of beef to ~300 kg $CO_2$eq/kg product, with ~115 kg / kg coming from emissions reduction. This is very much in line with the estimate of 99.5 obtained from the Supplement of P&N.

14) Regarding the Sala et al., (2021) estimate: my understanding is that Sala et al. estimate annual emissions of 0.58 Gt $CO_2$, lower than the 1 Gt $CO_2$ used in your analysis – please check this. Is this estimated 120 Gt $CO_2$ (which may now need to be revised on the 0.58 Gt $CO_2$ figure), accounting for the additional soy-protein used to replace that food in the food system, as done on the previous animal product estimates?

We have removed this estimate of 120 Gt from the manuscript as it's overly speculative. Sala et al. describe declining yields from 1.47 Gt to 0.58 Gt with successive trawling of the same area, and it's unclear from the manuscript how to accurately model that breakdown. A 0.58 Gt estimate would be conservative, but that's aqueous $CO_2$ and an unknown fraction of that enters the atmosphere, so it's probably safer to not try to compute on this. We have retained the reference and discussion but removed our previous effort to estimate how big an impact this might have on our calculations..

15) Top of page 14: the authors could cite Poore and Nemecek (2018, Nature) as supporting this finding from their own study.

Done.

16) The authors compared their model outputs to those of Riahi et al. (2017) – which outputs were compared?

Clarified.

17) Bottom of page 14: Regarding the economic and social impacts of a transition away from animal agriculture: a reference here would be good. Food security risk for some regions of the world would also be an issue.

Addressed in text.

18) The latest IPBES (IPBES, 2019) report would be a good reference to add to the risk to global biodiversity (in addition to existing Newbold et al. and WWF references).

Done.

19) "Global plant-only diets are feasible without….." – at the macro-level, perhaps, but there would be major changes in some regions, which shouldn't be forgotten.

Addressed in text.

20) Top of page 16: whilst there are significant risk associated with negative emission technologies (NETs), some IPCC scenarios of their deployment are infeasible, and they remain only in early development/operation, this statement needs tempering somewhat: CCS projects are in existence, and are capturing and storing carbon in the order of millions of tonnes (https://www.rff.org/publications/explainers/carbon-capture-and-storage-101/), though this is very far from the scale envisaged in many IPCC scenarios. There are far fewer BECCS projects in operation, and also very few DAC projects: https://www.iea.org/reports/direct-air-capture. These technologies are unproven at large-scale, and we still do not know exactly what their potential delivery of negative emissions/capture carbon is. If the authors want to highlight these risks, and suggest that whilst their scenario is infeasible so are many IPCC scenarios for NETs, they could cite Anderson and Peters (2016, Science) here.

**Methods:**

1) Page 19 ('Estimating species-species land use): no space between 'in' and '*m*' ("except for milk which is reported **in*m***^2).

Fixed

2) Page 24: 'scitik-learn' – should make aware that this is a Python software.

Fixed

3) Page 25: do authors mean '2100' here, not '2200'?

We computed it through 2200 but as we don't report anything beyond 2100 we have changed this.

4) Page 26: references needed for fuel efficiency and emission intensity, although as I have stated above the authors should consider this comparison.

Changed to using LCA as described above. Reference added.

**Tables:**

Table 1: seems to be a repetition of 'chickens' in the table (rows 6 and 7).

The second line should have said "Ducks" and has been corrected.

**Figures:**

1) Figure 2: there are several double spacings in the figure legend (and it looks like in other figure legends too).

Fixed.

2) Figure 6: the 10.6 km per liter of gas should be referenced.

Have updated to describe and cite the LCA we now use.

**Supplementary Figures:**

1) Reference to Figure 2-S1 through 2-S21: For my full understanding of these figures, please could the authors explain why under BAU, atmospheric concentration of CH4 increases decades beyond 2020, when they assume that BAU emissions remain constant at 2019 levels? I would assume the atmospheric concentration to level off sooner, because of the approx. 9-year half-life, but I assume I am misunderstanding something here.

The systems aren't at equilibrium at the beginning because emissions have exceeded decay for years. And even though emissions are constant, it takes longer than one half life to reach equilibrium.

2) Figure 5-S1 through 5-S4: again, for my understanding, the top-left pane shows the CO2 emissions under 'immediate elimination of animal agriculture' return to BAU emissions after 30 years. Please can the authors explain this, given this scenario is based on a plant-based food system which is lower annual emissions than the BAU system? I see that the biomass recovery is modelled over 30 years, but after that period you still have a food system operating on lower emissions than BAU?

Direct CO2 emissions from animal agriculture are only marginally lower than those for a plant based diet, and represent a small fraction of total $CO_2$ emissions. Hence, after carbon recovery finishes, net CO2 emissions - return to being dominated by energy and transportation with only a small decrease due to dietary change. Note that the situation for CH4 and N2O is quite different.

3) Typos in Figure 5-S5: "the decline **int he** first 30 years" and "**decreasess**".

Fixed.

#### Reviewer 2

**Introduction**

- General:
    - The introduction is a bit too "one-sided". It is important to also mention some of the positive aspects of animal production (e.g. contribution to economies, livelihoods, nutrition, soil fertility, …) -- as these would need to be taken into account as "opportunity cost" if livestock production was to stop. This is very much in line with the concept of one of the key references cited in the introduction of the paper, Hayek et al., who also explicitly refer to the "carbon opportunity cost" of using the lands for extensive food production .

We have addressed this briefly in the introduction.

- 
    - A wide range of sectors and processes contribute to global emissions and there is thus not one single solution (or even sector to improve) in order to tackle climate change. This point needs to be clearly stated.

In the introduction we reemphasize that eliminating animal agriculture alone would not solve the climate crisis.

- line 4: I do not find these figures in Hayek et al. and Strassburg et al. Can you specify page/line numbers?

Both Hayek and Strassburg estimate that historical land conversation is responsible for the release of ~800 Gt of CO2. Direct CO2 emissions from fossil fuels are estimated at 1,650 Gt of CO2 (Friedlingstein et al., 2020). Hence ~⅓ of emissions have been due to land conversion.

- p.5, last two lines: The figures mentioned in Hayek et al. and Strassburg et al. are lower.

Hayek, 3rd paragraph: Here we quantify the total carbon opportunity cost of animal agricultural production to be 152.5 (94.2–207.1) gigatons of carbon (GtC) in living plant biomass across all continents and biomes (Fig. 1 and Supplementary Table 3). We approximated the potential for CO2 removal in soil and litter as an additional 63GtC (Supplementary Table 4).

152.5 + 63 = 215.5 Gt C, which is 788 Gt $CO_2$.

Strassburg, see Figure 2b, which reports a maximum capacity of ~900 Gt $CO_2$ recovery. We do not use the Strassburg data directly, rather it serves as an independent confirmation of the value from Hayek.

- p.5, line 4: Can you clarify how the figure of 1,400 Gt was arrived at?

800 Gt from carbon recovery + 80 years * 7.5 Gt / year = 1,400 Gt.

- p.5, line 6: The statement "warming is cumulative" needs a bit of nuancing, as it is only applicable to long-lived climate pollutions (i.e. not to methane).

We have rephrased for clarity. Our intention was to point out that because warming is cumulative, the timing of an increase or decrease in RF due to changes in emissions matters. This is true irrespective of the gas whose levels are being adjusted. For long-lived gases like CO2 the cumulative effect manifests with pulses of positive/negative emissions. For short-lived gases like CH4 it requires sustained changes to see the effect. But in either case the timing of when changes take place matters.

- p.5, line 5-8: Some more detailed explanation underpinning the statement that "this understates impact of dietary change on global warming" would be helpful.

We have rephrased for clarity.

- p.6, last line second last paragraph: "assuming that all other sources of emissions remain constant at 2019 levels" - this seems a fundamentally wrong assumption, as the reduced production of food/nutrition/manure/income as a result of eliminating animal agriculture would need to be compensated for.

We explicitly account for replacement diets in our model, and thus do not hold agricultural emissions constant. This sentence should have read “assuming that all **non-agricultural** emissions sources remain constant at 2019 levels” - we have made this correction.

**Results,**

- General: This section also contains quite a bit of methodology. The manuscript would benefit from having all the details provided here integrated in the methodology section.

We have moved some additional methodological details to the methods where appropriate.

- second paragraph: Please indicate the data source of the total human emissions.

Done.

- p.7 biomass recovery: This explanation is missing in the methods section. Using this figure is also flawed, as Hayek et al calculates the potential sequestration associated with converting land to native/natural state while some of the land will have to be converted to cropland for plant-based food production.

Hayek accounts for land use for a replacement diet in their numbers. This was confirmed with the author.

**Methods,**

- General:
  - It would be helpful to start the methods section with an overview, ideally with schematic.

We drafted schematics to try to capture what we did clearly but were unable to come up with something that we felt enriched the paper, so hope that the changes to the manuscript make it clearer.

- There is a mismatch in the current methodology between steps that are "rough estimates" with steps that are worked out through complicated formulas that seem to imply high precision.  Kindly address.

We use exclusively data from published sources that we would classify not as rough estimates, but as best current measures of the extent of global animal agriculture, emissions due to animal and non-animal agriculture, and land use. We also explicitly stayed away from any complex formula in our analyses and believe the only thing that might be characterized as such is the RF calculation, which is the standard form used in the literature.

- The methodology combines many different steps, all associated with large uncertainties.  I believe the authors would need to address a few related points: (i) each of the separate uncertainties need to be clearly stated (methodology) and quantified (results); (ii) an estimate of the combined uncertainty needs to be included; (iii) this needs to be extensively discussed in the discussion session.

We have added a new section that explicitly addresses major areas of uncertainty and their impact on the results.

- p.18, last paragraph: Can you clarify to what the emission data was scaled?

Done.

- p.20, emissions from agriculture: Please, state explicitly which sector(s)' data you used.

Clarified.

- p.19, diet-linked emissions:
    - Please indicate which of the soybean scenarios from Behnke et al. were used (and the actual figure) for replacing the emissions associated with livestock production with emission associated with soy bean production when modeling reductions in livestock consumption? As the treatments in Behnke et al. are somehow "best practices" with e.g. low fertiliser rates and very localised, I doubt it is realistic to use this as a global GHGe estimate. Please, use a more globally representative data source and confirm that the number used is a realistic global GHGe estimate, by comparing with a number of other sources (across different agro-ecologies, systems, geographies).
    - Can you clarify if a similar replacement (animal product replaced by soy bean) for the land use estimates is also carried out? If not, I believe this is an adjustment that should be made, thereby also keeping in mind that in large areas used for animal feed production (e.g. the arid rangelands), plant-based protein production would require larger areas of land than animal-based protein production.

In response to your comments and those of Reviewer 1 we have switched from using the soy replacement diet, which we viewed as a limiting case, with emissions data from non-animal agriculture from FAOSTAT. This has a nominal impact on the results. In the sensitivity section we also include data  from Xu et al. 2021  for global plant-based diets based on a more comprehensive analysis. This reduced the projected positive impact by around 5%.

- p.20, Emissions projections:

- BFD: write in full

This was a reference to an analysis not in the current version of the MS and has been deleted.

- BAU scenario: fixing emissions at 2019 level is not realistic, as "reductions are likely to be achieved through e.g. increasing agricultural efficiency, reducing food waste, limiting excess consumption, increasing yields, and reducing the emission intensity of livestock production" (as mentioned in the introduction).

We explicitly did not attempt to model any of these factors, which we agree could reduce the impact of animal agriculture in the future. We do not make any claim that emissions from either agriculture or non-agricultural sources will continue at their current rates. Rather we project current rates out to the future as a way to capture the current impact of animal agriculture cognizant of the fact that the benefits of its elimination will accrue over time. We have clarified this motivation in the introduction.

We also note that, while it is likely true that reductions in emissions from animal agriculture can be achieved, current projections are that there will be increases in global consumption of animal products, potentially offsetting increased efficiency. We address both points in the manuscript.

- Can you provide some more detail about the **carbon recovery rate** (magnitude and assumptions; 30 vs 50 years)?

The 30 years is based on assumptions from Hayek, via Griscom but, as estimates vary widely, we have now included 50 and 70 years recovery periods in the sensitivity section.

- Estimating global non-anthropomorphic emissions:
  - Aren't these emissions already taken into account somewhere in the FAOSTAT "Environment_Emissions_by_Sector_E_All_Data_(Normalized)"

No. The categories of emissions in this dataset are “Agriculture total”, “Agricultural land use”, “Energy”, “Industrial processes and product use”, “Waste”, “International Bunkers” and “Other n.e.c.”. The “Other n.e.c.” values for the most recent year are 14 kT $CO_2$, 48 T $CH_4$, 565 kT $N_2O$, which are way too small to account for non-anthropogenic emissions.

- Projections of atmospheric gas levels (p.25):
  - What is the data source of the starting levels?

The values in the original paper were from a database we maintain of historical GHG levels from a variety of sources. For clarity and data integrity we have updated this to single-source data from NOAA.

- Computing emission and land carbon opportunity cost, Factor of 2:
  - as the terrestrial sinks are already included in the calculation of atmospheric C concentration, isn't this double-counting?

It is just reversing the 2 used to go from emissions to atmospheric levels.

Because of terrestrial/oceanic sinks 1 Gt of $CO_2$ emissions only yields an increase of 0.5 Gt atmospheric $CO_2$, thus a decrease of 0.5 Gt of $CO_2$ in the atmosphere is the equivalent of a reduction of 1 Gt of $CO_2$ emissions, hence the factor of 2.

- Computing Carbon Emissions Budgets for RF 2.6 and 1.9:
  - o Please explain why RF 2.6 and 1.9.

In the IPCC's Representative Concentration Pathway framework, 2100 RF values of 2.6 and 1.9 are used, respectively, as surrogates for 2.0C and 1.5C warming.

https://sedac.ciesin.columbia.edu/ddc/ar5_scenario_process/RCPs.html

  - o "RF calculations used in climate models", which climate models are being referred to?

Updated to be clear that we are specifically referring to MAGICC6.

  - o "the RF as calculated above" - which calculation exactly does this refer to (to "the complete RF output of MAGICC6" or to the calculations described in the Radiative Forcing section)?

Clarified in text.

- aCO2eq:
  - o first sentence: How were the $CO_2$ emission equivalents computed?
  - o "simulations described above" - please, specify where exactly is "above", i.e. which simulations are referred to?

Clarified in text.

- Product equivalents
  - o line 25: "per protein" missing.
  - o p.26: Please compare the calculated value of 470kg $CO_2$ eq/kg beef with some values in the literature - e.g. the FAOSTAT data source you used for estimating the overall emissions from agriculture - and explain where the huge difference is coming from.

As noted above in response to a suggestion from Reviewer 1, we updated the calculation as described in the response to use emission reductions through 2100 (it was previously 2050 to highlight the short term impact of eliminating animal agriculture) for consistency with the rest of the manuscript and makes our estimates more directly comparable to those in the literature. This is a more conservative assumption and results in a lower value of 297kg $CO_2$eq/kg beef.

We now compare that directly to the global mean estimate from Poore and Nemecek, and explain that the difference in magnitude comes primarily from our inclusion of carbon fixation on land taken out of agricultural use:

These product-specific aCO2eq's can be interpreted on a per product unit (Figure 6B) or per protein unit (Figure 6C) as emissions intensities. Eliminating the consumption of a kilogram of beef, for example, is equivalent to an emissions reduction of 297 kg CO2. 38 percent (113 kg aCO2eq) comes from reduced emission, in line with the mean estimate of 99.5 kg CO2eq from a systematic meta analysis of GHG emissions from agricultural products (Poore and Nemecek, 2018), with the remaining 62 percent from biomass recovery.

**Discussion,**

- p.15: Apart from calories, protein and fat, it is also worth to say something about micro-nutrients.

All essential micronutrients are readily available at scale from non animal sources. Any reasonably balanced plant based diet can be counted on for everything but Vitamin B12 and sometimes iron. B12 can be produced very inexpensively at scale from microbial sources. Iron requires more attention, but a reasonable plant based diet can cover it.

- treatment of methane: The long-and short-term warming effects of methane and CO2 are very different and there is an ongoing debate as to how to weigh the methane emissions; it would be worth saying something about that in the discussion.

The debate about methane involves, essentially, how to credit effects over different time horizons to emission pulses. By directly modeling methane levels from emissions and decay, and relating them to RF, we avoid this issue.

- perspectives: Please include a short discussion on the social and political feasibility of eliminating animal production completely.

We view this paper as being about the climate potential of eliminating animal agriculture. We expressly avoided offering what would essentially be an opinion about feasibility, as that is more a statement about politics and economics than anything else.

**Figure 1:** The conversion factors for methane and nitrous oxide used here are not standard. Please, adjust.

These are $GWP_{100}$ values used by FAO/GLEAM http://www.fao.org/gleam/model-description/en/ which are taken from the IPCC AR5, Chapter 8, Table 8.7 https://www.ipcc.ch/site/assets/uploads/2018/02/WG1AR5_Chapter08_FINAL.pdf using the values that include climate-carbon feedback.

#### Version #2 Submitted to JOURNAL on October 9, 2021

Version 2 is https://www.biorxiv.org/content/10.1101/2021.04.15.440019v4

#### Decision #2 Received from EDITOR on October 29, 2021

Dear Michael and Pat,

I have sent a response for your article.

Please address the comments in the attached and you will be totally ready.

Best regards,

EDITOR

#### Decision #2 Received from JOURNAL on October 30, 2021

Dear Dr. Eisen,

Thank you again for your submission to JOURNAL.

As relayed to you by email by EDITOR, the reviewers identified some issues with the R1 version of your manuscript that need to be addressed before your paper can be editorially accepted. I understand that you have supplied a revised version of the manuscript incorporating responses to these points by email yesterday, but am issuing a Minor Revision to enable you to submit these new filesManager. Once we have received the revised manuscript in the submission system, we will expedite a final editorial decision.

Thank you very much for your patience and understanding in this matter. We look forward to receiving the revised manuscript as soon as you are able.

All best wishes,

JOURNAL

### Reviewer 1

**Comments**

I commend the authors on the efforts taken to fully address comments from me and the other reviewer. It is good to see them thoroughly engaged with, including the addition of sensitivity testing in Figure 7, and extensively in the supplementary figures, which I think makes the study much more robust and emphasizes that the conclusion of the study withstands under sensitivity testing of the assumptions. As it appears in the new results, the assumption of what plant proteins replace animal products does not have a particularly sizeable impact on the results, which I think is a useful result.

Reviewing again the results, I particularly like the contribution this study makes in Figure 8, alongside the SSPs, where there has been a lack of radical scenarios concerning meat and dairy consumption.

I believe that this study now meets the necessary conditions for publication with JOURNAL, upon addressing a small number of comments on the new version of the manuscript below.

1. Not sure I agree with the removal of "this century" from the title as it details what that 68% refers to.

Restored

2. I think you interchange between "phase out" and "phaseout" – consistency (also consistency of "plant-based" and "plant based", e.g on PDF page 106).

Fixed

3. PDF page 89: change to "often focus on only one factor".

Done

4. A thought on the Hayek et al. (2021) estimates: we should expect a rise in non-livestock ruminants under ecosystem and biodiversity recovery (e.g bison), though that would of course be hard to model and likely to have only a modest reduction in the potential GHG and warming savings of a plant-based diet transition. Had you also thought about this? It could be worth noting in the text.

Have added the following to the sensitivity section: "In some areas, the removal of land from use in animal agriculture may lead to an increase in wild ruminant population. Although this is difficult to model globally, this would offset some of the beneficial impacts of reductions in methane emissions from livestock (Kelliher and Clark, 2010)."

5. Good to see the strong emphasis that this debate typically overlooks ongoing emissions, not the reversible warming impact (biomass recovery).

6. Regarding the use of Twine (2021), on my previous recommendation, I since spoke to the author of that paper and it appears he made an error, understandably given the inconsistent use of GWP weightings used by the FAO from which data was used. See Richard Twine's blog post for more info. I asked him how best to reflect this debate in a paper and he agreed stating the following could be one way forward:
"*estimated at 14.5% by the FAO, although this is based on outdated data and likely now represents an underestimate (Twine, 2021)*"

The Food and Agriculture Organization (FAO) of the United Nations estimates that emissions from animal agriculture represent around 7.1 Gt CO2eq per year (Gerber et al., 2013), 14.5% of annual anthropogenic greenhouse gas emissions, although this is based on outdated data and likely now represents and underestimate (Twine, 2021)

Page 94 of PDF: "decay in the atmosphere on relevant timescales" – doesn't seem worded correctly. They decay on a different timescale to $CO_2$ is the relevant point.

However, a substantial fraction of the emissions impact of animal agriculture comes from methane (CH4) and nitrous oxide (N2O), which decay far more rapidly than CO2 (the half-lives of CH4 and N2O are around 9 and 115 years, respectively), and recent studies have highlighted the need to consider these atmospheric dynamics when assessing their impact (Allen et al., 2018, 2016; Cain et al., 2019)

7. Page 94 of PDF: "How… would alter 2019 net anthropogenic emissions" – I don't think the language quite works here. The impact of phaseout isn't altering emissions in 2019, but you are comparing the impact to emissions in 2019.

Removed 2019 from that sentence.

8. Page 97 of PDF: "Coming from our not including", suggest change to 'our exclusion of'

Done.

9. Page 101 of PDF: I am trying to understand the final sentence here, and find myself confused. PHASE-POD has the same effect, through the end of the century, as a 68% cut in BAU CO2 emissions. This is comparable to eliminating all CO2 by 2050. It seems to me that $CO_2$ elimination by 2050 is a bigger impact than a 68% cut over the century, because after 2050 that $CO_2$ level is zero out to 2100 in the former case. I think this confusion is arising because of comparing emission totals over different time horizons? Also, eliminating all $CO_2$ by 2050 is essentially meeting the Paris Agreement but you state that your results find the phaseout of animal products alone would not be sufficient. Is this because you are only looking at eliminating CO2, not CH4 and N20. It is likely my misunderstanding, but please consider how you can improve clarity to the reader here.

We have removed the sentence about Net Zero, as we agree this was/is confusing.

10. Page 100 of PDF: sub-title refers to offsetting 65% but elsewhere this is written as 68%.

Fixed.

11. Page 102 of PDF: use of “percent” (“62 percent”), consistency needed, with ‘%’ used elsewhere. This also appears elsewhere (e.g “38 percent”, also on PDF page 102).

Fixed.

12. Page 103 of PDF: “2021 model year” – change to ‘2021 model’? ‘year’ seems unnecessary.

Fixed.

13. Much better discussion of the carbon recovery of land.
14. Page 105 of PDF: Change “from Hayek” to “from Hayek et al. (2021)”.

Fixed.

15. Page 106 of PDF: “emissions projected emissions” – typo here?

Fixed.

16. 0.58 to 1.47 – this is per year right? And 1.47 was the initial year of disturbance?

Fixed.

17. Page 109 of PDF: is that “400 million” in the Springmann et al (2018) reference, or does it need referencing separately? It’s a good and supportive statistic, as long as it is referenced.

Added reference.

18. Page 109 of PDF: Good to emphasize the conservative nature of your estimate regarding projected increase in meat consumption. If you wanted to cite it, Searchinger et al (2019) projects a 70% increase in animal-based foods between 2010 and 2050.

Added reference.

19. Page 111 of PDF: “they anticipate”: change to ‘the IPCC’. Later in this paragraph you state “Thus, all currently viable solutions” after having (very reasonably) questioned viability of BECCS at scale. Perhaps rephrase to “Thus, all potential solutions” to avoid contradiction.

Fixed.

20. Page 122 of PDF: “Mt” used and then “MT” later in sentence.

Fixed.

#### Reviewer 2

(Note: This contains the reviewer's original comments in black, our original responses in red, their replies in green, and additional requests in orange, and our final response in purple.)

The topic of the paper, how to tackle the climate emergency and how reduced or eliminated animal agriculture can contribute, is highly relevant.

The singular focus of the paper on animal agriculture as THE climate change solution should be nuanced, as many sectors and processes contribute to GHG emissions. It is not possible to tackle climate change by focusing on one single solution (or even one single sector). In addition, some of the trade-offs and barriers/challenges in relation to eliminating animal agriculture need to be brought more clearly to the readers' attention.

Although the focus of this paper is on animal agriculture, it was not our intention to suggest that it is the only solution. Indeed in both the text and analyses we emphasized that even if the full extent of the potential we highlight here were realized, it would still not be enough to solve the climate crisis. We have made this point clearer at multiple points in the manuscript.

This is highly appreciated

We also explore the challenges to eliminating animal agriculture in more detail.

Thanks

Most importantly, however, the results as presented (e.g. offset of 70% possible by eliminating animal agriculture) that could justify this strong focus, need to be corroborated and presented with much more nuance. The methodology followed in the manuscript combines many different calculations based on many assumptions and with large separate uncertainties associated with it.

As discussed above, we have added a section exploring the sensitivity of our results to assumptions made in our analyses so that readers can get a better sense of the degree of uncertainty in our calculations and its sources.

I believe the section exploring the sensitivities to the assumptions adds value to the paper

More detailed comments (about methodology as well as the other parts of the manuscript) are attached.

We have responded below to the reviewer's comments and modified the manuscript where appropriate.

**Introduction**

- General:
  - The introduction is a bit too "one-sided". It is important to also mention some of the positive aspects of animal production (e.g. contribution to economies, livelihoods, nutrition, soil fertility, …) -- as these would need to be taken into account as "opportunity cost" if livestock production was to stop. This is very much in line with the concept of one of the key references cited in the introduction of the paper, Hayek et al., who also explicitly refer to the "carbon opportunity cost" of using the lands for extensive food production .
  -

We have addressed this briefly in the introduction.

Thanks

- 
  - A wide range of sectors and processes contribute to global emissions and there is thus not one single solution (or even sector to improve) in order to tackle climate change. This point needs to be clearly stated.

In the introduction we reemphasize that eliminating animal agriculture alone would not solve the climate crisis.

Thanks

- line 4: I do not find these figures in Hayek et al. and Strassburg et al. Can you specify page/line numbers?

Both Hayek and Strassburg estimate that historical land conversation is responsible for the release of ~800 Gt of $CO_2$. Direct $CO_2$ emissions from fossil fuels are estimated at 1,650 Gt of $CO_2$ (Friedlingstein et al., 2020). Hence ~⅓ of emissions have been due to land conversion.

I do not seem to find the estimate of historical land conversion in Hayek et al., but maybe I am overseeing it somewhere.

I think our use of the word historical is confusing. All we mean is that it happened in the past.

Line 4 in the manuscript refers specifically to "The historical reduction in terrestrial biomass as native ecosystems were transformed to support grazing livestock and the cultivation of feed and forage crops" while the Strassburg et al. paper explicitly considers all lands converted from natural ecosystems, including those that were converted to croplands (which in my reading of their paper would also include food crops). So, I'd suggest to edit the sentence accordingly

We will remove the Strassburg reference here as we do not use the data explicitly in our calculations.

- p.5, last two lines: The figures mentioned in Hayek et al. and Strassburg et al. are lower.

Hayek, 3rd paragraph: Here we quantify the total carbon opportunity cost of animal agricultural production to be 152.5 (94.2–207.1) gigatons of carbon (GtC) in living plant biomass across all continents and biomes (Fig. 1 and Supplementary Table 3). We approximated the potential for CO2 removal in soil and litter as an additional 63GtC (Supplementary Table 4).

152.5 + 63 = 215.5 Gt C, which is 788 Gt $CO_2$.

Thanks for pointing this out.  I also appreciate the addition of "In the order of"…

Strassburg, see Figure 2b, which reports a maximum capacity of ~900 Gt $CO_2$ recovery. We do not use the Strassburg data directly, rather it serves as an independent confirmation of the value from Hayek.

As already noted above, Strassburg et al. explicitly consider all lands converted from natural ecosystems, including those that were converted to croplands. So, if you want to keep the last part of the sentence (i.e. "currentLY devoted to livestock production"), I'd suggest you do not cite Strassburg et al. here.

We have removed it.

- p.5, line 4: Can you clarify how the figure of 1,400 Gt was arrived at?

800 Gt from carbon recovery + 80 years * 7.5 Gt / year  = 1,400 Gt.
Thanks for the clarification.

- p.5, line 6: The statement "warming is cumulative" needs a bit of nuancing, as it is only applicable to long-lived climate pollutions (i.e. not to methane).

We have rephrased for clarity. Our intention was to point out that because warming is cumulative, the timing of an increase or decrease in RF due to changes in emissions matters. This is true irrespective of the gas whose levels are being adjusted. For long-lived gases like CO2 the cumulative effect manifests with pulses of positive/negative emissions. For short-lived gases like CH4 it requires sustained changes to see the effect. But in either case the timing of when changes take place matters.

- p.5, line 5-8: Some more detailed explanation underpinning the statement that "this understates impact of dietary change on global warming"  would be helpful.

We have rephrased for clarity.

Thanks for rephrasing and for adding a few words explicitly explaining the short-lived nature of methane.
A few suggestions:

- "…which…decay in the atmosphere on relevant timescales, …": the term "relevant timescales" is not very clear.  Did you mean "relative short timescales"?  If so, I'd have my reservations stating "relative short timescales" in relation to $N_2O$ and would consider removing it.  But maybe you mean ""…decay in the atmosphere within the timescale (or time window) covered by our analysis"? Please, edit accordingly.

We have edited to say:

However, a substantial fraction of the emissions impact of animal agriculture comes from methane ($CH_4$) and nitrous oxide ($N_2O$), which decay far more rapidly than $CO_2$ (the half-lives of $CH_4$ and $N_2O$ are around 9 and 115 years, respectively).

- I feel that the last part of the last sentence "having a greater cumulative effect on warming" is confusing in relation to short-lived GHGs; as it is the long-lived ones that have a cumulative effect on warming. I would thus suggest you edit the sentence as follows: "Of critical importance, many of the beneficial effects on greenhouse gas levels of eliminating livestock would accrue rapidly, via biomass recovery and decay of short-lived atmospheric methane $CH_4$. ~~and therefore~~ Their cooling influence would be felt for an extended period of time~~, having a greater cumulative effect on warming~~."

Done.

- p.6, last line second last paragraph: "assuming that all other sources of emissions remain constant at 2019 levels" - this seems a fundamentally wrong assumption, as the reduced production of food/nutrition/manure/income as a result of eliminating animal agriculture would need to be compensated for.

We explicitly account for replacement diets in our model, and thus do not hold agricultural emissions constant. This sentence should have read "assuming that all **non-agricultural** emissions sources remain constant at 2019 levels" - we have made this correction.
Thanks

**Results,**

Additional comments:

- “The dietary scenarios include the immediate replacement of all animal agriculture with a plant-only diet (IMM-POD), a more realistic gradual transition, over a period of 15 years” – consider editing as follows: “The dietary scenarios include the immediate replacement of all animal agriculture with a plant-only diet (IMM-POD), and a more ~~realistic~~ gradual transition, over a period of 15 years”

Done.

- “We updated estimates of global emissions from animal agriculture by scaling country-, species- and product-specific emission intensities… with country-specific data on primary production of livestock products”: I do not think the word “scaling” is the most appropriate here; maybe just use “multiplying”?

“We updated estimates of global emissions from animal agriculture using country-, species- and product-specific emission intensities from the Global Livestock Environmental Assessment Model (MacLeod et al., 2018), and country-specific data on primary production of livestock products from the Food and Agriculture Organization (FAO) database FAOSTAT (FAO, 2021).”

- General: This section also contains quite a bit of methodology. The manuscript would benefit from having all the details provided here integrated in the methodology section.

We have moved some additional methodological details to the methods where appropriate.

- second paragraph: Please indicate the data source of the total human emissions.

Done.

Thanks

- p.7 biomass recovery: This explanation is missing in the methods section. Using this figure is also flawed, as Hayek et al calculates the potential sequestration associated with converting land to native/natural state while some of the land will have to be converted to cropland for plant-based food production.

Hayek accounts for land use for a replacement diet in their numbers. This was confirmed with the author.

This is indeed the case; thanks for pointing out and confirming.

I would still like to suggest to remove the reference to Strassburg et al. as they do not present results for land currently used in livestock production separately from all agricultural land.

Comment to the new section:

- "…recent modeling work by (Strassburg et al., 2020) that half of the biomass recovery potential of land currently used for animal agriculture could be realized by restoration of 25% of the relevant land". I'd suggest to edit as follows: "…recent modeling work by (Strassburg et al., 2020) indicates that half of the biomass recovery potential of land currently used for ~~animal~~ agriculture could be realized by restoration of 25% of the relevant land"

Done.

**Methods**

- General:
  - It would be helpful to start the methods section with an overview, ideally with schematic.

We drafted schematics to try to capture what we did clearly but were unable to come up with something that we felt enriched the paper, so hope that the changes to the manuscript make it clearer.

Accepted

  - There is a mismatch in the current methodology between steps that are "rough estimates" with steps that are worked out through complicated formulas that seem to imply high precision. Kindly address.

We use exclusively data from published sources that we would classify not as rough estimates, but as best current measures of the extent of global animal agriculture, emissions due to animal and non-animal agriculture, and land use. We also explicitly stayed away from any complex formula in our analyses and believe the only thing that might be characterized as such is the RF calculation, which is the standard form used in the literature.

I acknowledge that the term "rough estimates" might not have been appropriate. The data used does, however, come with large uncertainties. Which might lead to a large variation in terms of e.g. resulting RF values associated with the min-max range of these estimates. The new section on sensitivity deals with this in an acceptable way.

  - The methodology combines many different steps, all associated with large uncertainties. I believe the authors would need to address a few related points: (i) each of the separate uncertainties need to be clearly stated (methodology) and quantified (results); (ii) an estimate of the combined uncertainty needs to be included; (iii) this needs to be extensively discussed in the discussion session.

We have added a new section that explicitly addresses major areas of uncertainty and their impact on the results.

The inclusion of the sensitivity analysis associated with the assumptions (replacement of animal-sourced food & duration of biomass recovery) is highly appreciated.

- p.18, last paragraph: Can you clarify to what the emission data was scaled?

Done.

OK

- p.20, emissions from agriculture: Please, state explicitly which sector(s)' data you used.

Clarified.

- p.19, diet-linked emissions:
    - Please indicate which of the soybean scenarios from Behnke et al. were used (and the actual figure) for replacing the emissions associated with livestock production with emission associated with soy bean production when modeling reductions in livestock consumption? As the treatments in Behnke et al. are somehow "best practices" with e.g. low fertiliser rates and very localised, I doubt it is realistic to use this as a global GHGe estimate. Please, use a more globally representative data source and confirm that the number used is a realistic global GHGe estimate, by comparing with a number of other sources (across different agro-ecologies, systems, geographies).

    - Can you clarify if a similar replacement (animal product replaced by soy bean) for the land use estimates is also carried out? If not, I believe this is an adjustment that should be made, thereby also keeping in mind that in large areas used for animal feed production (e.g. the arid rangelands), plant-based protein production would require larger areas of land than animal-based protein production.

In response to your comments and those of Reviewer 1 we have switched from using the soy replacement diet, which we viewed as a limiting case, with emissions data from non-animal agriculture from FAOSTAT. This has a nominal impact on the results. In the sensitivity section we also include data from Xu et al. 2021 for global plant-based diets based on a more comprehensive analysis. This reduced the projected positive impact by around 5%.

I believe this is an improvement.

New comment to the "replacement diets" section

- "…scaling non-livestock agricultural emission intensities for unit protein by protein required to match that provided by the livestock being replaced" This sentence doesn't flow

When modeling reductions in livestock consumption, we assumed protein from livestock products would be replaced with equivalent amount of protein from current food crops, and used per unit protein emission intensities computed from FAOSTAT to infer emissions from this replacement diet.

- p.20, Emissions projections:
  - BFD: write in full

This was a reference to an analysis not in the current version of the MS and has been deleted.

Thanks

- BAU scenario: fixing emissions at 2019 level is not realistic, as "reductions are likely to be achieved through e.g. increasing agricultural efficiency, reducing food waste, limiting excess consumption, increasing yields, and reducing the emission intensity of livestock production" (as mentioned in the introduction).

We explicitly did not attempt to model any of these factors, which we agree could reduce the impact of animal agriculture in the future. We do not make any claim that emissions from either agriculture or non-agricultural sources will continue at their current rates. Rather we project current rates out to the future as a way to capture the current impact of animal agriculture cognizant of the fact that the benefits of its elimination will accrue over time. We have clarified this motivation in the introduction.

We also note that, while it is likely true that reductions in emissions from animal agriculture can be achieved, current projections are that there will be increases in global consumption of animal products, potentially offsetting increased efficiency. We address both points in the manuscript.

Accepted

- Can you provide some more detail about the **carbon recovery rate** (magnitude and assumptions; 30 vs 50 years)?

The 30 years is based on assumptions from Hayek, via Griscom but, as estimates vary widely, we have now included 50 and 70 years recovery periods in the sensitivity section.

The recovery period is now clear.

It can be deducted from the sensitivity analysis section (in results) that the magnitude of the carbon recovery rate is extracted from Hayek et al. It would be good to mention that in the methods section too.

Done.

- Estimating global non-anthropomorphic emissions:
    - Aren't these emissions already taken into account somewhere in the FAOSTAT "Environment_Emissions_by_Sector_E_All_Data_(Normalized)"

No. The categories of emissions in this dataset are “Agriculture total”, “Agricultural land use”, “Energy”, “Industrial processes and product use”, “Waste”, “International Bunkers” and “Other n.e.c.”. The “Other n.e.c.” values for the most recent year are 14 kT $CO_2$, 48 T $CH_4$, 565 kT $N_2O$, which are way too small to account for non-anthropogenic emissions.

OK

- Projections of atmospheric gas levels (p.25):
    - What is the data source of the starting levels?

The values in the original paper were from a database we maintain of historical GHG levels from a variety of sources. For clarity and data integrity we have updated this to single-source data from NOAA.
Clear, thanks

- Computing emission and land carbon opportunity cost, Factor of 2:
    - as the terrestrial sinks are already included in the calculation of atmospheric C concentration, isn't this double-counting?

It is just reversing the 2 used to go from emissions to atmospheric levels.

Because of terrestrial/oceanic sinks 1 Gt of CO2 emissions only yields an increase of 0.5 Gt atmospheric CO2, thus a decrease of 0.5 Gt of CO2 in the atmosphere is the equivalent of a reduction of 1 Gt of CO2 emissions, hence the factor of 2.
Thanks for the clarification

- Computing Carbon Emissions Budgets for RF 2.6 and 1.9:
    - Please explain why RF 2.6 and 1.9.

In the IPCC's Representative Concentration Pathway framework, 2100 RF values of 2.6 and 1.9 are used, respectively, as surrogates for 2.0C and 1.5C warming.

This is quite clear. I believe it would be worth explicitly referring to these warming targets in the manuscript.

https://sedac.ciesin.columbia.edu/ddc/ar5_scenario_process/RCPs.html

- 
    - "RF calculations used in climate models", which climate models are being referred to?

Updated to be clear that we are specifically referring to MAGICC6.

Thanks

- 
    - "the RF as calculated above" - which calculation exactly does this refer to (to "the complete RF output of MAGICC6" or to the calculations described in the Radiative Forcing section)?

Clarified in text.

Thanks

- aCO2eq:
    - first sentence: How were the CO2 emission equivalents computed?
    - "simulations described above" - please, specify where exactly is "above", i.e. which simulations are referred to?

Clarified in text.
Ok

- Product equivalents
    - line 25: "per protein" missing.
    - p.26: Please compare the calculated value of 470kg CO2 eq/kg beef with some values in the literature - e.g. the FAOSTAT data source you used for estimating the overall emissions from agriculture - and explain where the huge difference is coming from.

As noted above in response to a suggestion from Reviewer 1, we updated the calculation as described in the response to use emission reductions through 2100 (it was previously 2050 to highlight the short term impact of eliminating animal agriculture) for consistency with the rest of the manuscript and makes our estimates more directly comparable to those in the literature. This is a more conservative assumption and results in a lower value of 297kg CO2eq/kg beef.

We now compare that directly to the global mean estimate from Poore and Nemecek, and explain that the difference in magnitude comes primarily from our inclusion of carbon fixation on land taken out of agricultural use:

These product-specific aCO2eq's can be interpreted on a per product unit (Figure 6B) or per protein unit (Figure 6C) as emissions intensities. Eliminating the consumption of a kilogram of beef, for example, is equivalent to an emissions reduction of 297 kg CO2. 38 percent (113 kg aCO2eq) comes from reduced emission, in line with the mean estimate of 99.5 kg CO2eq from a systematic meta analysis of GHG emissions from agricultural products (Poore and Nemecek, 2018), with the remaining 62 percent from biomass recovery.
OK; I can see that you have these new numbers in the results section
However, in the methods section, it still reads 470kg.

Apologies. Fixed.

**Discussion,**

- p.15: Apart from calories, protein and fat, it is also worth to say something about micro-nutrients.

All essential micronutrients are readily available at scale from non animal sources. Any reasonably balanced plant based diet can be counted on for everything but Vitamin B12 and sometimes iron. B12 can be produced very inexpensively at scale from microbial sources. Iron requires more attention, but a reasonable plant based diet can cover it.

This does not negate the fact that e.g. B12 deficiency is already prevalent in several developing countries where a considerable fraction of the population will not be in a position to source supplements or access a sufficiently diverse and healthy plant-based diet to compensate for the micro-nutrients they now get from low levels of consumption of animal products. I understand you do not want to discuss this in lots of detail, but would suggest that you explicitly mention nutrition security in your section on the potential economic and social impacts.

Have added a comment to this effect.

May I also suggest that you edit the last part of the last sentence in that section along the following lines "And, while it is expected that …, investment will also be required to prevent local food and nutrition insecurity in regions where wide-scale access to a diverse and healthy plant-based diet is lacking" - "heavily reliant on animal based foods" may be interpreted as having large intakes of animal sourced food.

Done.

I do appreciate very much that that section is there, by the way. Global modeling efforts, like yours, have great value in outlining the theoretical potential of drastic interventions such as a rapid global phase-out of animal production, but it is also important to note that practical on-the-ground implementation of such strategies needs contextualization and that trade-offs with social/economic/health impacts do need to be taken into account.

- treatment of methane: The long-and short-term warming effects of methane and CO2 are very different and there is an ongoing debate as to how to weigh the methane emissions; it would be worth saying something about that in the discussion.

The debate about methane involves, essentially, how to credit effects over different time horizons to emission pulses. By directly modeling methane levels from emissions and decay, and relating them to RF, we avoid this issue.
OK

New comment:
- ✔ Can you provide a reference for "an additional 4635 million km2 - … - would be needed to support the required growth in livestock populations"?

Adjusted to clarify that this is our calculation and explained: "For example, using land use data from (Poore and Nemecek 2018) and consumption data from FAOSTAT, extending the current diet of wealthy industrialized countries (OECD) to the current global population would require an additional 35 million $km^2$ to support livestock production - an area roughly equal to the combined area of Africa and Australia."

- perspectives: Please include a short discussion on the social and political feasibility of eliminating animal production completely.

We view this paper as being about the climate potential of eliminating animal agriculture. We expressly avoided offering what would essentially be an opinion about feasibility, as that is more a statement about politics and economics than anything else.

Accepted

**Figure 1:** The conversion factors for methane and nitrous oxide used here are not standard. Please, adjust.

These are $GWP_{100}$ values used by FAO/GLEAM http://www.fao.org/gleam/model-description/en/ which are taken from the IPCC AR5, Chapter 8, Table 8.7 https://www.ipcc.ch/site/assets/uploads/2018/02/WG1AR5_Chapter08_FINAL.pdf using the values that include climate-carbon feedback.

OK
